# GLUT4 translocation with insulin: revisiting the case for dose-dependent quantal release

**DOI:** 10.1101/2025.04.11.648479

**Authors:** Irina Romenskaia, Cynthia Corley Mastick, Adelle C.F. Coster

## Abstract

In mammalian fat and muscle cells, insulin stimulates the translocation of the glucose transporter GLUT4 from intracellular storage compartments to the plasma membrane in adipocytes and muscle cells, significantly increasing glucose uptake. In unstimulated (basal) cells, GLUT4 is sequestered in non-cycling/very slowly cycling compartments. Insulin mobilizes GLUT4 by releasing it from sequestration, enabling its exocytosis and continuous cycling between the plasma membrane and endosomes. Upon insulin withdrawal, GLUT4 is rapidly internalized and re-sequestered internally for future activation. Dynamic studies using tagged GLUT4 have revealed that this trafficking mechanism is regulated post-translationally, with GLUT4 undergoing repeated cycles of mobilization and sequestration in response to fluctuating insulin levels.

The trafficking of GLUT4 under basal and maximal insulin concentrations can be modeled as a single cycling pool, with different amounts of GLUT4 in the actively cycling pool. In this model, insulin regulates both the rate constant of exocytosis, *k*_*ex*_, and the distribution of GLUT4 between the cycling and a non-cycling pool. Here, we present modeling of the kinetics of GLUT4 trafficking in 3T3-L1 adipocytes over a range of insulin concentrations, under steady state, and in transition after adding insulin or after adding an inhibitor of exocytosis. Given the observed characteristics of the experimental data, parsimonious explanatory models incorporating different hypotheses of the insulin-dependence of the GLUT4 recycling system are optimized to the data sets to identify dominant processes acting in the dynamics. The steady-state data is best fit with a model that includes a dose-dependent increase in the size of the cycling pool at submaximal insulin concentrations (quantal release). Simultaneous fits of the transition and steady-state data indicate that insulin regulates a second kinetics rate constant, in addition to increasing *k*_*ex*_ and the cycling pool size.

**Summary:** Experimental and modeling investigations of the trafficking of GLUT4 in 3T3-L1 adipocytes at different insulin doses shows the dominant effects of the insulin dose on the dynamics. The data is best fit, particularly at submaximal insulin concentrations, by a model in which both the exocytosis rate and the size of the cycling pool increases with insulin.

## Introduction

The protein GLUT4 is the main insulin-facilitated glucose transporter in mammalian fat and muscle cells. It is a passive transporter and so allows glucose to pass according to the driving concentration gradient. As the protein is membrane-embedded and does not exist freely in the cells, glucose transport through this channel is regulated by translocation of the protein itself. Thus, it has been observed that the expression of GLUT4 at the plasma membrane in the absence of insulin is very low, approximately 20% of the cell total, whereas in very high insulin, >100nM, is of the order of 70% of the cell total (1,2).

There have been numerous studies on the action of insulin in adipocytes and muscle cells, both from the perspective of the biochemical signalling pathway and the translocation of GLUT4 (see for instance (3,4) for reviews). Most of the work was, however, conducted on cells either in basal (0nM) insulin or at maximal insulin concentrations – at least an order of magnitude higher than physiological levels (usually 100nM or more). Some exceptions include (1,2) which measured the steady state response and the translocation from the basal state of GLUT4 in 3T3-L1 adipocytes in the presence of 0, 0.24, 3.2, and 100nM insulin and (5) which tracked the differential response of the surface of levels of GLUT4 as increasing amounts of insulin (0, 0.1, 1, and 100nM) were sequentially applied.

Here we characterize the insulin-dose dynamics of GLUT4 in 3T3-L1 adipocytes over a range of insulin doses under steady state with respect to insulin, and the transitory behavior when insulin is first applied. The dynamics under inhibition of exocytosis via the PI3K inhibitor LY294002 is also characterized.

Parsimonious explanatory modeling of this data is used to explore the characteristics of the dynamics to investigate some of the major processes that could underlie the observed behavior and to test possible mechanisms.

## Experimental Procedures

### Tissue Culture, Viral Infections, and Antibodies and Reagents

3T3-L1 cells were sourced from ATCC and cultured as fibroblasts in DMEM complete medium, consisting of high-glucose DMEM supplemented with 10% calf serum, 2 mM L-glutamine, 50 units/mL penicillin, and 50μg/mL streptomycin. Fibroblasts were plated at near-confluence (achieving full confluence within 24 hours) and subsequently refed with complete medium containing 10% calf serum. Experiments were conducted 1-2 days post-confluence. For experimental purposes, cells were differentiated into adipocytes as previously described (6).

The lentiviral HA-GLUT4/GFP reporter protein was prepared and transduced into fibroblasts as previously described (6). Cells were infected at a viral titer that resulted in approximately 50% of cells expressing the construct. At this titer, most infected cells contained only a single virion, with no detectable cytopathic effects (6,7). Uninfected cells were used as internal controls to account for cellular autofluorescence and nonspecific antibody binding or uptake. This reporter has been thoroughly characterized; when expressed at the low levels observed in our infected cells, the HA-GLUT4/GFP reporter exhibits trafficking behavior consistent with endogenous GLUT4 (6,8-10).

The HA.11 monoclonal antibody (α-HA; Covance) was purchased as ascites and purified using a 1-ml rProteinA-FF column (GE Healthcare) following established protocols (6). The purified antibody was labeled with AlexaFluor647 (AF647) using a protein labeling kit (Invitrogen), following the manufacturer’s instructions. Excess dye was removed by performing two rounds of desalting into PBS using 10-ml Zeba spin columns (7000 molecular weight cutoff, ThermoScientific) (11). Protein concentrations and labeling efficiency were measured via absorption spectroscopy (6).

### Flow Cytometry

Flow cytometry, gating, and analysis were performed as previously described (6,7,11,12). Labeled cells in 96-well plates were kept on ice and washed three times with 200 μL of ice-cold PBS. Adipocytes were incubated with 20 μL of collagenase (Type III, 1 mg/mL in PBS with 2% BSA; Worthington) at 37°C or 4°C for 10 minutes, then resuspended in 180 μL of PBS. The cell suspensions were gently filtered through a 100-μm cell strainer to remove clumps and analyzed using an Accuri C6 cytometer. Detection thresholds for adipocytes were set at 1,000,000 for forward scatter height (FSC-H) and 500,000 for side scatter height (SSC-H).

Log intensities of forward scatter, side scatter, and fluorescence signals (FL1: 488 nm excitation/533 nm emission; FL2: 488 nm excitation/585 nm emission; FL3: 488 nm excitation/>670 nm emission; FL4: 640 nm excitation/675 nm emission) were collected for each cell. Adipocytes were selectively gated using CFlow Plus software (Accuri) to exclude residual fibroblasts and cellular debris, as described previously (6). Cells infected with HA-GLUT4/GFP were distinguished from uninfected cells using a two-dimensional histogram of FL1-H (GFP) versus FL3-H (autofluorescence).

The geometric mean fluorescence of gated populations was calculated using FCS Express (De Novo Software). Mean fluorescence values of uninfected cells, representing cellular autofluorescence (FL1) and nonspecific antibody labeling (FL4), were subtracted from the corresponding mean values for infected cells in the same sample before calculating the mean fluorescence ratio (MFR) values (7).

### GLUT4 Transition and Inhibition Assays/Surface GLUT4 Labeling

Cells expressing HA-GLUT4/GFP were incubated for 2 hours in low serum medium (LSM), consisting of DMEM complete medium with 0.5% fetal bovine serum. For the basal-to-insulin transition, varying doses of insulin were added to the cells for increasing durations. For inhibition experiments, which involved transitioning from the insulin-stimulated state to the phosphatidylinositol 3-kinase (PI3K)-inhibited state, insulin was applied during the final 45 minutes of preincubation in LSM, followed by the addition of the PI3K inhibitor LY294002 (LYi, 50 μM; EMD Biochemicals) for increasing durations. Surface GLUT4 was labeled by rapidly chilling the cells on an ice-water slurry and incubating them with LSM containing 50 μg/mL AF647-α-HA for 1 hour on ice. After antibody removal, the cells were washed, digested with collagenase, and analyzed by flow cytometry as described previously (7,11). The data is normalized in these assays to the average level at 60 minutes in 100nM insulin.

### Steady-state Anti-HA Antibody (α-HA) Uptake Assay

Cells were serum-starved in LSM as previously described. For insulin-stimulated uptake, some cells were treated with varying doses of insulin during the final 45 minutes of starvation. At specific time points, the medium was replaced with 30 μL of warm LSM containing 50μg/mL AF647-α-HA, with or without insulin, and incubation continued. At the end of the time course, cells were chilled, washed, digested with collagenase, and analyzed by flow cytometry (7,11). Under these assay conditions, surface HA-GLUT4 is labeled nearly instantaneously, and the antibody remains bound throughout the procedure. The data is normalized in this assay to the average level at 300 minutes in 100nM insulin.

### Data reporting and Normalisation

To account for variability in reporter construct expression across cell populations, mean fluorescence ratio (MFR) values were calculated for each sample (mean fluorescence of AF647 divided by mean fluorescence of GFP). The α-HA data are presented normalized to the total GLUT4 that can be labeled in 100 nM insulin-stimulated cells in each assay, (MFR/MFR_max_), Labeled GLUT4/Max.

### Computational Methods

Optimized least-squares fits of the models were undertaken using the trust region algorithm ‘fit’ function in Matlab (Matlab R2024a, Mathworks 2024). The options were left at the default values. For the models described by coupled differential equations, these were simulated using the Matlab explicit Runge-Kutta (4,5) ODE45 function using the default settings. The outputs were then coupled to the ‘fit’ function using custom scripts.

### Replicates

The range of insulin concentrations, sample times and number of replicates for each of the Uptake, Transition, and Inhibition assays are shown in Table 1.

**Table 1.**
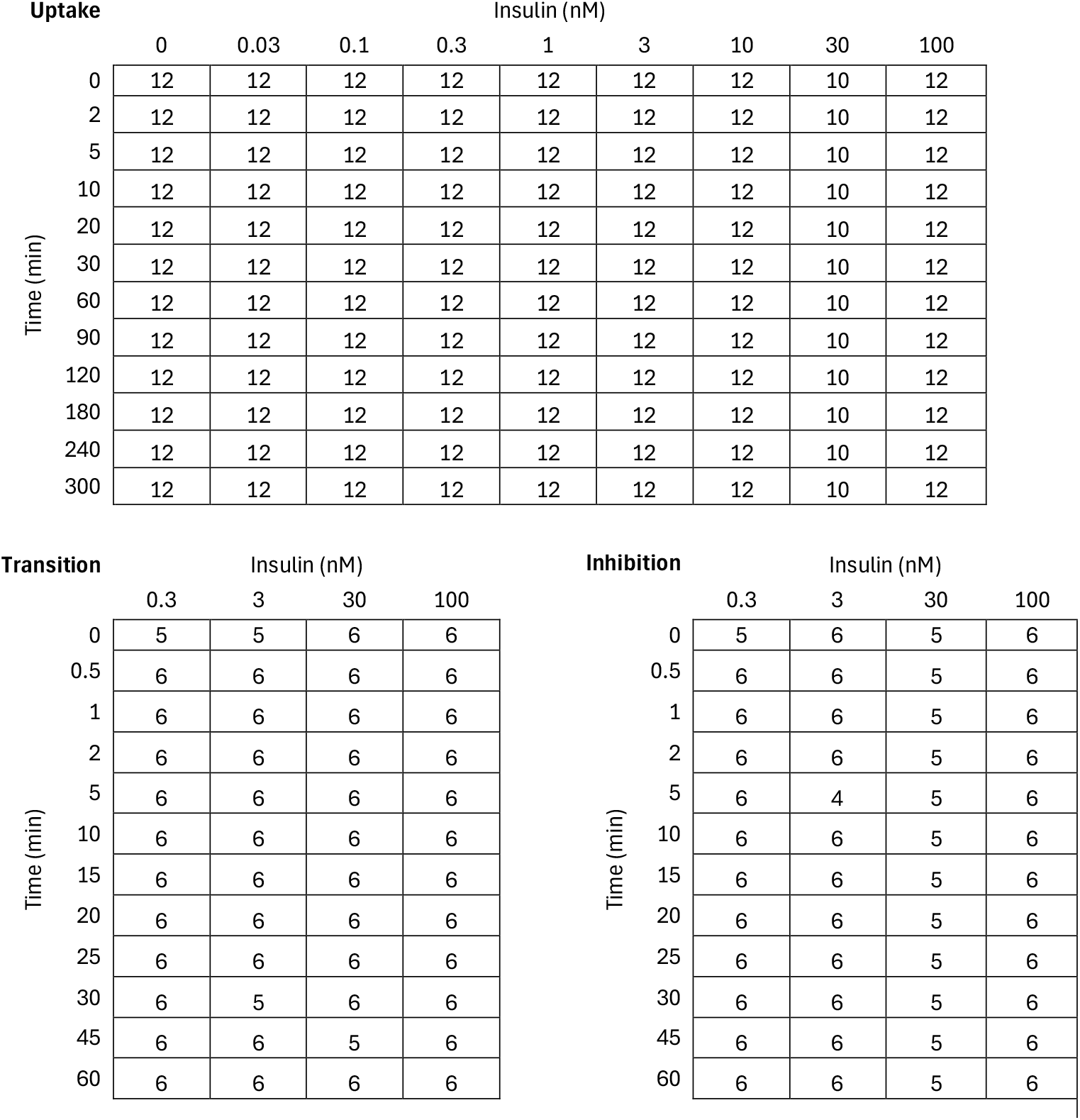
The number of replicates for each of the experimental assays as a function of the insulin concentrations and sample times.

## Characteristics of the Experimental Assays

### Steady State Experiments

The Steady-state Anti-HA Antibody (α-HA) Uptake Assay tracks the cumulative amount of GLUT4 molecules that have transited to the plasma membrane in the presence of a given level of insulin. It is an indicator of the overall exocytosis rate in a steady state of insulin. It is assumed that the unlabeled molecules are “instantly” labeled (on the time scale of the experiment) as they are brought to the plasma membrane by exocytosis and this labelling is assumed to be irreversible.

The assay was performed with different levels of insulin applied to the cells between 0 and 100nM. The results of the assay, Figure 1, which reports the total amount of GLUT4 that has recycled to the surface of the cell during the given time, are normalized to the mean reported level in the presence of 100nM insulin at 300 minutes.

**Figure 1.**
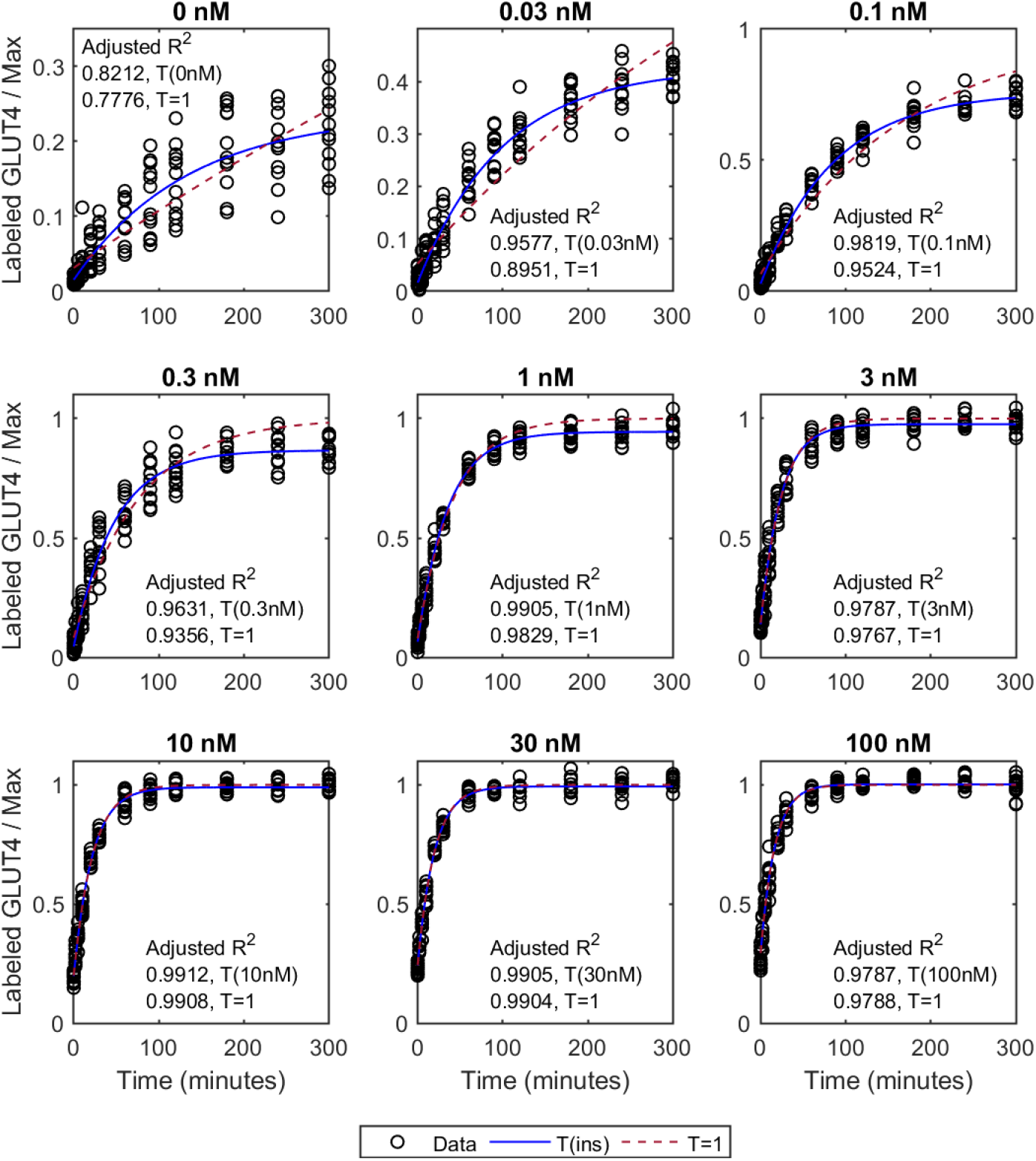
Least-squares fits for the Uptake Assay to the T(ins) and T=1 models as a function of the time of the data for each insulin concentration (indicated above each subplot). The Adjusted R^2^ for each fit is indicated inset on the plots. Note that the two models are practically identical for 100nM in line with the normalization of the data. Here the fits to Equation (2) are presented, but identical results were achieved using Equation (1).

It can be seen in Figure 1 that the Uptake Assay data at any given level of insulin is well described by single time constant for all exocytosis, i.e., the data followed the trend of an exponential rise to a plateau, i.e.,

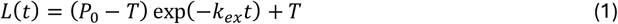

where *L* is the level of labeled GLUT4 as a function of time, *t* ; *P*_0_ is the initial level of labeled GLUT4 (that initially at the plasma membrane); *k*_*ex*_ is the rate constant for exocytosis; and *T* is the plateau level corresponding to the amount of GLUT4 recycling in the system.

A recycling system with a single time constant can be described as a parsimonious linear system to describe the location of GLUT4 in the cell in one of two compartments – the plasma membrane, *P*, and internal structures, *E*. This system has one overall exocytosis rate for all exocytosis pathways, *k*_*ex*_, and one endocytosis rate, *k*_*en*_. These rates represent the average rate of all processes acting to bring GLUT4 to the surface and re-internalize it respectively.

Conservation of GLUT4 over the time scale of the experiments indicates that *P* (*t*) + *E* (*t*) = *T*, for all times, *t*, and 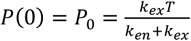, giving an alternative version of the output of the uptake assay as,

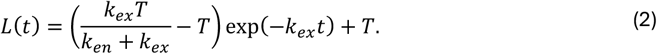

Note both descriptions of the Uptake Assay have the same number of parameters and *k*_*ex*_ and *T* have the same definition in Equations (1) and (2).

There has been debate in the literature regarding whether the total amount of GLUT4 that recycles to the plasma membrane is insulin dependent. Here we consider two models to investigate this using the dose response data results of the uptake assay.

Model T(ins) is described by Equations (1) and (2) in which the plateau level, *T*, is assumed to be insulin concentration dependent. Recalling that the data is normalized to the mean GLUT4 level at 300 min with 100nM insulin, Model T=1 assumes the plateau level, *T* = 1 in Equations (1) and (2), i.e., all GLUT4 is recycling irrespective of the insulin concentration.

Models T=1 and T(ins) were fitted to the data using least-squares optimization, Figure 1 and Table 2. The fits using Equation (1) produce practically identical results to those using Equation (2) for Models T(ins) and T=1 – they simply either estimated *P*_0_ or *k*_*en*_. It is useful to quantify the *k*_*en*_ values, however, as this allows us to assess whether these vary as a function of the insulin dose. The inferred parameter values are compared in Figure 2.

**Table 2.**
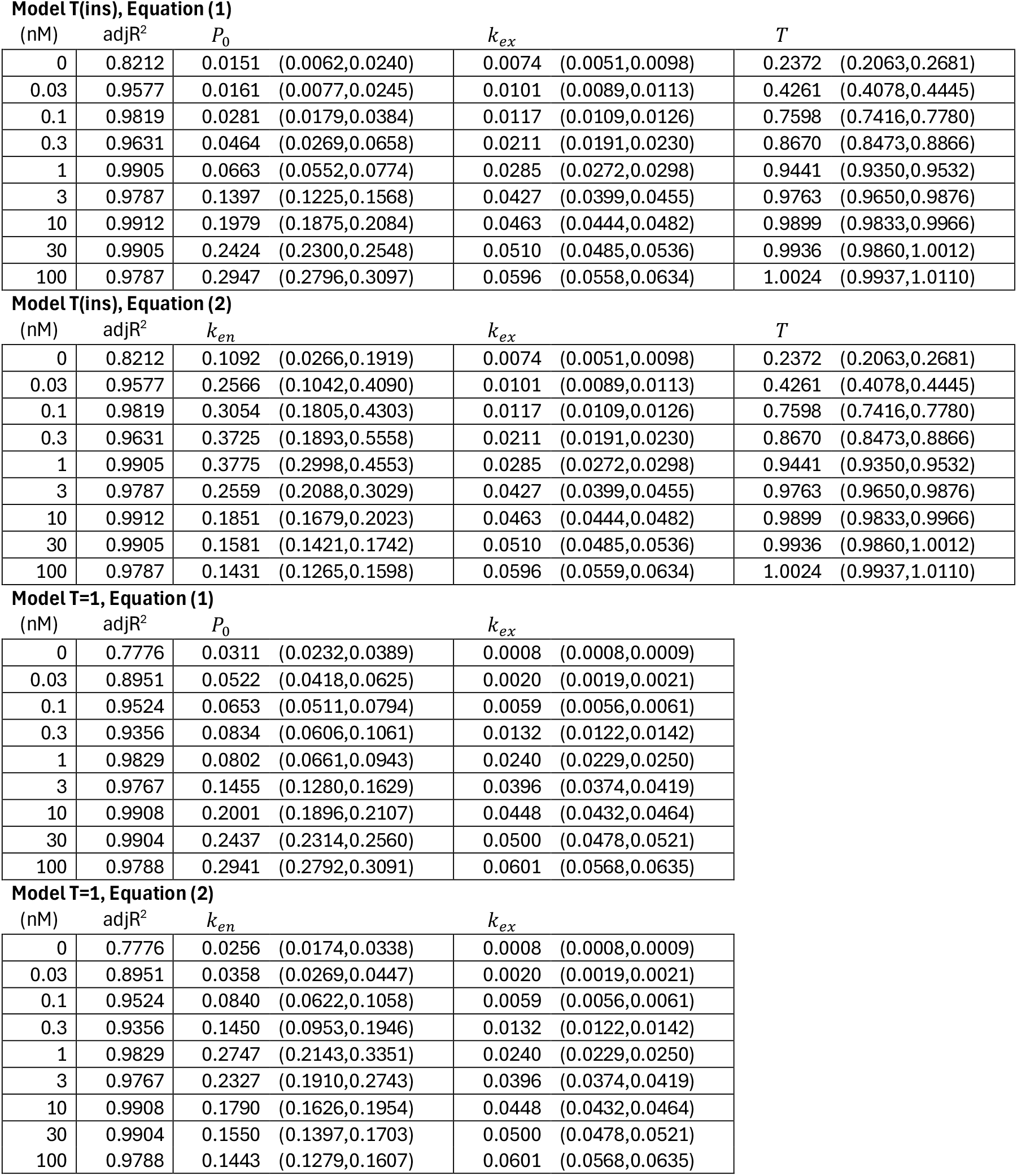
Parameter values for the least-squares fits to the uptake data. The 95% confidence interval is reported in brackets. The adjusted R^2^ value (adjR^2^) accounts for the number of degrees of freedom in the model.

**Model T(ins), Equation (1)**
| (nM) | adjR<sup>2</sup> | $P_0$ | $k_{ex}$ | $T$ |
| --- | --- | --- | --- | --- |
| 0 | 0.8212 | 0.0151 (0.0062,0.0240) | 0.0074 (0.0051,0.0098) | 0.2372 (0.2063,0.2681) |
| 0.03 | 0.9577 | 0.0161 (0.0077,0.0245) | 0.0101 (0.0089,0.0113) | 0.4261 (0.4078,0.4445) |
| 0.1 | 0.9819 | 0.0281 (0.0179,0.0384) | 0.0117 (0.0109,0.0126) | 0.7598 (0.7416,0.7780) |
| 0.3 | 0.9631 | 0.0464 (0.0269,0.0658) | 0.0211 (0.0191,0.0230) | 0.8670 (0.8473,0.8866) |
| 1 | 0.9905 | 0.0663 (0.0552,0.0774) | 0.0285 (0.0272,0.0298) | 0.9441 (0.9350,0.9532) |
| 3 | 0.9787 | 0.1397 (0.1225,0.1568) | 0.0427 (0.0399,0.0455) | 0.9763 (0.9650,0.9876) |
| 10 | 0.9912 | 0.1979 (0.1875,0.2084) | 0.0463 (0.0444,0.0482) | 0.9899 (0.9833,0.9966) |
| 30 | 0.9905 | 0.2424 (0.2300,0.2548) | 0.0510 (0.0485,0.0536) | 0.9936 (0.9860,1.0012) |
| 100 | 0.9787 | 0.2947 (0.2796,0.3097) | 0.0596 (0.0558,0.0634) | 1.0024 (0.9937,1.0110) |

**Model T(ins), Equation (2)**
| (nM) | adjR<sup>2</sup> | $k_{en}$ | $k_{ex}$ | $T$ |
| --- | --- | --- | --- | --- |
| 0 | 0.8212 | 0.1092 (0.0266,0.1919) | 0.0074 (0.0051,0.0098) | 0.2372 (0.2063,0.2681) |
| 0.03 | 0.9577 | 0.2566 (0.1042,0.4090) | 0.0101 (0.0089,0.0113) | 0.4261 (0.4078,0.4445) |
| 0.1 | 0.9819 | 0.3054 (0.1805,0.4303) | 0.0117 (0.0109,0.0126) | 0.7598 (0.7416,0.7780) |
| 0.3 | 0.9631 | 0.3725 (0.1893,0.5558) | 0.0211 (0.0191,0.0230) | 0.8670 (0.8473,0.8866) |
| 1 | 0.9905 | 0.3775 (0.2998,0.4553) | 0.0285 (0.0272,0.0298) | 0.9441 (0.9350,0.9532) |
| 3 | 0.9787 | 0.2559 (0.2088,0.3029) | 0.0427 (0.0399,0.0455) | 0.9763 (0.9650,0.9876) |
| 10 | 0.9912 | 0.1851 (0.1679,0.2023) | 0.0463 (0.0444,0.0482) | 0.9899 (0.9833,0.9966) |
| 30 | 0.9905 | 0.1581 (0.1421,0.1742) | 0.0510 (0.0485,0.0536) | 0.9936 (0.9860,1.0012) |
| 100 | 0.9787 | 0.1431 (0.1265,0.1598) | 0.0596 (0.0559,0.0634) | 1.0024 (0.9937,1.0110) |

**Model T=1, Equation (1)**
| (nM) | adjR<sup>2</sup> | $P_0$ | $k_{ex}$ |
| --- | --- | --- | --- |
| 0 | 0.7776 | 0.0311 (0.0232,0.0389) | 0.0008 (0.0008,0.0009) |
| 0.03 | 0.8951 | 0.0522 (0.0418,0.0625) | 0.0020 (0.0019,0.0021) |
| 0.1 | 0.9524 | 0.0653 (0.0511,0.0794) | 0.0059 (0.0056,0.0061) |
| 0.3 | 0.9356 | 0.0834 (0.0606,0.1061) | 0.0132 (0.0122,0.0142) |
| 1 | 0.9829 | 0.0802 (0.0661,0.0943) | 0.0240 (0.0229,0.0250) |
| 3 | 0.9767 | 0.1455 (0.1280,0.1629) | 0.0396 (0.0374,0.0419) |
| 10 | 0.9908 | 0.2001 (0.1896,0.2107) | 0.0448 (0.0432,0.0464) |
| 30 | 0.9904 | 0.2437 (0.2314,0.2560) | 0.0500 (0.0478,0.0521) |
| 100 | 0.9788 | 0.2941 (0.2792,0.3091) | 0.0601 (0.0568,0.0635) |

| (nM) | adjR<sup>2</sup> | $k_{en}$ | $k_{ex}$ |
| --- | --- | --- | --- |
| 0 | 0.7776 | 0.0256 (0.0174,0.0338) | 0.0008 (0.0008,0.0009) |
| 0.03 | 0.8951 | 0.0358 (0.0269,0.0447) | 0.0020 (0.0019,0.0021) |
| 0.1 | 0.9524 | 0.0840 (0.0622,0.1058) | 0.0059 (0.0056,0.0061) |
| 0.3 | 0.9356 | 0.1450 (0.0953,0.1946) | 0.0132 (0.0122,0.0142) |
| 1 | 0.9829 | 0.2747 (0.2143,0.3351) | 0.0240 (0.0229,0.0250) |
| 3 | 0.9767 | 0.2327 (0.1910,0.2743) | 0.0396 (0.0374,0.0419) |
| 10 | 0.9908 | 0.1790 (0.1626,0.1954) | 0.0448 (0.0432,0.0464) |
| 30 | 0.9904 | 0.1550 (0.1397,0.1703) | 0.0500 (0.0478,0.0521) |
| 100 | 0.9788 | 0.1443 (0.1279,0.1607) | 0.0601 (0.0568,0.0635) |

**Figure 2.**
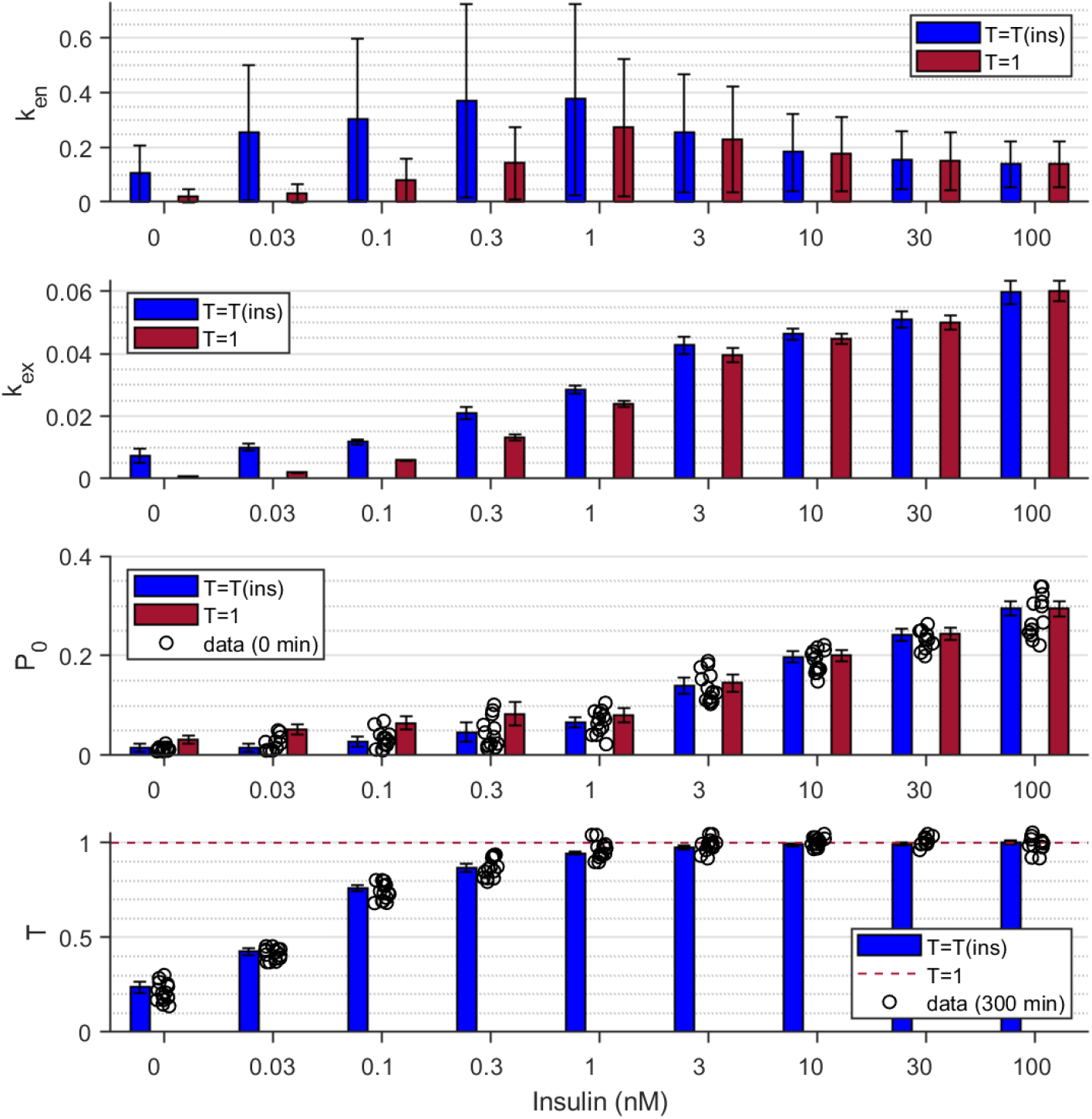
Parameter values of the least-squares fits of the Uptake Assay to the T(ins) and T=1 models as a function of insulin concentration (categorical axis). The error bars indicate the 95% confidence intervals for the parameter values. The initial plasma membrane level, P_0_, inferred in each model is also compared to the uptake data at 0 minutes, when the surface GLUT4 is initially labeled. Similarly, the data at 300 minutes is shown in comparison to the inferred recycling total. Note that the two models are practically identical for 100nM. The parameter *k*_*en*_ was inferred using Equation (2). Note that identical values for *k*_*ex*_, *P*_0_, and *T* were inferred using both Equations (1) and (2).

As Model T=1 has one fewer parameter than Model T(ins), the Adjusted R^2^ of the fits (which takes into account the number of parameters in each model) was determined to allow a comparison of the relative goodness of fit. Whilst neither model perfectly fitted the basal uptake data, Model T(ins) had higher Adjusted R^2^ values for doses up to 1nM compared to Model T=1.

Note that the Adjusted R^2^ are very similar in the two model variants for high insulin doses. Indeed, at and above 3nM they are practically indistinguishable. Allowing the plateau level to vary with insulin results in improvements in the fits at the lower insulin doses – in particular for the basal data. Additionally, the residuals across each dose response for each assay were visualized to determine whether the distribution of the residuals was uniformly distributed about zero at each time during the experiment, Figure 3. In general, better models have residuals that are also uniform across the time course. Note that the results at the maximal, 100nM, insulin dose are practically identical, as the normalization of the experiment causes the plateau to occur at *T* = 1 for this dose.

**Figure 3.**
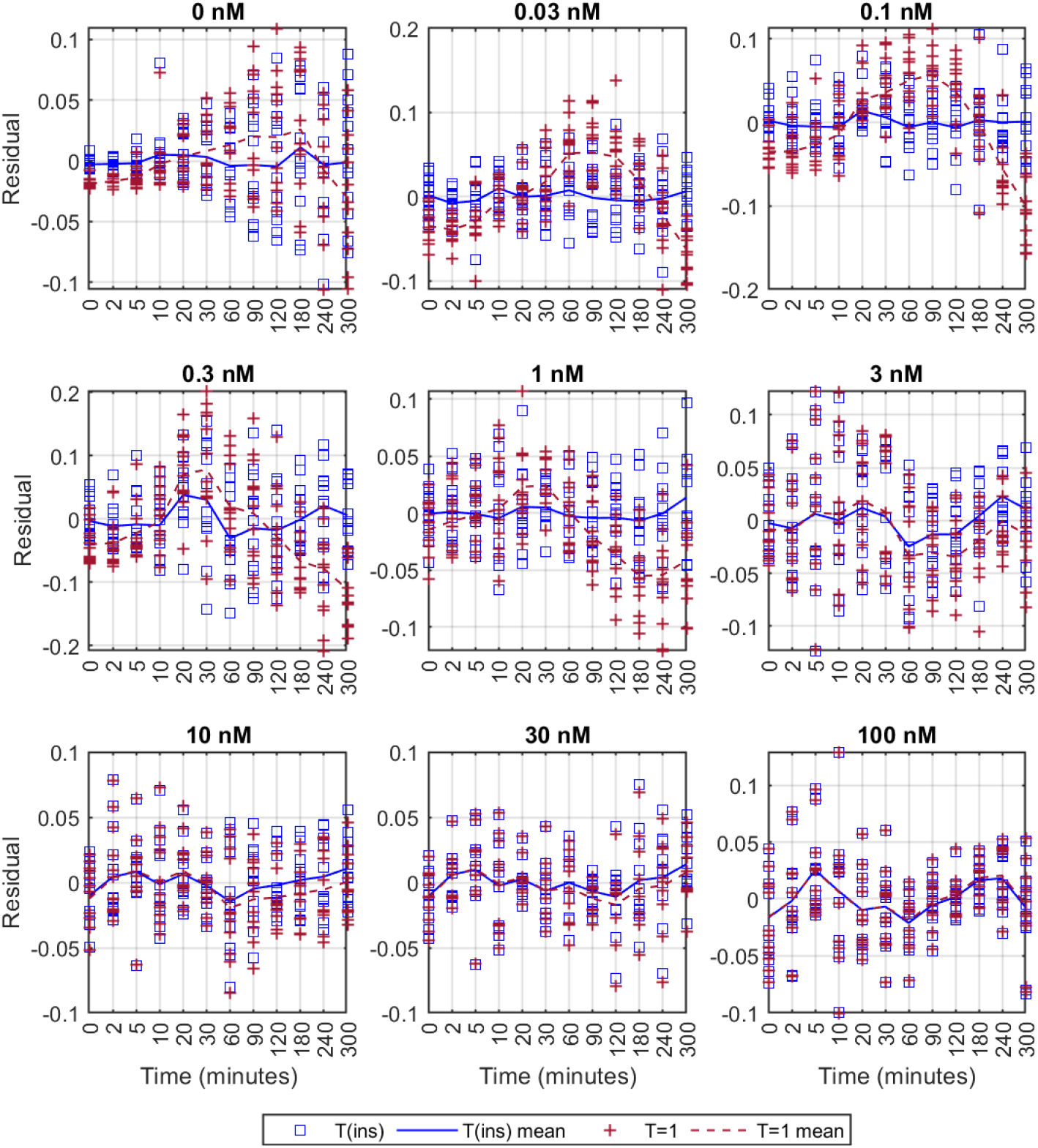
Residuals of the least-squares fits for the Uptake Assay to the T(ins) and T=1 models as a function of the time of the data for each insulin concentration (indicated above each subplot). Note the time scale is categorical. The residuals are for each individual data point are shown as squares (T(ins)) and pluses (T=1), with their mean values at each time connected as solid (T(ins)) and dashed (T=1) lines. Note, good models and fits produce residuals that are symmetric about zero and there should not be observable trends across the time course. Note also that the two models were practically identical for 100nM and thus have the same residual structure. Here Equation (2) was used for the fits, but identical results were achieved using Equation (1).

The better fit of Model T(ins) at sub-maximal insulin doses is also observed in the residuals for the fits, Figure 3. The residuals for the Model T=1 were more widely spread about zero, particularly at low insulin doses, compared to those for Model T(ins).

The distribution of all the residuals across the entire time course of the assay are shown for the two models in Figure 4, where it can be seen that the distribution is broader and skewed away from zero for Model T=1 compared to Model T(ins) for most of the insulin doses.

**Figure 4.**
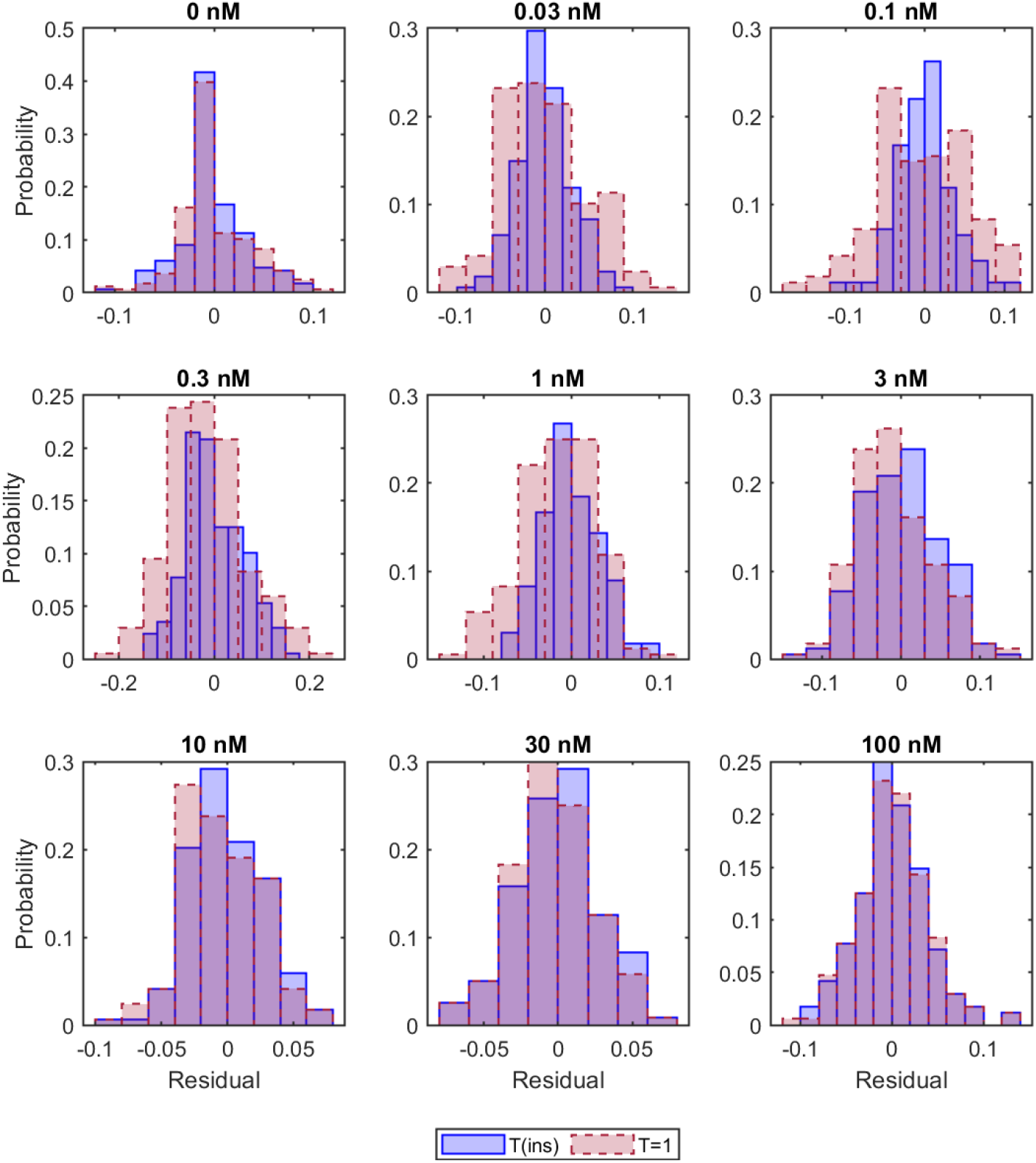
Overall residuals of the least-squares fits for the Uptake Assay to the T(ins) and T=1 models as a function of the time of the data for each insulin concentration (indicated above each subplot). Note, good models and fits produce residuals that are symmetric about zero. Note also that the two models were practically identical for 100nM and thus have the same residual structure. Here Equation (2) was used for the fits, but identical results were achieved using Equation (1).

The improved fit of the data with an insulin-dependent plateau indicates that the amount of GLUT4 recycling in the system may be controlled by the level of insulin present.

### Is endocytosis constant with insulin?

It can be seen in Figures 1 and 2 and Table 2 that for both Model T(ins) and Model T=1 the exocytosis rate, *k*_*ex*_, increases with insulin. There are also changes in the endocytosis rate, *k*_*en*_, with insulin, but with a less discernable trend. It has been previously reported that the endocytosis rate is relatively insulin independent (e.g., (11)). To test whether a model where the endocytosis rate is insulin independent is a reasonable description of the system we undertake least-squares optimization of uptake data from all doses simultaneously using a single endocytosis rate, while allowing the exocytosis rates to be insulin-dependent, i.e., the output of the uptake assay data, *L* (*i, t*) is described as

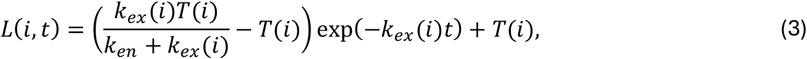

for insulin, *i* = 0, 0.03, 0.1, 0.3, 1, 3, 10, 30, 100nM, and where *t* is time.

Again, we can consider two cases, Model T(ins), k_en_, in which the total recycling, *T* (*i*), is insulin dependent, and Model T=1, *k*_*en*_, in which *T* (*i*) = 1, for all *i*.

The fits of these models to the uptake assay data are shown in Figures 5 and Table 3. The same trends were observed in the fits to these models as were in the fits with independent endocytosis rates (which were only constrained by the data at the individual insulin doses). Model T(ins) and Model T(ins), *k*_*en*_ showed very few differences at all insulin doses, as can be seen from the mean residuals of the fits, Figure 6. All models were practically identical at high insulin doses, however the correspondence of both models with fixed *T* = 1 showed more deviations from the data at lower insulin levels, particularly the basal state.

**Table 3.**
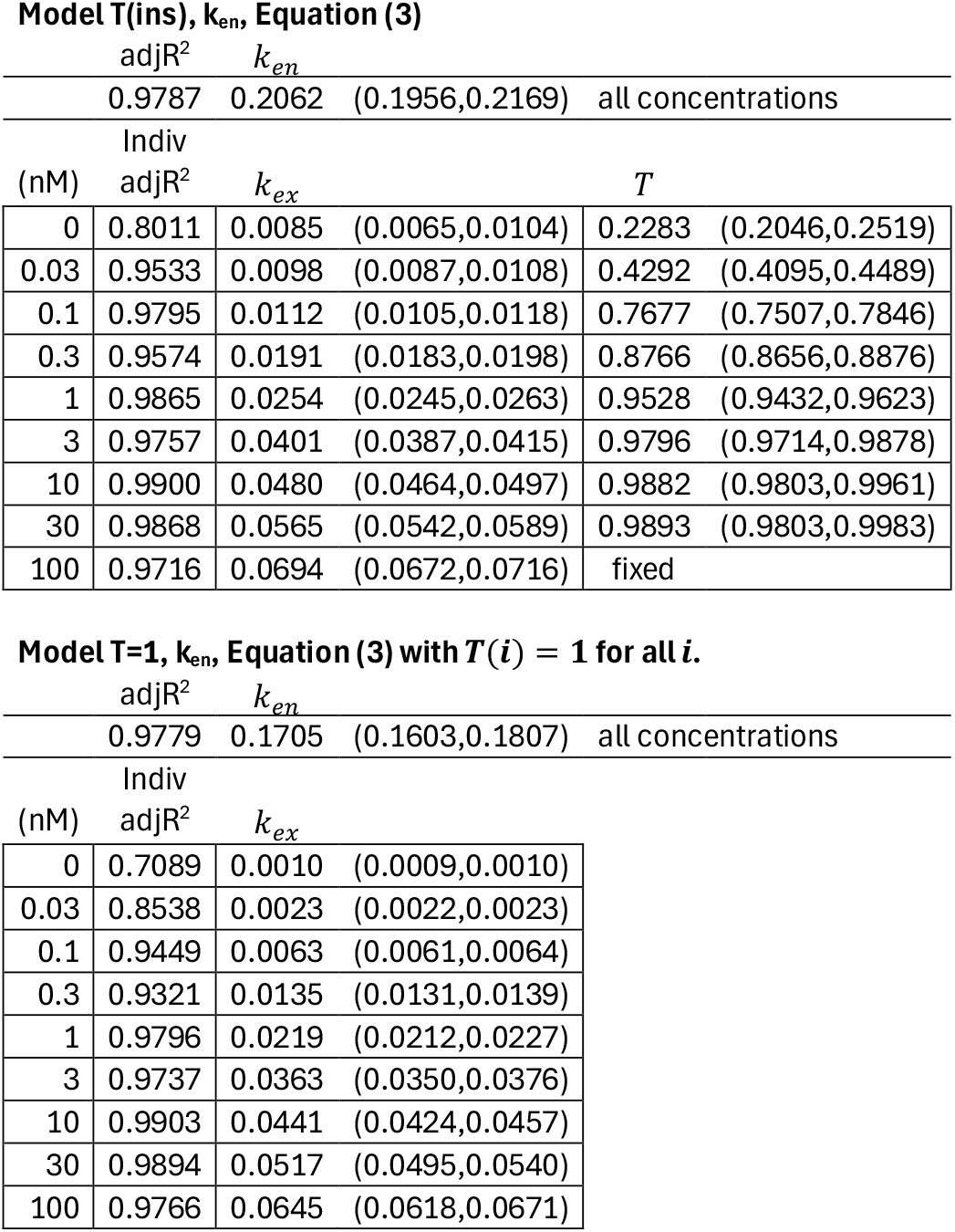
Parameter values for the least-squares fits to the uptake data. The 95% confidence interval is reported in brackets. The adjusted R^2^ value (adjR^2^) accounts for the number of degrees of freedom in the model. The Indiv adjR^2^ assesses the fit on the subset of data for that insulin dose.

**Figure 5.**
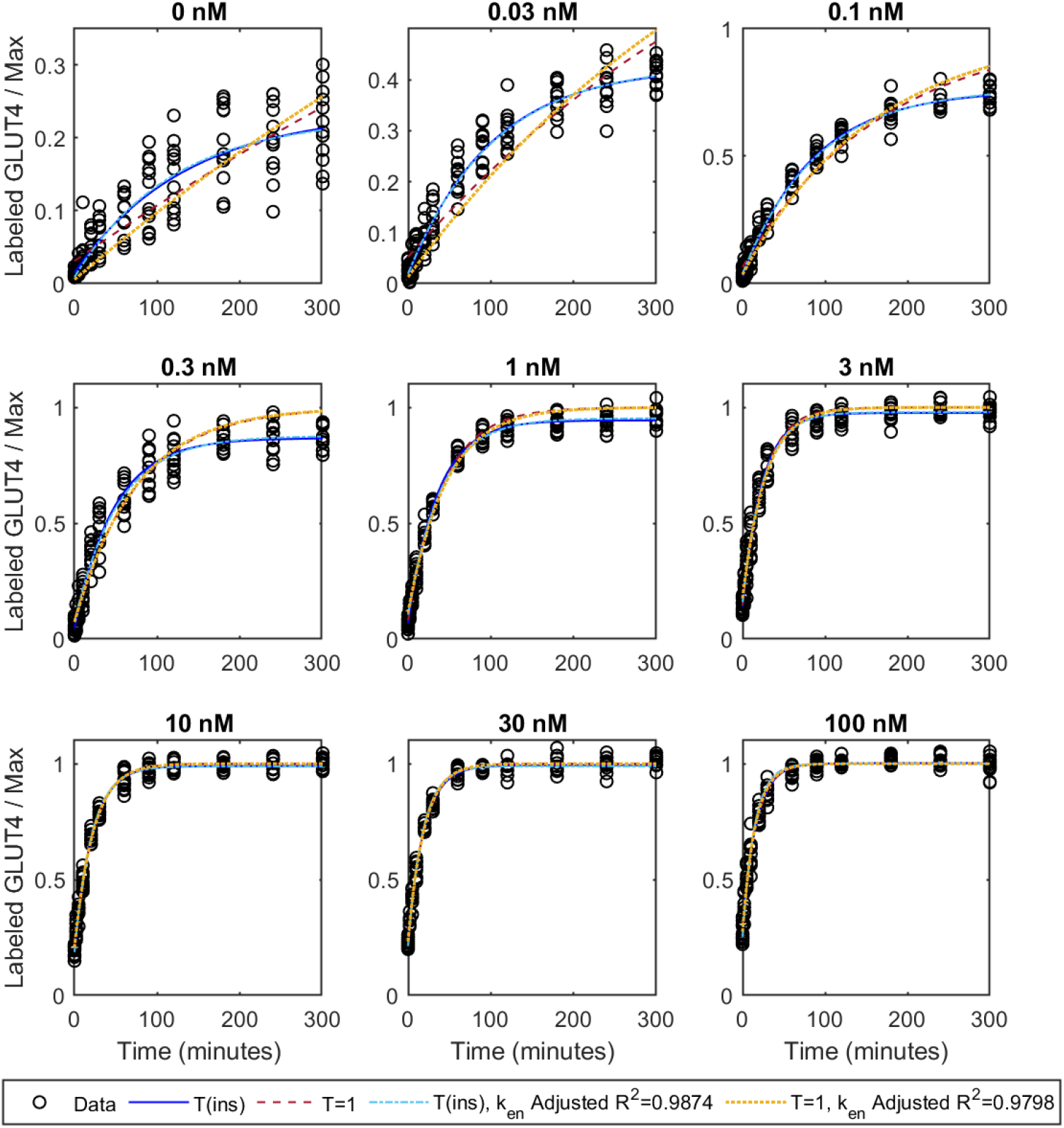
Uptake Assay as a function of the time for each insulin concentration. Least-squares fits of Models T(ins), k_en_ and T=1, k_en_ in which the endocytosis rate was independent of insulin are shown. Note these fits were constrained by the data at all insulin concentrations. The overall Adjusted R^2^ for these fits is shown in the legend. The fits to Models T(ins), and T=1 (Figure 1) are shown for comparison.

**Figure 6.**
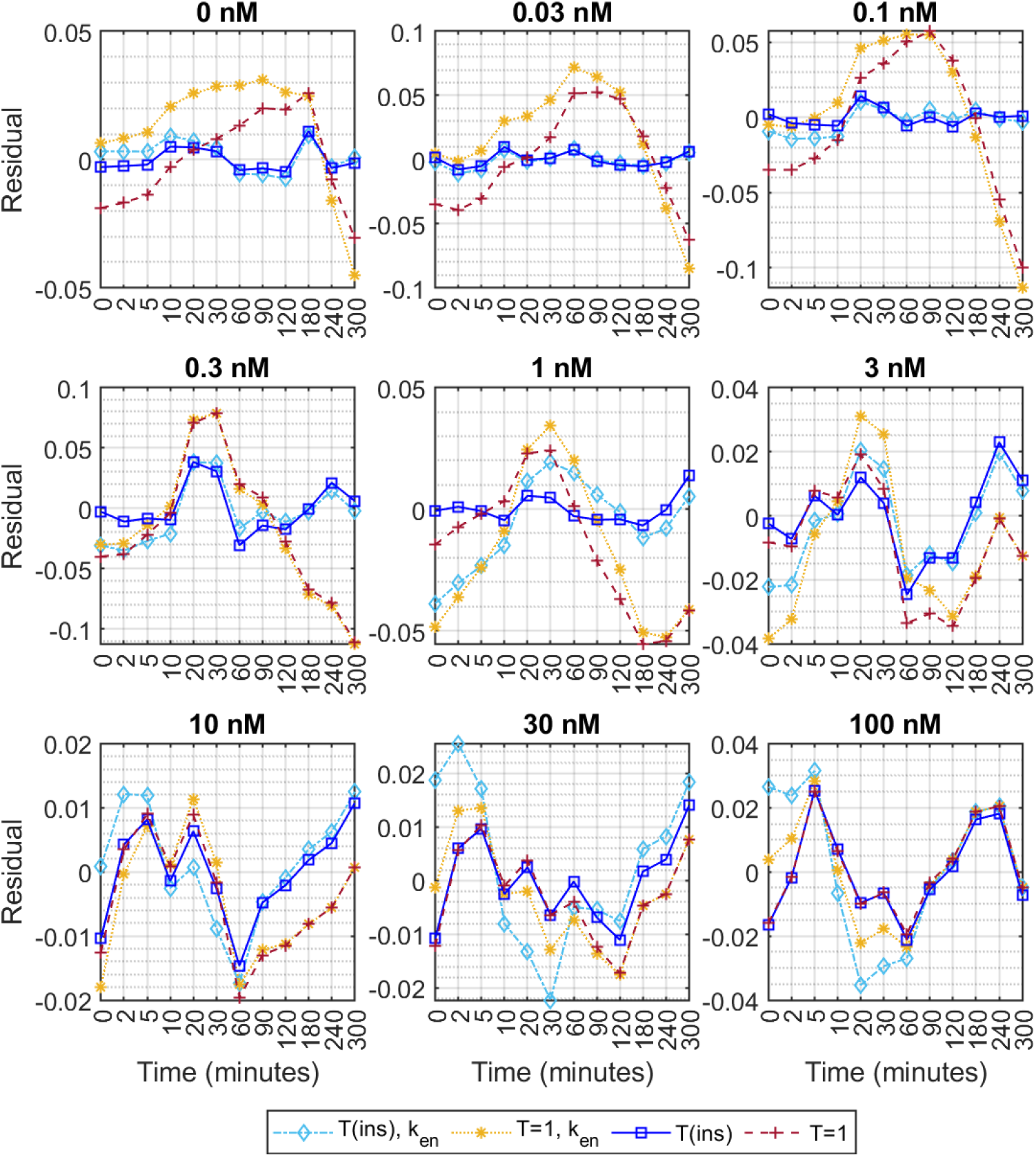
Mean residuals of the least-squares fits for the Uptake Assay to the T(ins), k_en_ and T=1, k_en_ models as a function of the time of the data for each insulin concentration (indicated above each subplot). Note the time scale is categorical. The mean residuals for the individual T(ins) and T=1 models (Figure 3) are shown for comparison.

The overall Adjusted R^2^ for Model T(ins), *k*_*en*_, was 0.9874 and 0.9798 for Model T=1, *k*_*en*_, i.e., commensurate with the Adjusted R^2^ values for Model T(ins) and Model T=1 at each insulin concentration (inset in Figure 5, and Table 1). Thus, it is plausible that the endocytosis rate could be independent of the insulin dose, which concurs with previous findings, e.g., (1,2,11).

Note that it is not possible to replicate the observed dynamics if the exocytosis rate is assumed to be insulin independent, even if the endocytosis rate is insulin dependent. This is because exocytosis is the primary determinant of the change in labeling in the assay. Whilst fits with *T* dependent on insulin can replicate the observations, the endocytosis rates required are unphysically large (data not shown).

### Transition Experiments

The Transition Assay uses surface labelling to track the plasma membrane level of GLUT4 as the system adjusts from the basal insulin steady state in response to the application of a given concentration of insulin. If there is sequestration of GLUT4 away from the recycling pathway the total amount recycling could also change. In these experiments the observations are normalized to the mean plasma membrane level at 60 min when the maximal level (100nM) of insulin was applied. Figure 7 shows that the data for each concentration rose to a plateau with a single constant, i.e.,

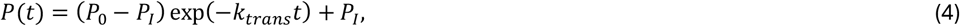

where *P*(*t*) is the amount of GLUT4 at the plasma membrane at time *t, P*_0_ is the initial level of labeled GLUT4 at the plasma membrane in the basal insulin state, *P*_*I*_ is the plateau level, corresponding to the amount of GLUT4 at the plasma membrane when the system is in the steady state in the presence of insulin, and *k* is the rate at which the transition from the basal to insulin-stimulated steady state occurs.

**Figure 7.**
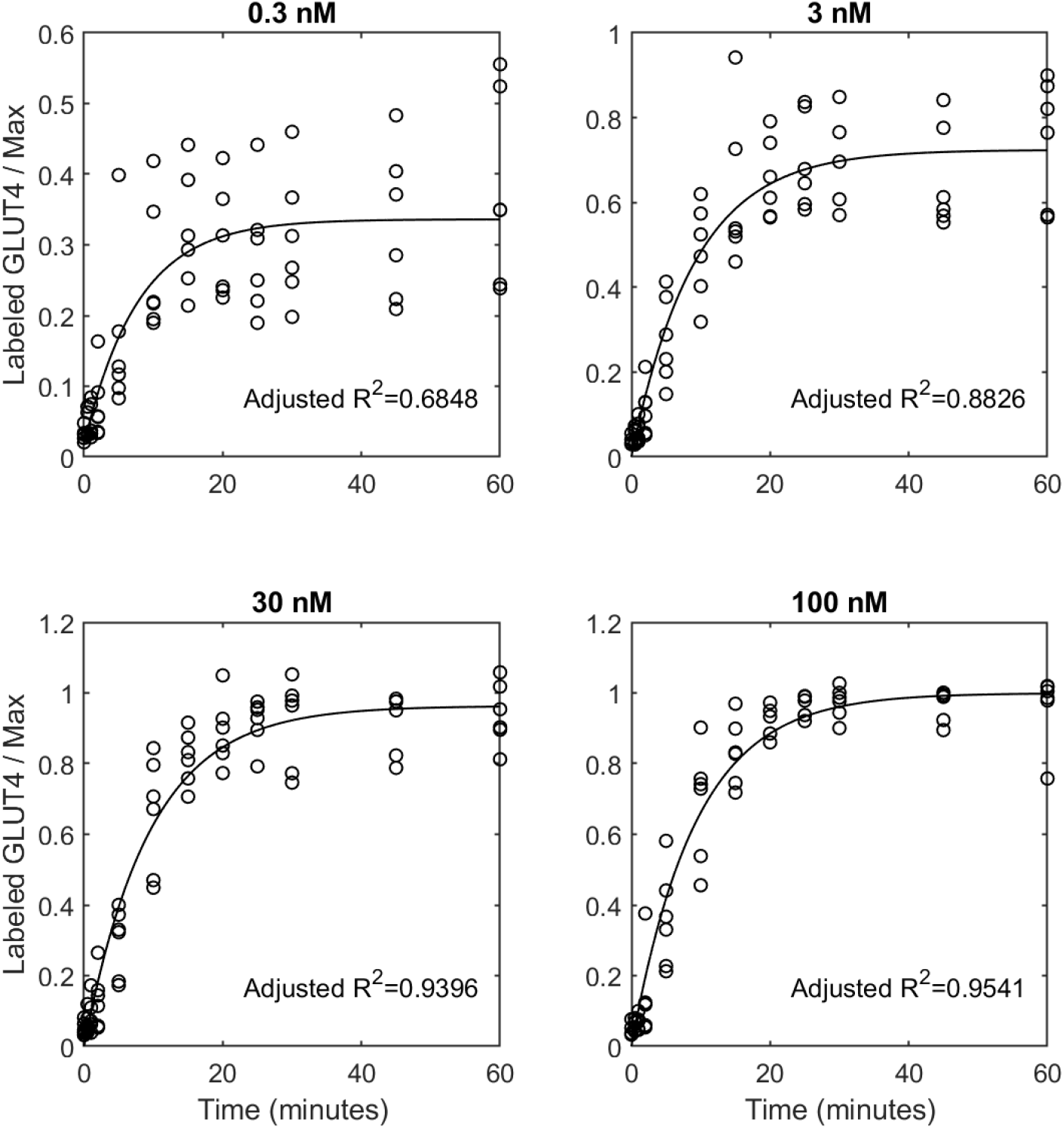
Least-squares fits for the Transition Assay as a function of the time of the data for each insulin concentration. Note the data was quite noisy at 0.3nM leading to a reduced Adjusted R^2^ for this data set.

Rather than a rate constant for the process, we can also equivalently fit a time constant for the transition, *τ*_*trans*_, i.e.,

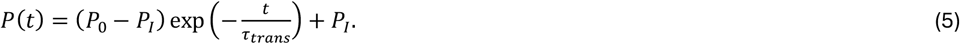

The results and parameter values for the least-squares fits to Equations (4) and (5) for the GLUT4 transition experiment when 0.3, 3, 30, and 100nM insulin were applied are shown in Figure 7 and Table 4. The fits indicate that the half-time for the transition between the basal and insulin-stimulated steady states range between 5.4 to 6.6 minutes. The transition rates are similar to the inferred endocytosis rates in the Uptake assay, particularly at the higher insulin doses.

**Table 4.**
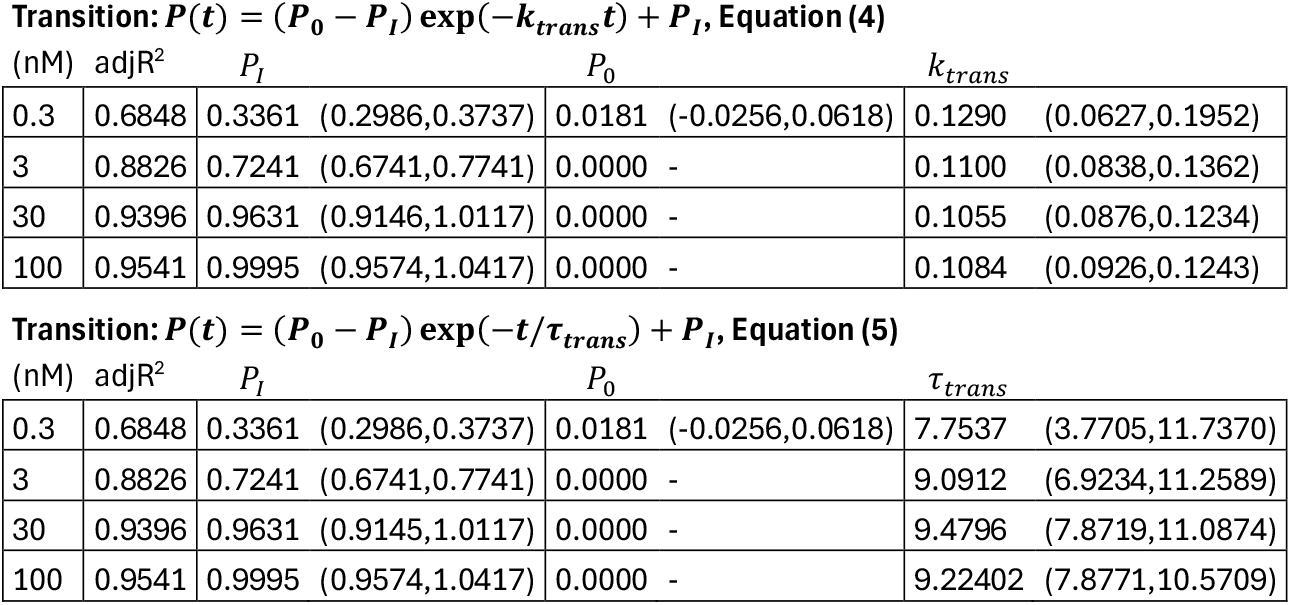
Parameter values for the least-squares fit to Equations (4) and (5) for the Transition data. The 95% confidence interval is reported in brackets or with a dash if the value was fixed at the bound. The adjusted R^2^ value accounts for the number of degrees of freedom in the model. Note that the inferred values of the common parameters between the two versions of the fits are identical.

**Transition: $P(t) = (P_0 - P_I) \exp(-k_{trans}t) + P_I$ , Equation (4)**
| (nM) | adjR<sup>2</sup> | $P_I$ | $P_0$ | $k_{trans}$ |
| --- | --- | --- | --- | --- |
| 0.3 | 0.6848 | 0.3361 (0.2986,0.3737) | 0.0181 (-0.0256,0.0618) | 0.1290 (0.0627,0.1952) |
| 3 | 0.8826 | 0.7241 (0.6741,0.7741) | 0.0000 - | 0.1100 (0.0838,0.1362) |
| 30 | 0.9396 | 0.9631 (0.9146,1.0117) | 0.0000 - | 0.1055 (0.0876,0.1234) |
| 100 | 0.9541 | 0.9995 (0.9574,1.0417) | 0.0000 - | 0.1084 (0.0926,0.1243) |

Table 4. Parameter values for the least-squares fit to Equations (4) and (5) for the Transition data.
| (nM) | adjR<sup>2</sup> | $P_I$ | $P_0$ | $\tau_{trans}$ |
| --- | --- | --- | --- | --- |
| 0.3 | 0.6848 | 0.3361 (0.2986,0.3737) | 0.0181 (-0.0256,0.0618) | 7.7537 (3.7705,11.7370) |
| 3 | 0.8826 | 0.7241 (0.6741,0.7741) | 0.0000 - | 9.0912 (6.9234,11.2589) |
| 30 | 0.9396 | 0.9631 (0.9145,1.0117) | 0.0000 - | 9.4796 (7.8719,11.0874) |
| 100 | 0.9541 | 0.9995 (0.9574,1.0417) | 0.0000 - | 9.22402 (7.8771,10.5709) |

### Inhibiting the exocytosis of GLUT4

Similarly to the steady state assay, in the inhibition assay the system was stabilsed into a steady state in the presence of a given level of insulin. The experiment tracks the plasma membrane level of GLUT4 as a function of time after the application of a PI3K exocytosis inhibitor, LY294002 (LYi), which inhibits the rate constant of exocytosis by approximately 90% (11). The observations are the plasma membrane level of GLUT4, normalized to the steady state level in 100nM insulin alone. It can be seen in Figure 8 that the system evolves as an exponential decay from the plasma membrane level in steady state, i.e.,

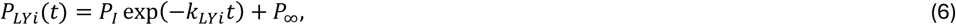

or equivalently,

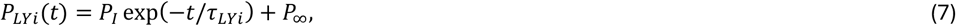

where *P*_*LYi*_ (*t*) is the amount of GLUT4 at the plasma membrane at time *t, P*_*I*_ is the initial level of labeled GLUT4 at the plasma membrane in the insulin steady state, *P*_∞_ is the plateau level, corresponding to the amount of GLUT4 at the plasma membrane when the system is in the steady state in the presence of LYi, *k*_*en*_ is the rate constant for removal from the plasma membrane and *τ*_*LYi*_ the equivalent time constant at which the transition to this steady state occurs.

**Figure 8.**
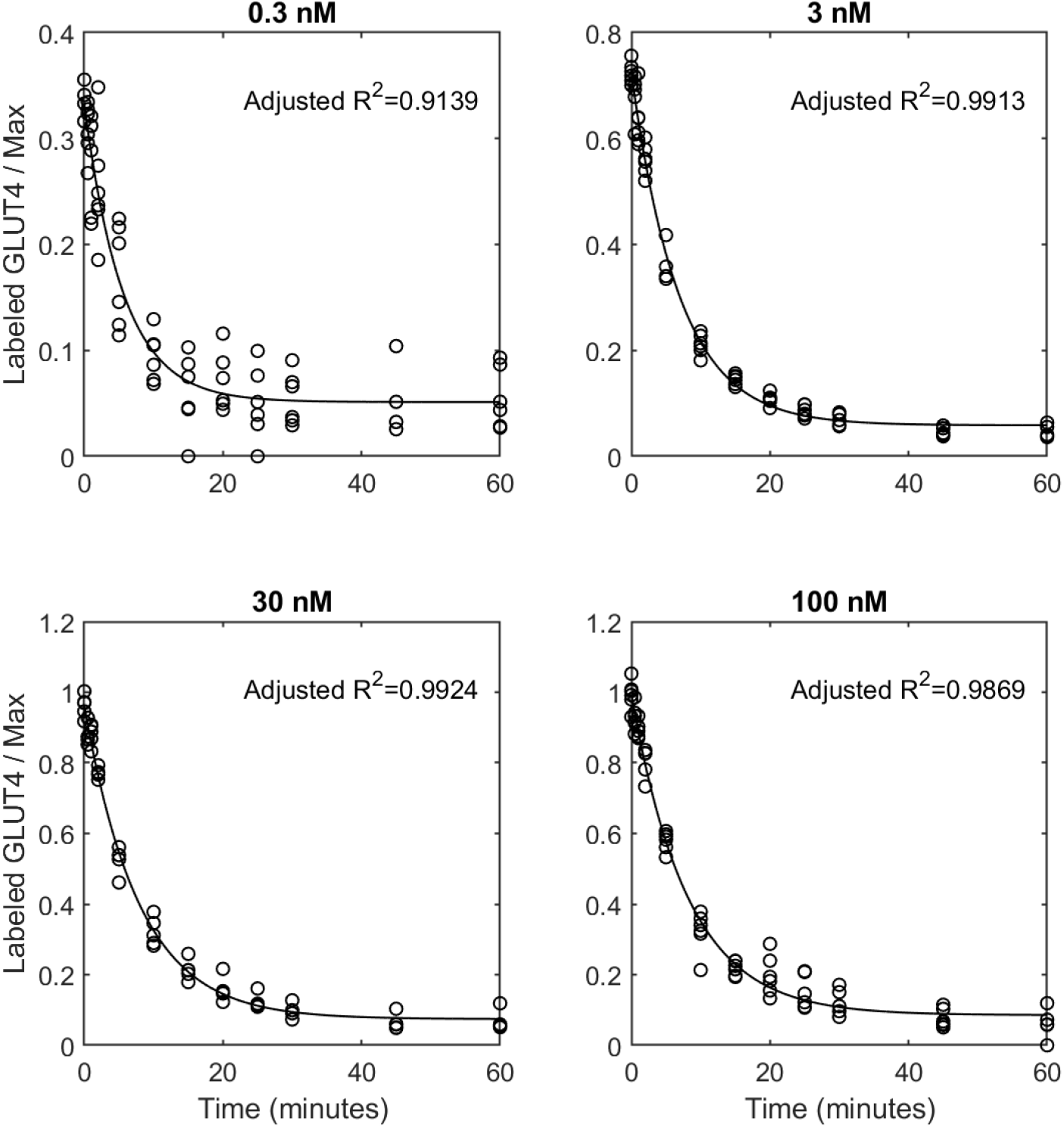
Least-squares fits for the Inhibition Assay as a function of the time of the data for each insulin concentration. The Adjusted R^2^ for the fit to each concentration is indicated on the subfigures.

The results and parameter values for the least-squares optimized fits to Equations (6) and (7) for the GLUT4 inhibition experiment in the presence of 0.3, 3, 30, and 100nM insulin are shown in Figure 8 and Table 5. The half-times of the inhibition time courses range between 3.9 and 5.6 minutes, and, similarly to the Transition assay, the rate constants were comparable to the endocytosis rates inferred in the Uptake assay fits (Table 1), commensurate with this assay predominantly being driven by endocytosis (11,12).

**Table 5.**
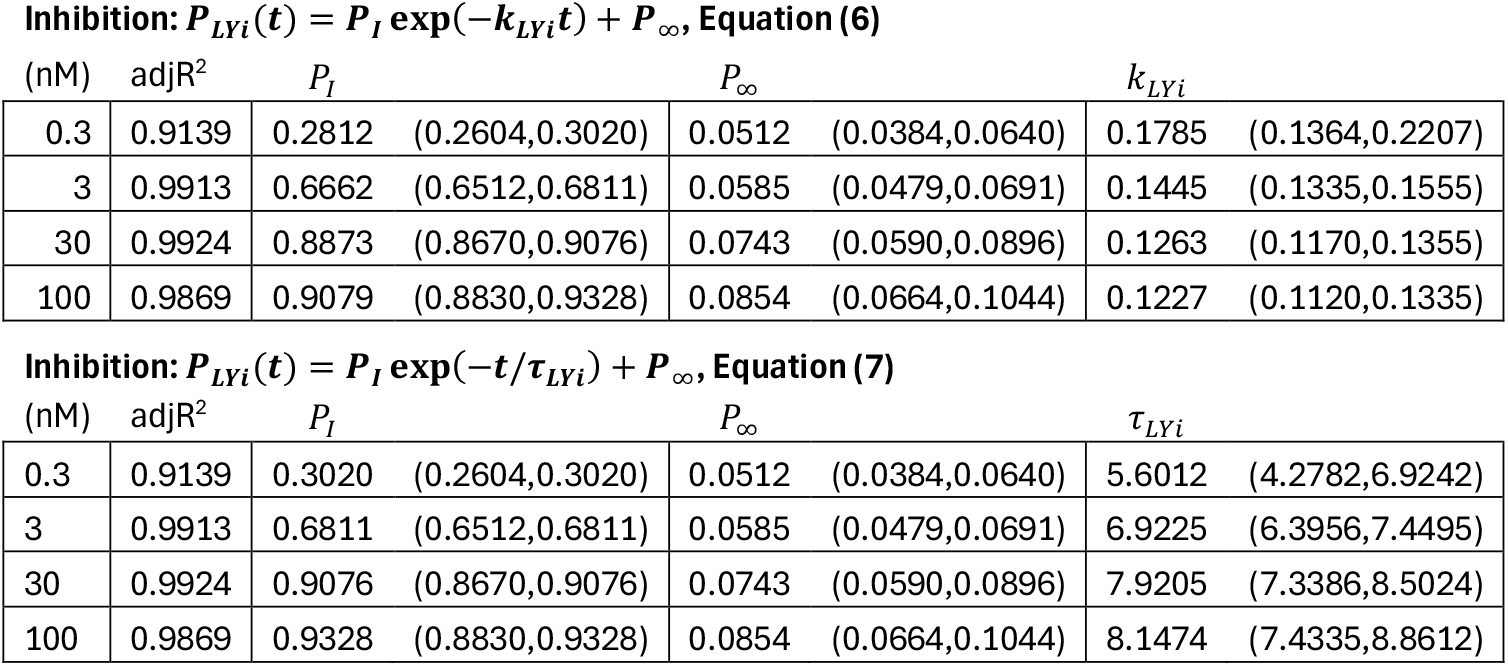
Parameter values for the least-squares fit to Equations (6) and (7) for the inhibition assay data. The 95% confidence interval is reported in brackets. The adjusted R^2^ value accounts for the number of degrees of freedom in the model. Note that the inferred values of the common parameters between the two versions of the fits are identical.

**Inhibition: $P_{LYi}(t) = P_I \exp(-k_{LYi}t) + P_\infty$ , Equation (6)**
| (nM) | adjR<sup>2</sup> | $P_I$ | $P_\infty$ | $k_{LYi}$ |
| --- | --- | --- | --- | --- |
| 0.3 | 0.9139 | 0.2812 (0.2604,0.3020) | 0.0512 (0.0384,0.0640) | 0.1785 (0.1364,0.2207) |
| 3 | 0.9913 | 0.6662 (0.6512,0.6811) | 0.0585 (0.0479,0.0691) | 0.1445 (0.1335,0.1555) |
| 30 | 0.9924 | 0.8873 (0.8670,0.9076) | 0.0743 (0.0590,0.0896) | 0.1263 (0.1170,0.1355) |
| 100 | 0.9869 | 0.9079 (0.8830,0.9328) | 0.0854 (0.0664,0.1044) | 0.1227 (0.1120,0.1335) |

Table 5 Parameter values for the least-squares fit to Equations (6) and (7) for the inhibition assay data.
| (nM) | adjR<sup>2</sup> | $P_I$ | $P_\infty$ | $\tau_{LYi}$ |
| --- | --- | --- | --- | --- |
| 0.3 | 0.9139 | 0.3020 (0.2604,0.3020) | 0.0512 (0.0384,0.0640) | 5.6012 (4.2782,6.9242) |
| 3 | 0.9913 | 0.6811 (0.6512,0.6811) | 0.0585 (0.0479,0.0691) | 6.9225 (6.3956,7.4495) |
| 30 | 0.9924 | 0.9076 (0.8670,0.9076) | 0.0743 (0.0590,0.0896) | 7.9205 (7.3386,8.5024) |
| 100 | 0.9869 | 0.9328 (0.8830,0.9328) | 0.0854 (0.0664,0.1044) | 8.1474 (7.4335,8.8612) |

**Table 6.**
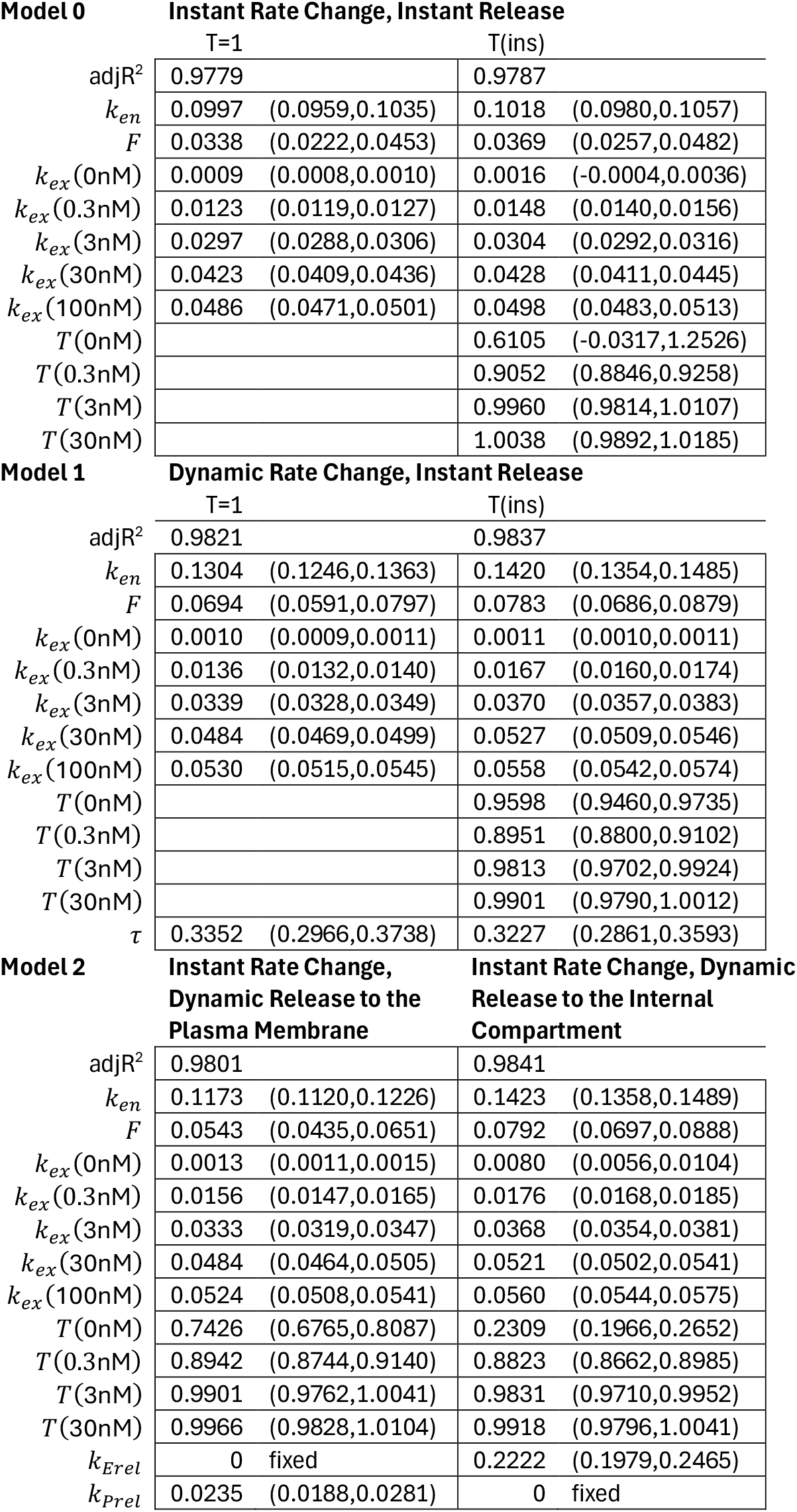
Parameter values of the different model hypotheses from the least-squares fits using all the assay data simultaneously. The 95% confidence interval is reported in brackets. The adjusted R^2^ value (adjR^2^) accounts for the number of degrees of freedom in the models.

### Part 2: Parsimonious Models of GLUT4 Recycling

It was seen in the independent fits to the different assay data that the rate constants for endocytosis in the uptake assay, the transition rate constant, and the inhibition transition rate were similar. It also appears that it is plausible that some GLUT4 is sequestered from the recycling pathway in sub-maximal levels of insulin.

Rather than fitting the data using individual assay models, the data from the assays can be used to simultaneously constrain models of GLUT4 recycling, to test hypotheses about the dynamics of the system.

Taking a parsimonious approach, we consider GLUT4 recycling between a single compartment representing internal membrane structures and one representing the plasma membrane. The model here considers one overall exocytosis rate, representing the combination of all exocytotic processes, and similarly a single endocytosis rate representing all endocytic processes. Sequestered amounts, if any, are held in a separate compartment, and may be released upon the application of insulin. If the system is in steady state with respect to insulin, we assume that any release has been completed, and that the amount of GLUT4 recycling between the internal structures and the plasma membrane is constant.

We assume that there is negligible re-sequestration of GLUT4 occurring over the timescale of the experiments, i.e., sequestered amounts, if any, are only released during the observations. A schematic of the model is shown in Figure 9.

**Figure 9.**
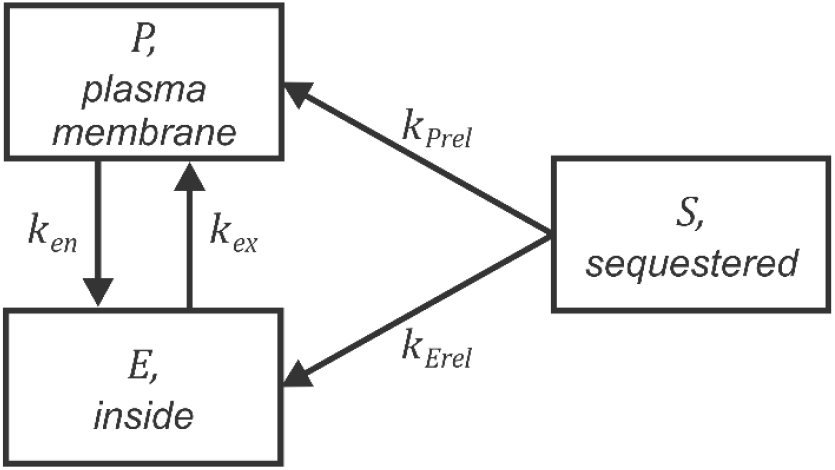
Schematic of the model of GLUT4 recycling. GLUT4 can be present in one of three compartments: the plasma membrane, *P*, internal membrane structures, such as endosomes, *E*, and sequestered structures, *S*. GLUT4 is exocytosed at a rate *k*_*ex*_ and endocytosed at rate *k*_*en*_, and two possible release pathways are considered, from the sequestered structures to the plasma membrane at a rate *k*_*prel*_, and to the internal structures at a rate *k*_*Erel*_. Note that it is assumed that re-sequestration of GLUT4 into storage compartments is negligible, occurring over a much longer timescale than the experimental observations.

The dynamics are given by

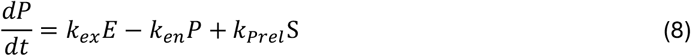

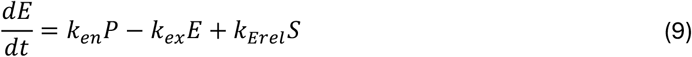

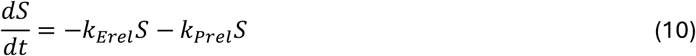

where *t* is time,

*P* is the amount of GLUT4 at the plasma membrane,

*E* is the amount of GLUT4 in the internal compartment,

*S* is the amount of sequestered GLUT4,

*k*_*ex*_ is the exocytosis rate from the internal compartment to the plasma membrane,

*k*_*en*_ is the endocytosis rate from the plasma membrane to the internal compartment,

*k*_*prel*_ is the rate of release of sequestered GLUT4 to the plasma membrane, and

*k*_*Erel*_ is the rate of release of sequestered GLUT4 to the internal compartment.

The uptake and inhibition assay-specific variables are given by

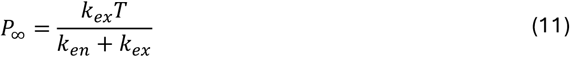

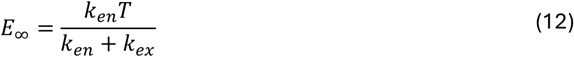

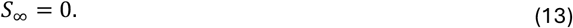

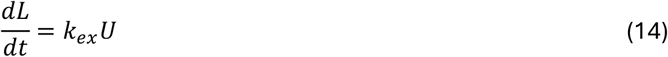

where *t* is time,

*L* is the amount of labeled GLUT4 in the uptake assay,

*U* is the amount of unlabeled GLUT4 in the uptake assay,

*P*_*LYi*_ is the amount of GLUT4 at the plasma membrane in the inhibition assay,

*E*_*LYi*_ is the amount of GLUT4 in the internal compartment in the inhibition assay, and

*F* is the reduction factor of LYi on the exocytosis rate.

Conservation of GLUT4 over the time scale of the experiments indicates that

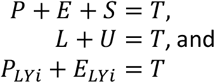

where *T* is the total amount of GLUT4 recycling (which may be insulin-dependent).

The steady states of the system Equations (8)-(10) are

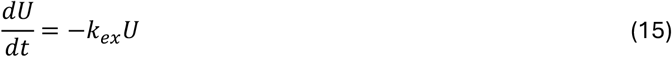

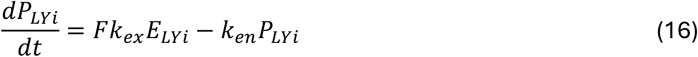

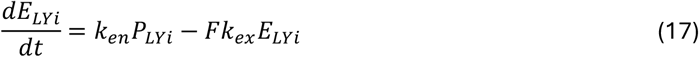

The initial conditions of the uptake and inhibition assays set the labeled level to the steady state level of GLUT4 at the plasma membrane for *i*nM of insulin, i.e.,

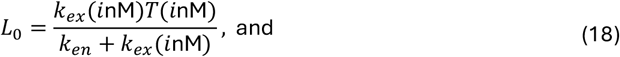

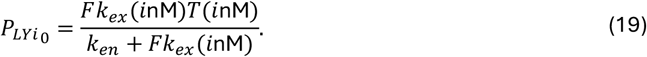

Depending on the hypothesis, any or all of the parameters of the system may be insulin or time dependent. As it was seen earlier that the goodness of fit to the models with insulin-independent endocytosis were comparable to those with insulin-dependent endocytosis rates, to constrain the number of parameters in the following investigations it is assumed from this point that the endocytosis rate, *k*_*en*_, is independent of insulin. It is also assumed that the total amount of GLUT4 recycling in the presence of the maximal insulin level *T* (100nM) = 1.

### Correspondence to the Experimental Data

The Transition assay tracks the evolution of the variable *P*, the amount at the plasma membrane, as the system evolves from the basal steady state to the stimulated steady state after the application of insulin, normalized to the steady state level in maximal insulin, i.e., *P* / *P*_∞_(100nM). The Uptake assay tracks the evolution of labeled GLUT4, again normalized to the level recycling in maximal insulin, *L*/ *T* (100nM). The inhibition assay tracks the amount of GLUT4 at the plasma membrane, normalized to the initial level in maximal insulin, *P*_*LYi*_ / *P*_∞_(100nM).

### Model Constraints

The Uptake assay is in a steady state with respect to insulin. The data for the different levels of insulin in the experiments constrains the exocytosis, *k*_*ex*_, and endocytosis, *k*_*en*_, rate constants and the total amount recycling *T* (if insulin-dependent), Table 1.

The Inhibition assay is also in an insulin steady state, and constrains the same parameters as the Uptake assay, as well as the inhibition factor, *F*, which is assumed here to be insulin independent.

The Transition assay tracks the system as it evolves from the basal to the insulin-stimulated steady state. These data also constrain the exocytosis and endocytosis rates, as well as the rates at which sequestered amounts, if any, are released to the recycling pathway, *k*_*Erel*_ and *k*_*prel*_ are constrained by this data.

Some model parameters are constrained by multiple data sets, linking the information from the different assays to test whether the modeling hypothesis is able to encode the observations across the different assay protocols.

### Modelling all the observations simultaneously

Can a single model explain the observations from all three assays? To explore some of the ways that we could explain the features of the data we consider some different hypotheses of insulin action below.

Here we are taking a parsimonious modeling approach to identify whether different hypotheses of insulin action could be explanatory for dominant processes in the system. Each model will then be fitted using the data from all three assays simultaneously to assess their efficacy in explaining the observations. Here, a single set of model parameters determine all the outputs of the model, corresponding to the Uptake, Transition, and Inhibition assay measurements.

### Model 0: Instant Rate Change, Instant Release

The uptake and inhibition assays are conducted in a steady state with respect to insulin. Thus, any sequestered GLUT4 is assumed to have already been released in these experiments. The Transition assay however tracks the transition between the basal and insulin-stimulated states.

If there is no sequestration *T* (*i*nM) = *T* (0nM) = 1. If *T* is insulin-dependent, the initial sequestered amount is *T* (*i*nM) − *T* (0nM) for insulin level *i*. For the instant release hypotheses, it is assumed that any additional GLUT4 is instantly added to the internal compartment upon the application of insulin. This effectively reduces the system to a two-compartment model

An alternate hypothesis, in which the sequestered GLUT4 was instantly added to the plasma membrane upon the application of insulin was also investigated but was found to fit more poorly (not shown). This is unsurprising as the surface amount of GLUT4 at 0 minutes did not greatly vary with respect to the applied level of insulin, see Figure 10 and *P*_0_ values in Table 4. It would be expected that the initial level would be a function of the applied insulin if GLUT4 was instantly released to the surface.

**Figure 10.**
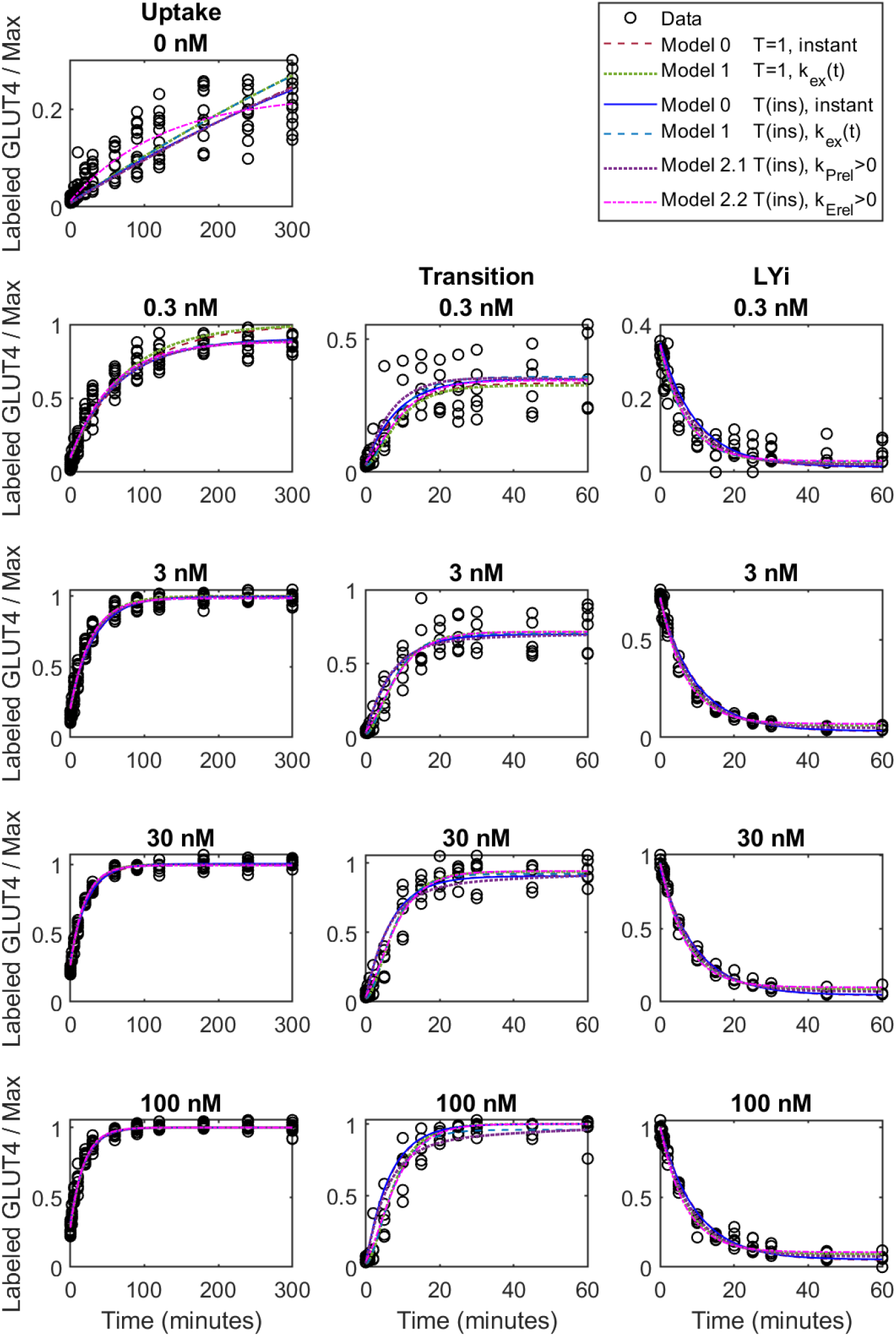
Assay data and models with all the data constraining the least squares fit as a function of the time of the data for each insulin concentration.

Thus, the initial conditions of the Transition assay are

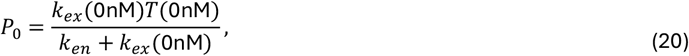

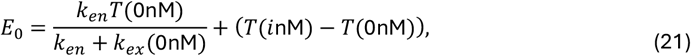

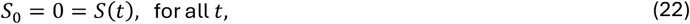

and the dynamics evolve according to Equations (8)-(10) with the insulin stimulated parameter values.

For the Uptake assay and the Inhibition assay, the system is in a steady state with respect to the given level of insulin, Equations (18) and (19) respectively.

This simplest model assumes that the time for the system to adjust to the application of insulin is negligible, i.e., the change in exocytosis rate is instantaneous compared to the observations of the system. Here it is assumed that the release of any sequestered GLUT4 is also instantaneous as described above. The system then evolves according to Equations (8)-(17) with all parameters set at their insulin-stimulated values. Two variants of this model were considered – one in which there was no sequestered GLUT4, i.e., *T* = 1, for all insulin levels, and one in which the total was insulin-dependent. This is effectively the model that was assumed in the characterization of the Uptake assay above, however in this case we constrain the model parameters using the data from all three assays.

### Model 1: Dynamic Rate Change, Instant Release

An alternate model retains the instantaneous release of any sequestered GLUT4 but allows the exocytosis rate in the Transition assay to dynamically change from the basal rate to that for the applied insulin dose, *i*nM.

If insulin induces a change in exocytosis, if this change occurs with a time constant *τ*, the exocytosis rate can be modelled as

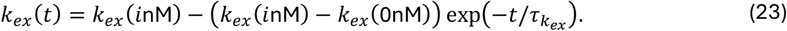

The dynamics and assumptions are otherwise identical to that of Model 0. The dynamically evolving exocytosis rate, Equation (23) can be implemented by the inclusion of an additional differential equation coupled to Equations (8)-(10),

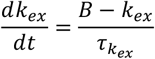

where *B* = *k*_*ex*_ (*i*nM) and initial condition, *k*_*ex*_ = *k*_*ex*_ (0nM).

Again, two variants of this model were considered – one in which there was no release, *T* = 1, for all insulin levels, and one in which the total was insulin-dependent.

### Model 2: Instant Rate Change, Dynamic Release

A third model was explored in which the insulin-stimulated change in the exocytic rate constant was instantaneous with respect to the transition, but allows the increase in the size of the cycling pool to dynamically change from the basal rate to that for the applied insulin dose, *i*nM

The initial amount in the sequestered store in the Transition assay,

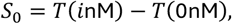

needs to enter the recycling pathway when *i*nM insulin is applied. Here we consider the two cases for the pathways shown in Figure 8. We assume here that the release of the sequestered GLUT4 is the only effect of the applied insulin, and that the exocytosis rate, *k*_*ex*_, instantaneously changes to its insulin-stimulated value.

#### 2.1: Instant Rate Change, Dynamic Release to the Plasma Membrane

Model 2.1 considers the dynamic release of the sequestered GLUT4 to the plasma membrane, i.e., *k*_*prel*_ > 0 and *k*_*Erel*_ = 0 in Equations (8)-(10).

#### 2.2: Instant Rate Change, Dynamic Release to the Internal Compartment

Model 2.2 considers the dynamic release of GLUT4 of the sequestered GLUT4 to the plasma membrane, i.e., *k*_*prel*_ = 0 and *k*_*Erel*_ > 0 in Equations (8)-(10).

### Testing the Hypotheses

The data for 0, 0.3, 3, 30, and 100nM insulin from the Uptake, Transition, and Inhibition assays was used to simultaneously constrain the parameters of each of the models, as these were the concentrations that were common in the data from all assays. The results of the fits to the data are shown in Figures 10 and 11. The residuals for each of the models are shown in Figure 12. Note that the Uptake assay explicitly includes data under 0nM insulin conditions, constraining the parameters for this insulin-level, whereas the Transition assays start in a 0nM insulin steady state and then evolve under the influence of the applied insulin level, and so also constrain the basal state parameters.

**Figure 11.**
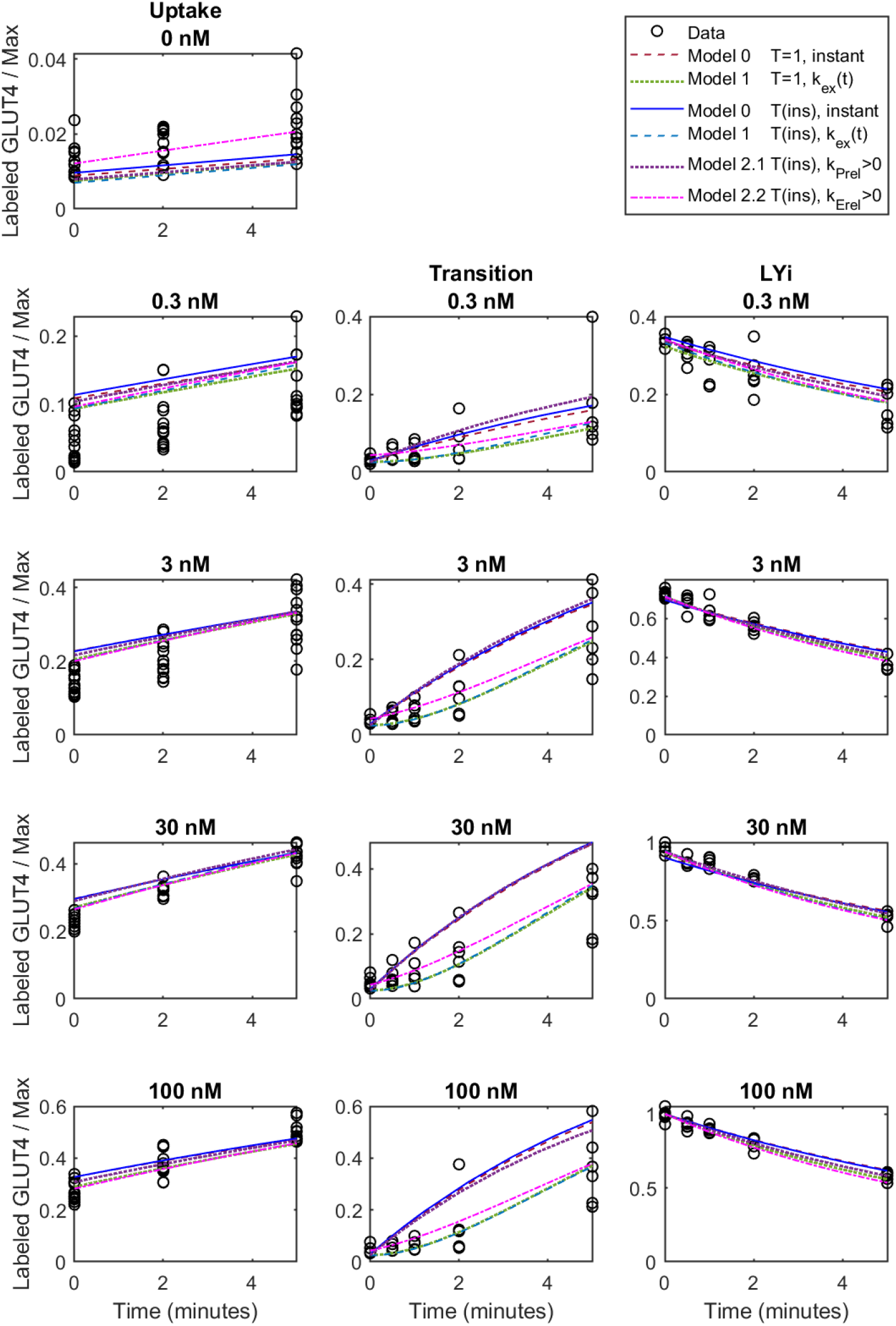
The first five minutes of the assay data and models (Figure 10) with all the data constraining the least squares fit as a function of the time of the data for each insulin concentration.

**Figure 12.**
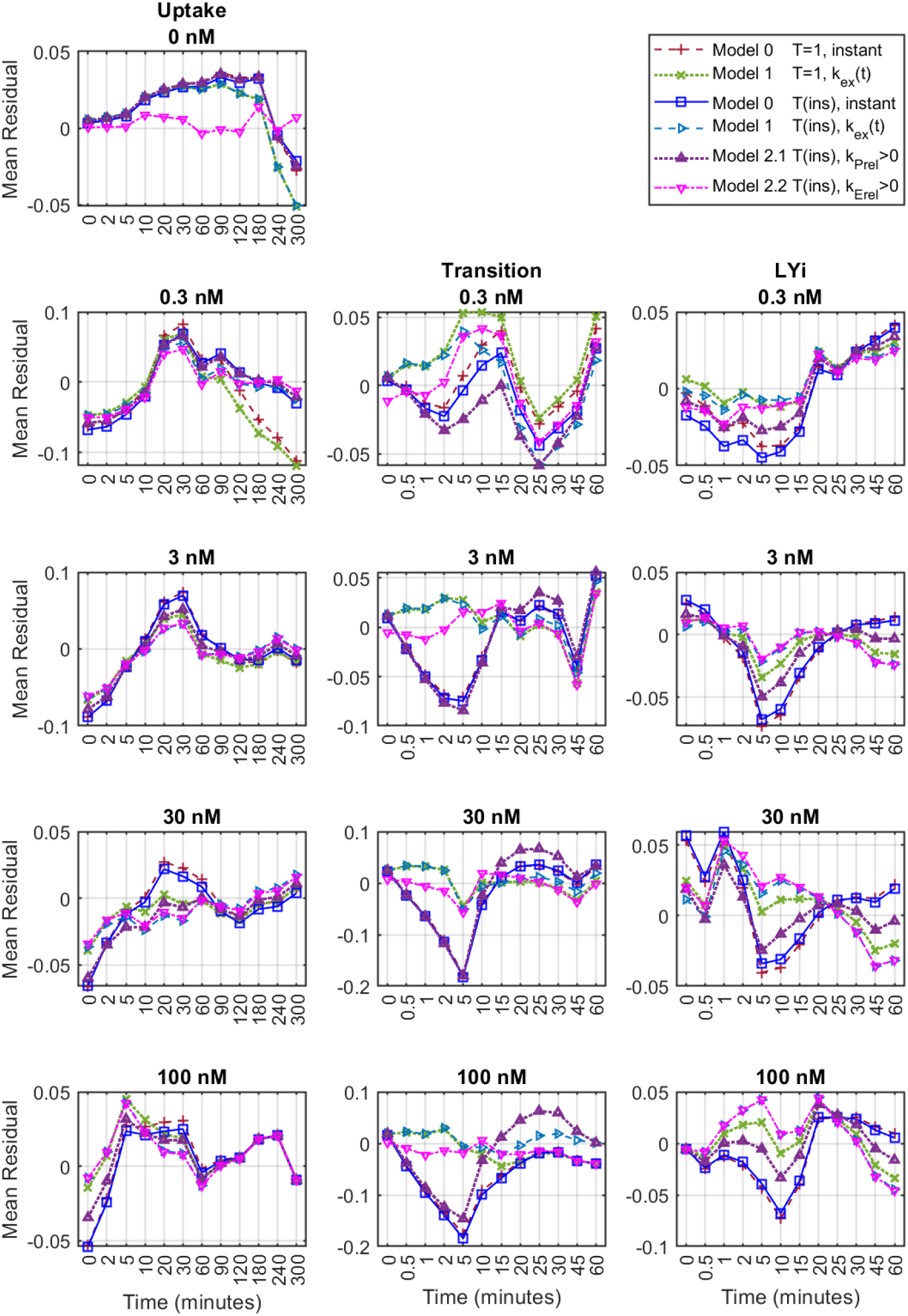
Mean Residuals at each time point (categorical axis) and concentration for the simultaneous fits to all three sets of assay data (Figure 10).

It can be seen in Figure 10 that the Inhibition data is similarly encoded by all the models at all concentrations. The uptake data for concentrations greater than 0.3nM is well described by all the model variants. The fits to the uptake data at 0 and 0.3nM and the transition data at early times for all concentrations have significant differences, see also the detail of the first five minutes of the assays in Figure 11.

Models 1 (for both T=1 and T(ins)) and 2.2 appear to capture the early time course of the transition data, however the residuals Model 1 (with T=1) for 0 and 0.3nM uptake are significantly worse than the others. Model 1 (with T(ins)) appears to fit the assays well, although the residuals at early times in the Transition assay are higher than those for Model 2.2.

The introduction of some sort of delay in the response of the system when insulin is applied (Models 1, 2.1, and 2.2) can be seen to help the correspondence of the models to the transition data. Releasing sequestered GLUT4 to the plasma membrane (or indeed instantaneously) does not appear to be as effective however, as this causes a greater deviation from the early transition data – particularly visible at higher concentrations, see Figure 12. Fixing the total recycling with respect to insulin causes a more rapid rise in the models compared to the uptake data in low insulin levels.

## Discussion

The characterization of the time courses for the Uptake, Transition, and Inhibition assays indicate that the overall dynamics of each has a single dominant time constant. It is clear from the least-squares fits that the overall exocytosis rate constant increases as a function of insulin, irrespective of whether it is assumed the total pool of GLUT4 is recycling or whether some is sequestered as a function of insulin. The endocytosis rate, on the other hand, is less variable as a function of insulin, and whilst is possible that this process could be insulin dependent, it is not necessary to assume this to recapitulate the data.

The minimum level of GLUT4 at the plasma membrane (in basal conditions) is approximately 1.5% of the maximum recycling level irrespective of the model hypothesis, and this ranges up to approximately 30% for maximal insulin (Model T(ins), Equation (1)), Table 2. These data are in accord with the understanding that insulin increases the surface expression of GLUT4.

Is GLUT4 sequestered at sub-maximal doses of insulin? We considered models in which the total recycling was either dependent or independent of insulin and used the Uptake assay data to constrain the model parameters. Although the overall Adjusted R^2^ of all the models were high, in Figures 1, 3, and 4, and Table 2, it can be seen that also allowing the plateau level in the assay output, the total amount of GLUT4 recycling, to be a function of insulin improves the match to the curvature of the time courses, which can also be seen in the residuals as a function of time. This was observed for all the individual replicate Uptake assay time courses, not just in the fits of the aggregated data from the 12 replicates at each concentration (Supplemental Figures S2a-i). Reducing the model degrees of freedom by constraining the endocytosis rate to be insulin-independent was not detrimental to the goodness of fit, thus indicating that this insulin dependence, if any, is a secondary effect, Figure 5 and Table 3.

The characterisation of the Transition assay, Figure 7 and Table 3, indicate that overall time constant for the transition time course ranges between 7.75 to 9.5 minutes. The plateau levels increased with insulin (noting that the normalisation of the assay was to the long-time level in maximum insulin). The half-time for the basal-insulin transition (5-6.5 minutes) is much faster than the half-time for the labelling of the GLUT4 in the Uptake assay (10-90 minutes) and does not change significantly with increasing concentrations of insulin.

The Inhibition assay showed an exponential decay to a low plateau, Figure 8, in line with the expectation that exocytosis is blocked, while endocytosis continues. The time constant of the response to the inhibitor, Table 5 was an increasing function of insulin, ranging between 5.6 to 8.1 minutes. The inferred inhibited plateau levels, *P*_∞_, indicated that the surface expression in the presence of the inhibitor and insulin, Table 5, ranged between 0.05 and 0.08 of the total GLUT4 recycling, commensurate with the basal levels.

To further probe which aspects of the system have dominant insulin dependence we considered the assays together, rather than independently. As each assay has a single dominant time constant, a simple three compartment model was employed, with two compartments representing the recycling from the internal cellular structures to the plasma membrane, and a third compartment allowing for the possible sequestration of GLUT4 as a function of insulin. We deliberately did not account for re-sequestration of GLUT4, as this is likely to be slow on the time scale of the Transition assay, and the Uptake and Inhibition assays are conducted in a steady state with respect to insulin.

The most parsimonious model considers the release of any sequestered GLUT4 to occur instantaneously when insulin is applied. Any release would largely only be observed in the Transition Assay, as the other assays are in a steady state with respect to insulin (although they constrain the model parameters). In this hypothesis the endocytosis rate constant is taken to be insulin-independent, and the exocytosis rate constant is insulin-dependent. The exocytosis rate constant was considered to either instantaneously change to its insulin-stimulated value upon the application of insulin (Model 0), or to change dynamically from the basal to the insulin-stimulated value (Model 1). In each case, models in which the total recycling pool was insulin-dependent and insulin-independent were considered. Models in which the exocytosis rate constants changed instantaneously, whilst the release of GLUT4 from sequestration changed dynamically were also considered (Model 2).

Model 0 was seen to capture most of the features of the steady dynamics. Similarly to the investigations using only the Uptake assay data, inspection of the residuals and the curvature of the outputs for the Uptake assay shows a better correspondence to the data at low insulin levels at the later time points if the total is allowed to be insulin-dependent, even allowing for the increased degrees of freedom in the model (Figures 10, 11, 12, Uptake; 0 and 0.3nM; dark blue and dark red).

The instant change hypothesis did not, however, capture the early time dynamics of the Transition assay data with or without sequestration (Figures 10, 11, 12, Transition; 3, 30 and 100 nM; dark blue and dark red). Different possibilities for the slower increase observed in the early times of the assay were considered.

Model 1 considered the dynamic change of the exocytosis rate as a function of time, allowing a smooth transition from the basal rate to the insulin-stimulated rate. The inclusion of a time-varying exocytosis rate only affects the Transition assay outputs, as the other assays are in a steady state with respect to insulin. Again, two variants were considered – insulin-independent and insulin-dependent totals, with any sequestered amounts being instantaneously transferred to the internal compartment of the recycling pathway upon the application of insulin. Adjusting for the inclusion of the additional parameter, it was found that the fit to the data was improved by the inclusion of a time-varying exocytosis rate with a time constant of approximately 0.3 minutes (Figures 10, 11, 12, Transition; 3, 30, and 100nM; green and aqua). As the dynamic rate did not affect the output of the other assays, any changes to the fits were largely driven by the better correspondence to the Transition assay data (although the other assays also constrained the parameters in the system). The residuals and shape of the outputs again indicate that the model with an insulin-dependent total appears to describe the Uptake assay data better than the model with insulin-independent totals (Figures 10, 11, 12, Uptake; 0 and 0.3nM; green and aqua).

Models 2.1 and 2.2 considered insulin-dependent totals in which the sequestered amounts were released at constant rates to the plasma membrane or internal structures respectively (Figures 10, 11, 12, Transition; 3, 30, and 100nM; purple and pink). Model 2.2 (dynamic release of sequestered GLUT4 to the internal compartment) best captures the trend of the data out of all the models considered here. Model 2.1 (dynamic release to the plasma membrane) was unable to capture the early dynamics of the transition assay.

It is possible that the slower increase in surface levels observed in the Transition assay could be due to the diffusion and uptake of insulin by the cells. However, this would be expected to be much faster than the time constants of the adjustment inferred from the modeling which were as small as 0.3 minutes in the case of a dynamic exocytosis rate and around 4.5 minutes for the dynamic release of sequestered GLUT4 to the internal compartment (which is approximately half the overall time constant of the Transition assay).

It is of course possible to consider that sequestered GLUT4 could be released to both internal membranes and the plasma membrane, and that the exocytosis rate could also be changing dynamically at the same time. However, allowing these further degrees of freedom also reduces the parameter identifiability. The parsimonious modeling here indicates that it is possible that the primary determinant of the observed dynamics of the system could be the sequestration of GLUT4 being released to internal structures in response to insulin.

## Conclusion

The characterization of the dose response of 3T3 L1 adipocytes here shows a graded response to increasing levels of insulin. This confirms the common understanding in the literature that increased insulin increases GLUT4 surface expression. Exocytosis was found to have an overall time constant between 16 min (maximum insulin) and >100 minutes for basal, whereas the overall endocytosis time constant was around 5-6 minutes.

The parsimonious modeling explored here suggests that the total GLUT4 recycling is insulin-dependent, although it appears there is little sequestration once the applied insulin is above 1nM. The models indicate that any sequestered GLUT4 is likely released to internal membrane structures and then recycled to the plasma membrane, accounting for the initial slower rate of increase in the plasma membrane levels when insulin is applied to the cells. This could involve trafficking of GLUT4 to a new compartment, or conversion of sequestered GLUT4 vesicles into cycling vesicles. An intriguing question that remains in this field is what is the mechanism to account for the dose dependent release of GLUT4 in response to submaximal concentrations of insulin? There is a rapid release or conversion of a population of GLUT4 vesicles (half time ∼3 minutes), then no further release over the course of these experiments despite the continuous presence of insulin (up to 6 hours). Incremental additions of increasing concentrations similarly release more vesicles into the cycling pool (5).

Furthermore, the nature of the dynamic conversion step (in exocytosis rate or in release) remains unknown. Future analysis of the effects of perturbations such as mutations in GLUT4 (2) or insulin-regulated proteins (e.g., AS160: (7,12), Rabs: (13,14)) on GLUT4 trafficking kinetics will help to resolve these questions. This paper shows the power of modeling to test hypotheses about the nature of these processes.

## Author Contributions

ACFC and CCM conceived and coordinated the study, analyzed the data, and prepared the manuscript.

ACFC and CCM designed, performed, and analyzed the mathematical models and simulations. IR designed, performed and analyzed the experiments.

All authors contributed to the writing of the manuscript, preparation of figures, reviewed the results, and approved the final version of the manuscript.

## References

1 Coster, A.C., Govers, R. and James, D.E. (2004) Insulin stimulates the entry of GLUT4 into the endosomal recycling pathway by a quantal mechanism. Traffic, 5, 763–771

2 Govers, R., Coster, A.C. and James, D.E. (2004) Insulin increases cell surface GLUT4 levels by dose dependently discharging GLUT4 into a cell surface recycling pathway. Mol. Cell. Biol., 24, 6456–6466

3 Klip, A., McGraw, T.E. and James, D.E. (2019) Thirty sweet years of GLUT4. J. Biol. Chem., 294, 11369–11381

4 Fazakerley, D.J., Koumanov, F., and Holman, G. D. (2022) GLUT4 On the move. Biochem. J., 479 (3), 445–462.

5 Burchfield, J.G., Lu, J., Fazakerley, D.J., Tan, S-X., Ng, Y., Mele, K., Buckley, M.J., Han, W., Hughes, W.E., and James, D.E. (2013) Novel Systems for Dynamically Assessing Insulin Action in Live Cells Reveals Heterogeneity in the Insulin Response, Traffic, 14, 259–273

6 Muretta, J.M., Romenskaia, I., and Mastick, C.C. (2008) Insulin releases GLUT4 from static storage compartments into cycling endosomes and increases the rate constant for GLUT4 exocytosis. J. Biol. Chem., 283, 311–323

7 Brewer, P.D., Romenskaia, I., Kanow, M.A., and Mastick, C.C. (2011) Loss of AS160 Akt substrate causes GLUT4 protein to accumulate in compartments that are primed for fusion in basal adipocytes. J. Biol. Chem., 286, 26287–26297

8 Martin, O. J., Lee, A., and McGraw, T. E. (2006) GLUT4 distribution between the plasma membrane and the intracellular compartments is maintained by an insulin-modulated bipartite dynamic mechanism. J. Biol. Chem., 281, 484–490

9 Dawson, K., Aviles-Hernandez, A., Cushman, S. W., and Malide, D. (2001) Insulin-regulated trafficking of dual-labeled glucose transporter 4 in primary rat adipose cells. Biochem. Biophys. Res. Commun., 287, 445–454

10 Lampson, M. A., Schmoranzer, J., Zeigerer, A., Simon, S. M., and McGraw, T. E. (2001) Insulin-regulated release from the endosomal recycling compartment is regulated by budding of specialized vesicles. Mol. Biol. Cell, 12, 3489–3501

11 Habtemichael, E. N., Brewer, P. D., Romenskaia, I., and Mastick, C. C. (2011) Kinetic evidence that GLUT4 follows different endocytic pathways than the receptors for transferrin and α2-macroglobulin. J. Biol. Chem., 286, 10115–10125

12 Brewer, P.D., Habtemichael, E.N., Romenskaia, I., Mastick, C.C. and Coster, A.C. (2014) Insulin-regulated GLUT4 translocation: membrane protein trafficking with six distinctive steps. J. Biol. Chem., 289, 17280–17298

13 Brewer, P.D., Habtemichael, E.N., Romenskaia, I., Coster, A.C. and Mastick, C.C. (2016) Rab14 limits the sorting of GLUT4 from endosomes into insulin-sensitive regulated secretory compartments in adipocytes. Biochem. J., 473, 1315–1327

14 Brewer, P.D., Habtemichael, E.N., Romenskaia, I., Mastick, C.C. and Coster, A.C. (2016) GLUT4 is sorted from a Rab10 GTPase-independent constitutive recycling pathway into a highly insulin-responsive Rab10 GTPase-dependent sequestration pathway after adipocyte differentiation. J. Biol. Chem., 291, 773–789

