## Supplementary Materials for "GLUT4 translocation with insulin: revisiting the case for dose-dependent quantal release"

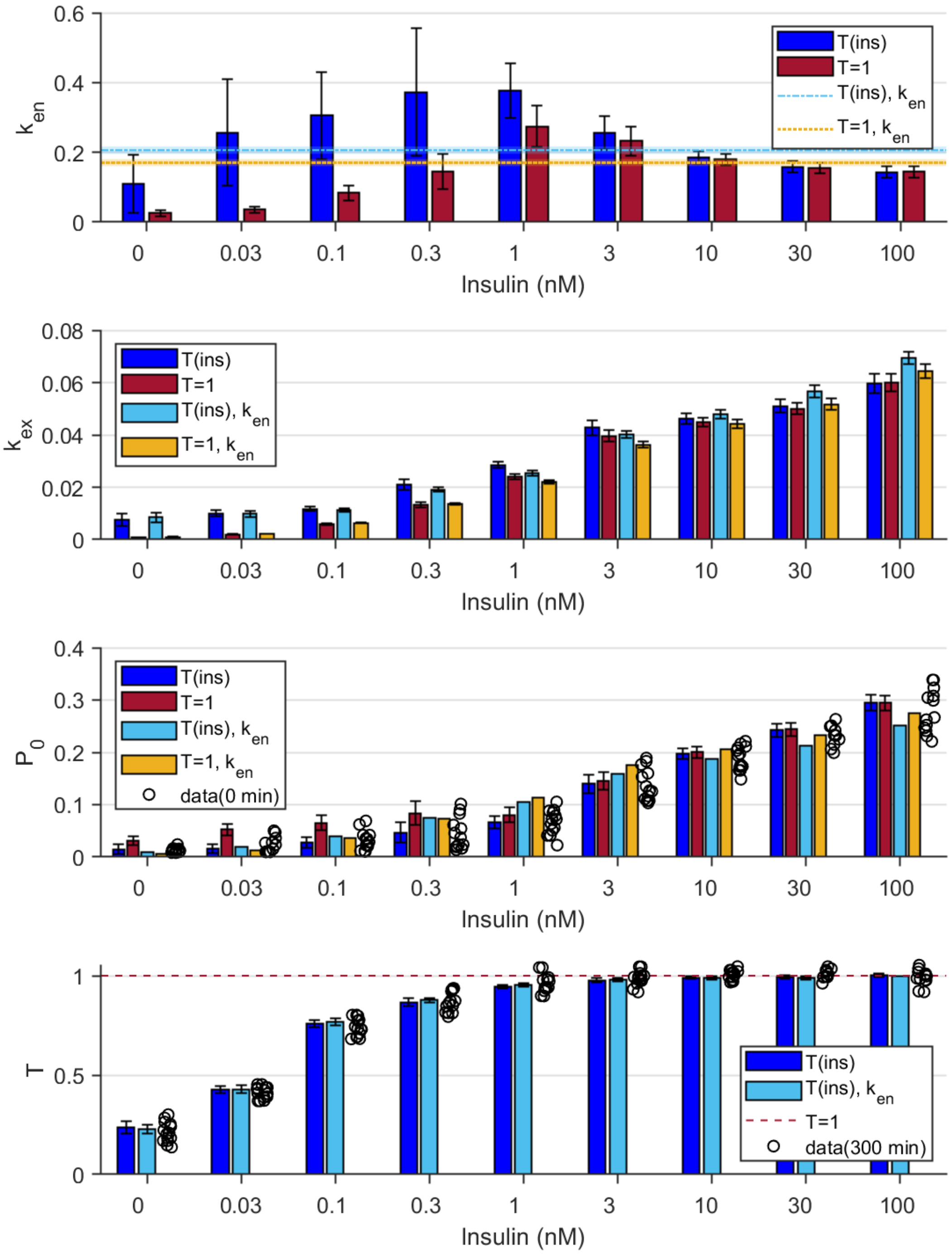


Figure S1. Parameter values of the least-squares fits of the Uptake Assay to the models as a function of insulin concentration (categorical axis). Models are T(ins) and T=1, Equations (1) and (2), and where the endocytosis rate is taken to be common across the concentrations, T(ins), $k_{en}$, and T=1, $k_{en}$, Equation (3). The error bars indicate the 95% confidence intervals for the parameter values. The initial plasma membrane level, $P_0$, inferred in each model is also compared to the uptake data at 0 minutes, when the surface GLUT4 is initially labeled. Similarly, the data at 300 minutes is shown in comparison to the inferred recycling total. The $k_{en}$ values for fits to Equation (3) are shown as a dotted line, with a shaded band showing the 95% confidence intervals.

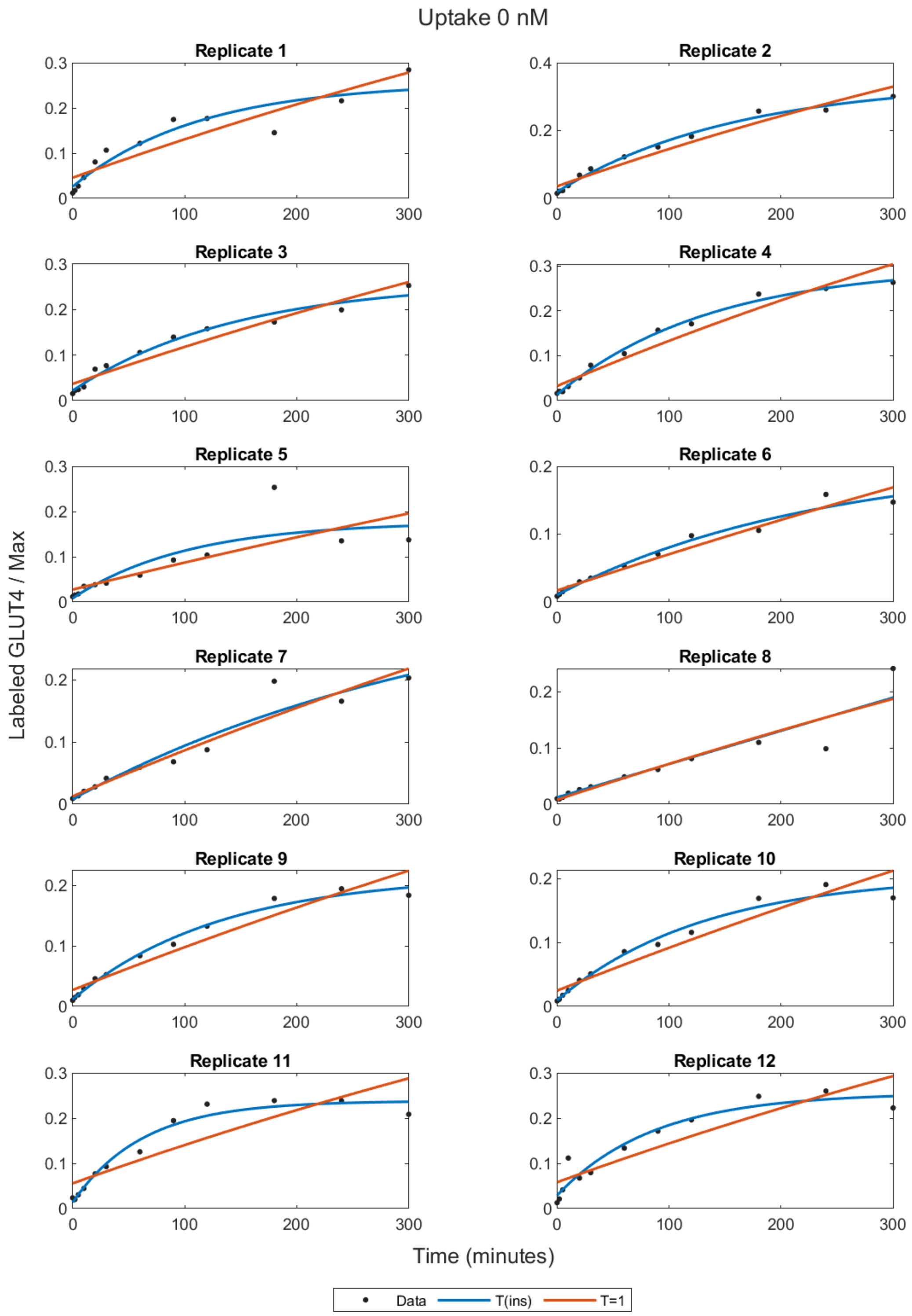


Figure S2a. Least-squares fits for the Uptake Assay to the T(ins) and T=1 models as a function of the time of the data for each replicate in the data set at 0nM insulin. Note that the inferred T value for the T(ins) fit of Replicate 8 was unphysically large (see Table S1). This fit was a result of the high variability of the data in the last two time points in this replicate. The confidence interval for T for this Replicate included zero, indicating that the estimate was unreliable. Similarly, the estimate for T for Replicate 7 was unreliable.

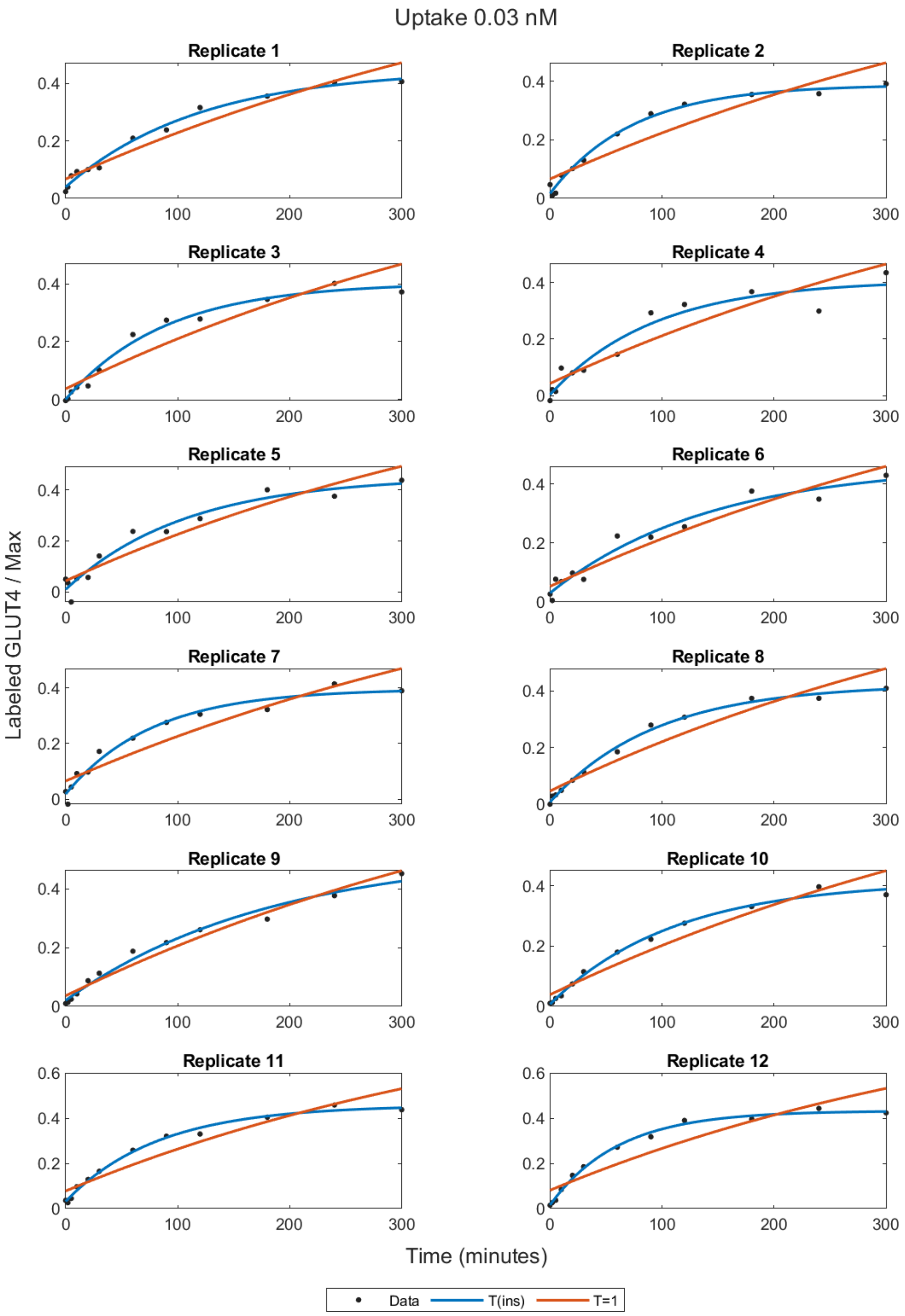


Figure S2b. Least-squares fits for the Uptake Assay to the T(ins) and T=1 models as a function of the time of the data for each replicate in the data set at 0.03nM insulin.

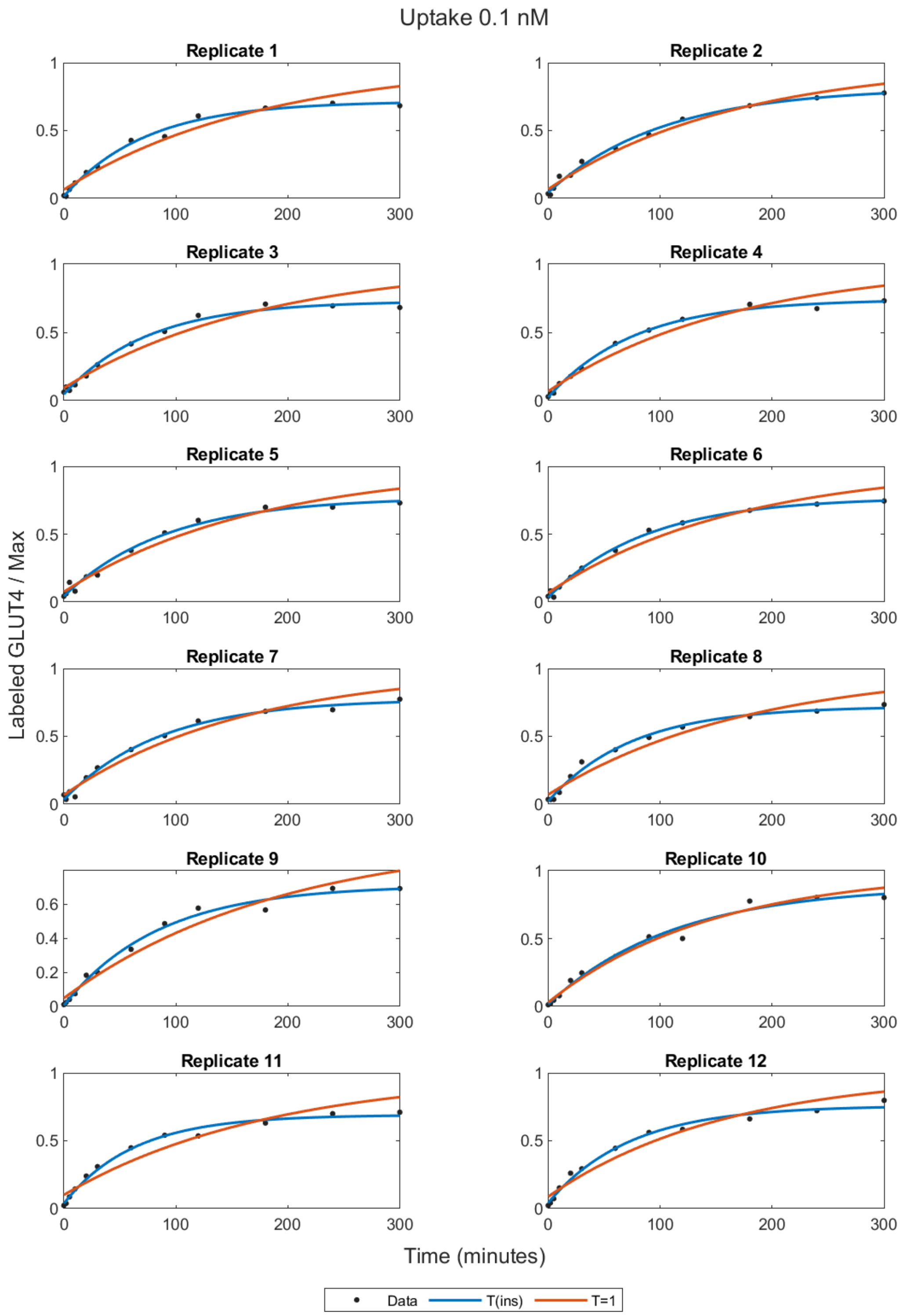


Figure S2c. Least-squares fits for the Uptake Assay to the T(ins) and T=1 models as a function of the time of the data for each replicate in the data set at 0.1nM insulin.

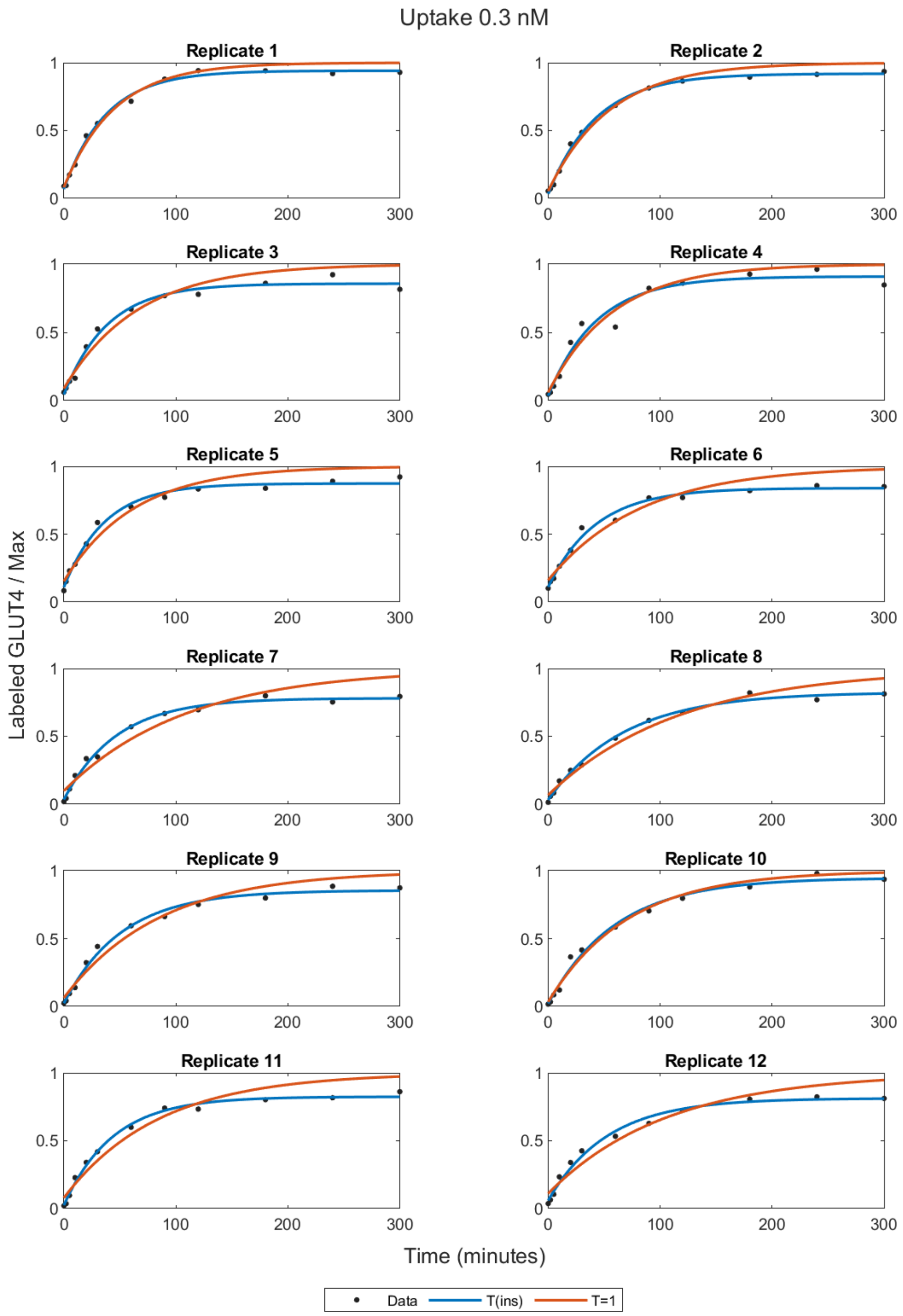


figure S2d. Least-squares fits for the Uptake Assay to the T(ins) and T=1 models as a function of the time of the data for each replicate in the data set at 0.3nM insulin.

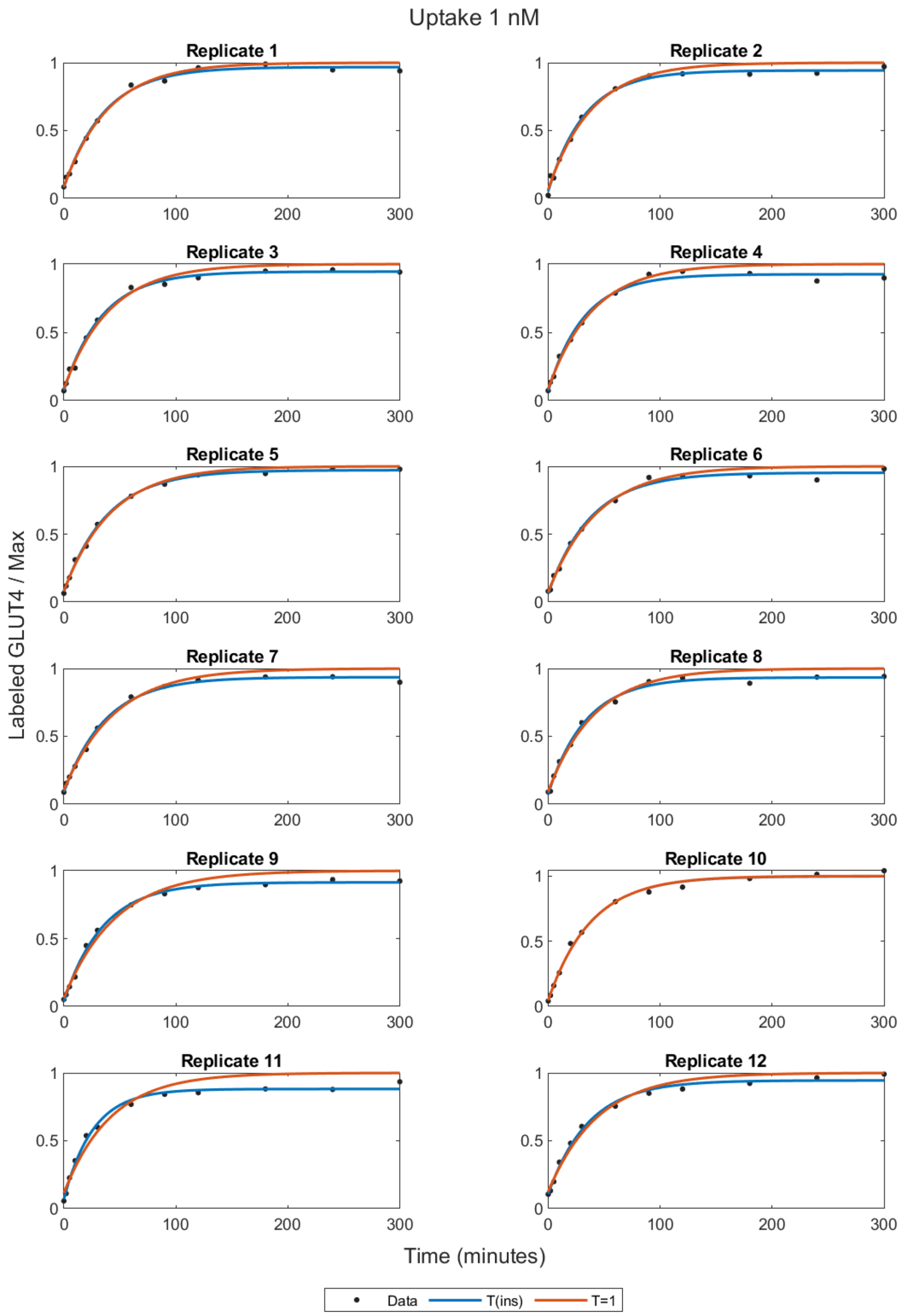


Figure S2e. Least-squares fits for the Uptake Assay to the T(ins) and T=1 models as a function of the time of the data for each replicate in the data set at 1nM insulin.

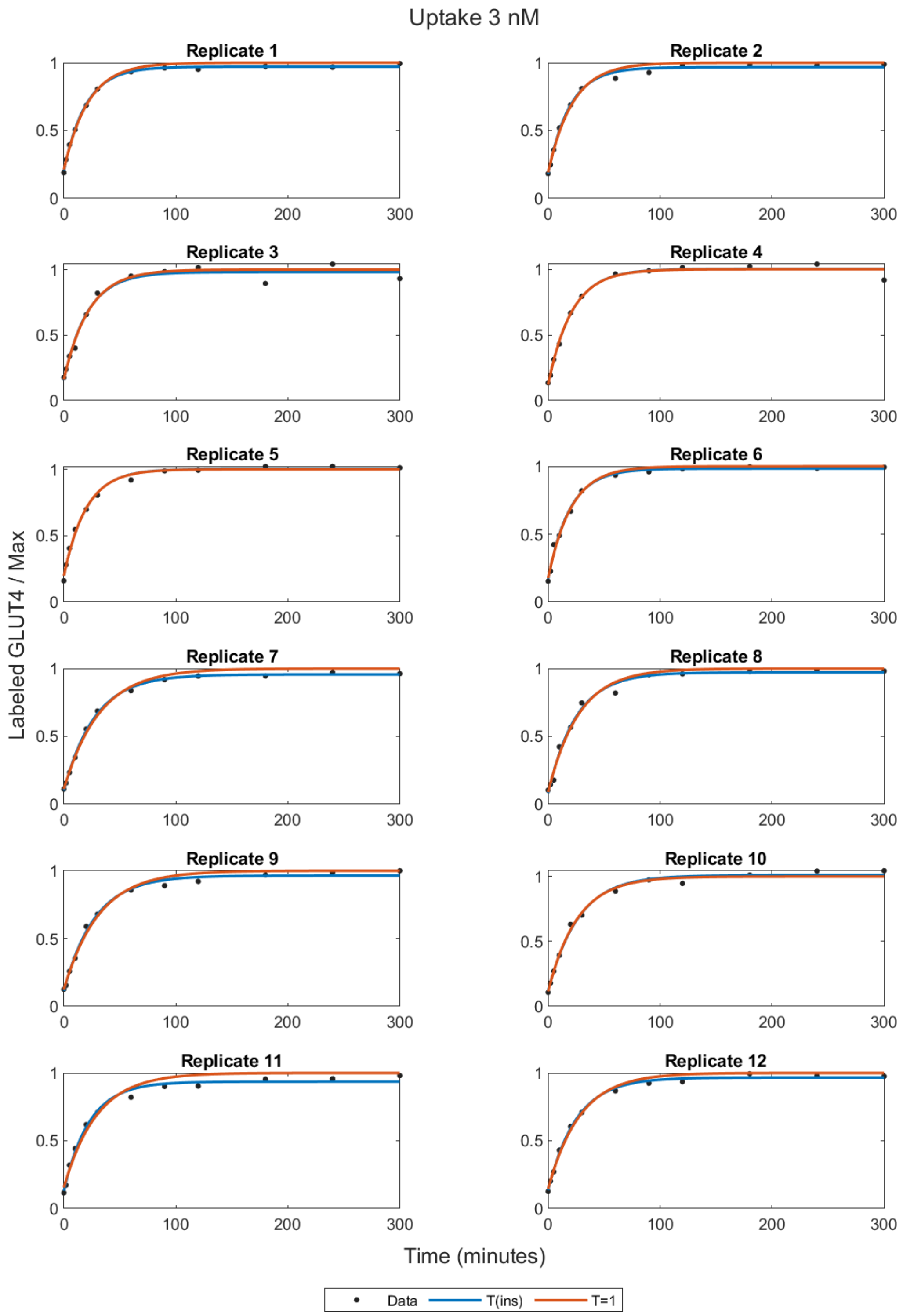


Figure S2f. Least-squares fits for the Uptake Assay to the T(ins) and T=1 models as a function of the time of the data for each replicate in the data set at 3nM insulin.

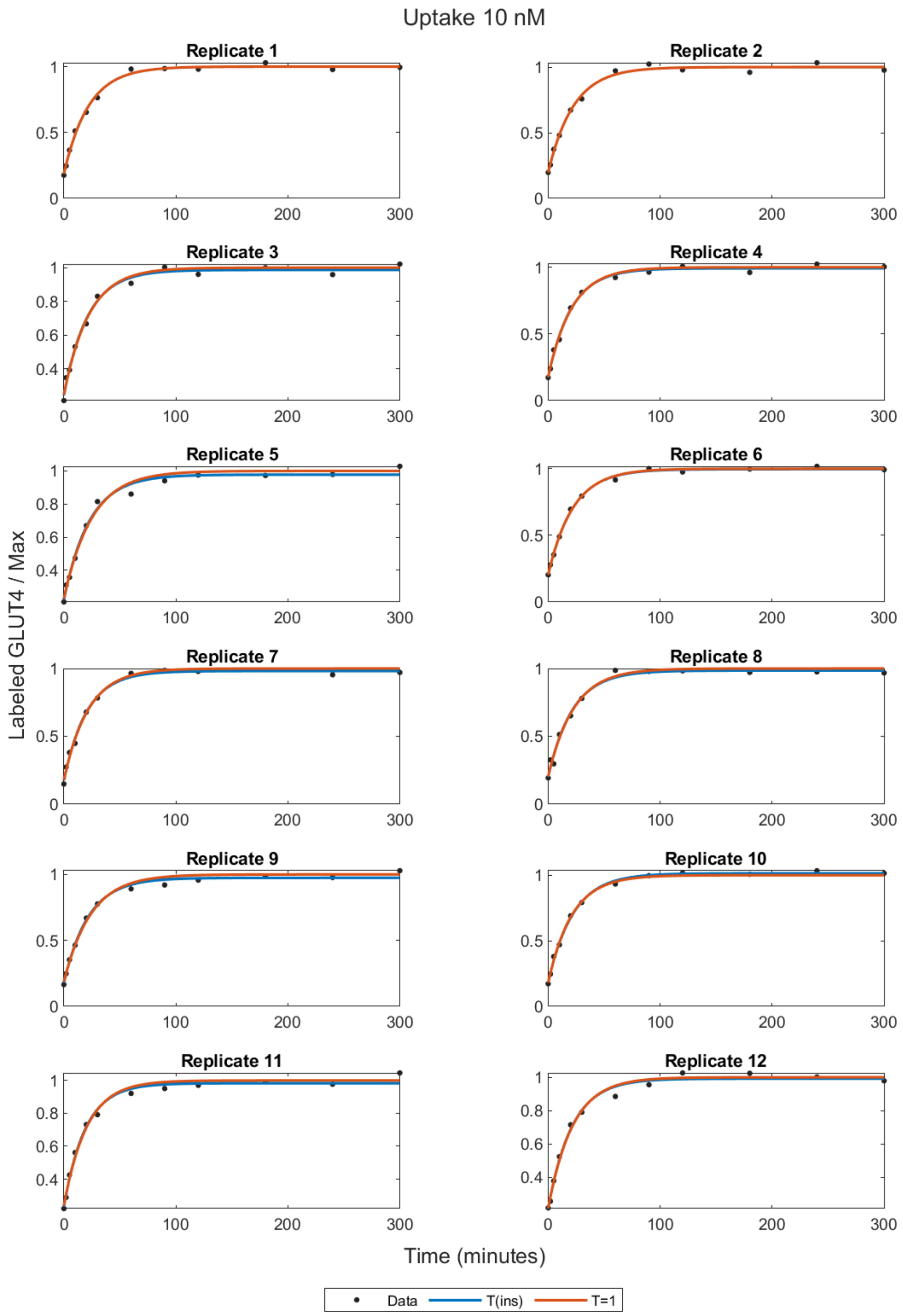


Figure S2g. Least-squares fits for the Uptake Assay to the T(ins) and T=1 models as a function of the time of the data for each replicate in the data set at 10nM insulin.

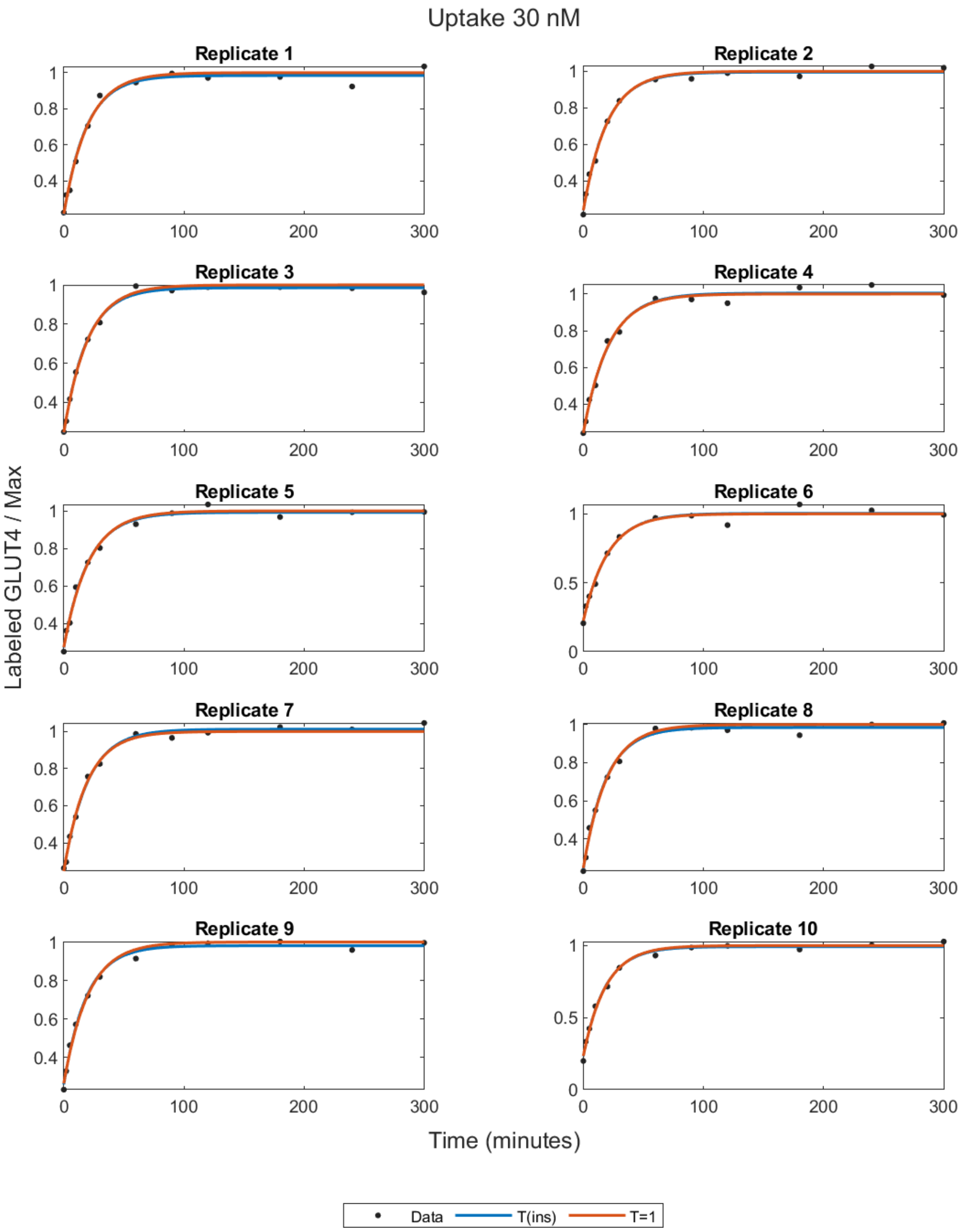


Figure S2h. Least-squares fits for the Uptake Assay to the T(ins) and T=1 models as a function of the time of the data for each replicate in the data set at 30nM insulin.

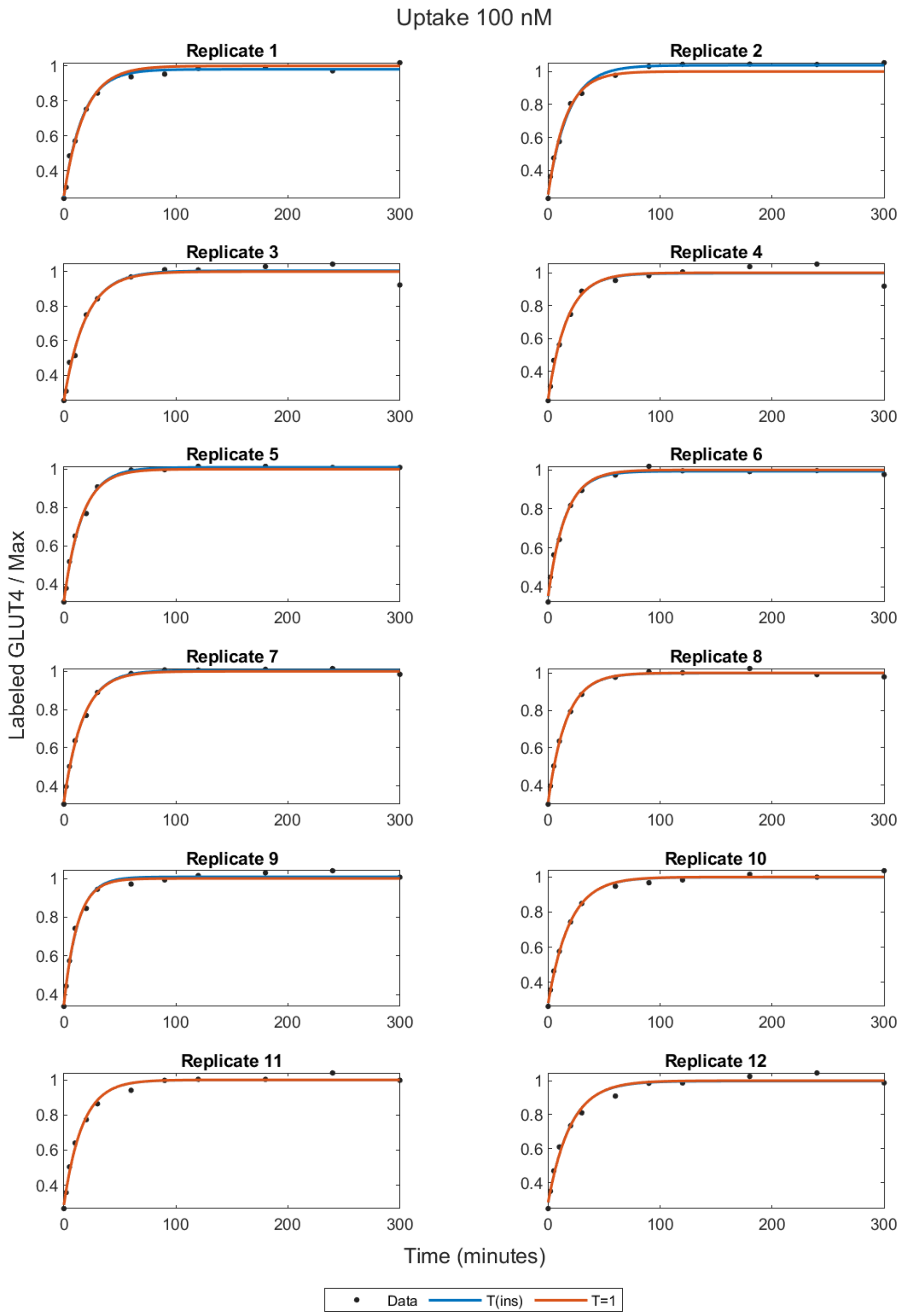


Figure S2i. Least-squares fits for the Uptake Assay to the T(ins) and T=1 models as a function of the time of the data for each replicate in the data set at 100nM insulin.

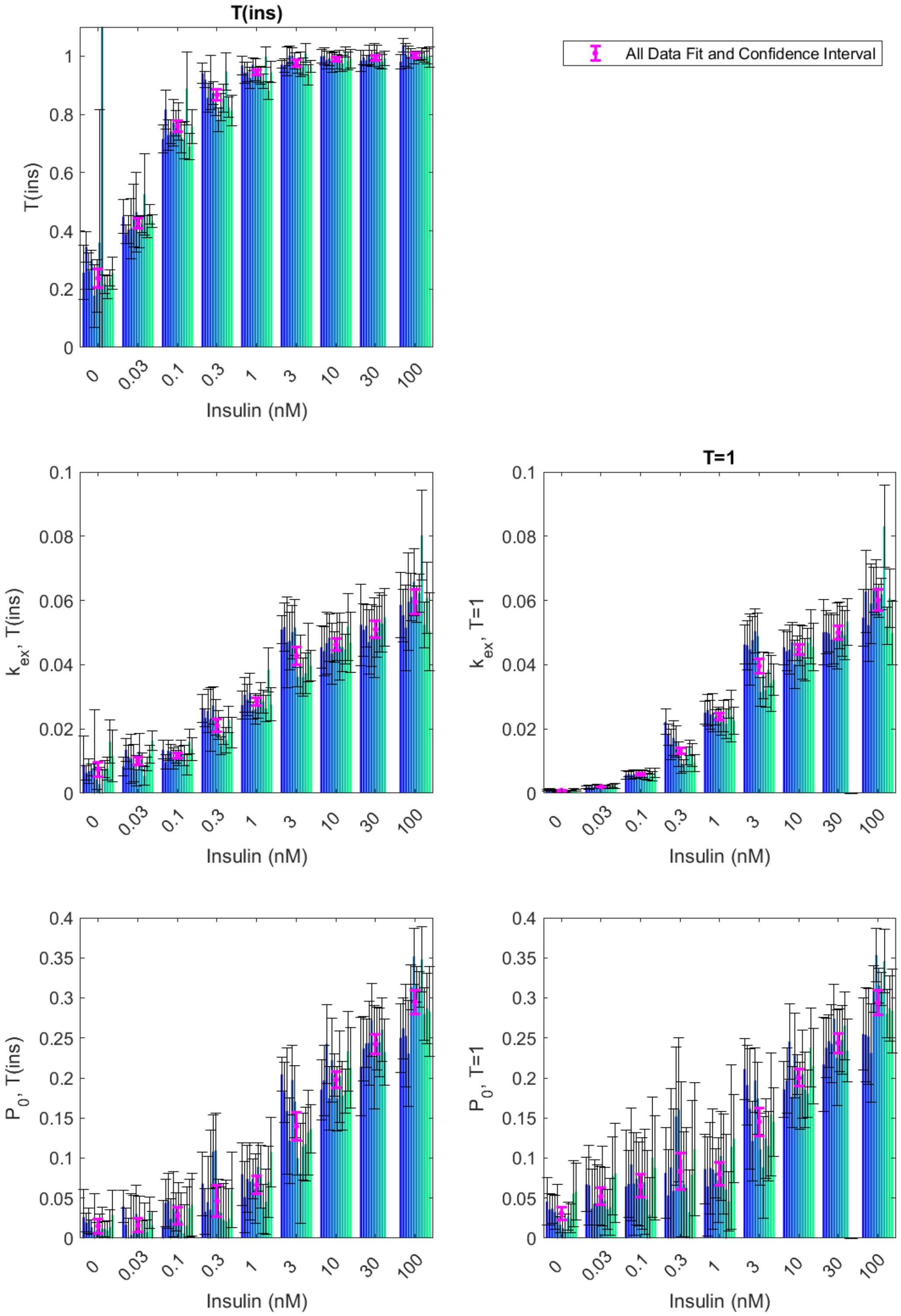


Figure S3. Parameter values of the least-squares fits of the Uptake Assay to the T(ins) and T=1 models for each of the replicate data sets as a function of insulin concentration (categorical axis). The parameter values for the fits to all the data are shown in magneta. The error bars indicate the 95% confidence intervals for the parameter values.

Table S1. Parameter values for the least-squares fits to the uptake data, Equation (1), with T=1. The 95% confidence interval is reported in brackets. The adjusted $R^2$ value (adj$R^2$) accounts for the number of degrees of freedom in the model.

| nM | Replicate | adjR2 | $k_{ex}$ | | $P_0$ | |
|---|---|---|---|---|---|---|
| 0 | 1 | 0.8493 | 0.0009 | (0.0007,0.0012) | 0.0458 | (0.0161,0.0755) |
| 0 | 2 | 0.9558 | 0.0012 | (0.0010,0.0014) | 0.0358 | (0.0163,0.0552) |
| 0 | 3 | 0.9347 | 0.0009 | (0.0007,0.0010) | 0.0368 | (0.0188,0.0548) |
| 0 | 4 | 0.9406 | 0.0011 | (0.0009,0.0013) | 0.0325 | (0.0116,0.0534) |
| 0 | 5 | 0.6088 | 0.0006 | (0.0003,0.0010) | 0.0274 | (-0.0119,0.0667) |
| 0 | 6 | 0.9536 | 0.0006 | (0.0005,0.0006) | 0.0168 | (0.0067,0.0269) |
| 0 | 7 | 0.9253 | 0.0008 | (0.0006,0.0009) | 0.0127 | (-0.0051,0.0304) |
| 0 | 8 | 0.8591 | 0.0007 | (0.0005,0.0009) | 0.0078 | (-0.0143,0.0298) |
| 0 | 9 | 0.9200 | 0.0008 | (0.0006,0.0009) | 0.0271 | (0.0096,0.0446) |
| 0 | 10 | 0.9091 | 0.0007 | (0.0006,0.0009) | 0.0246 | (0.0068,0.0425) |
| 0 | 11 | 0.7371 | 0.0009 | (0.0006,0.0013) | 0.0557 | (0.0145,0.0968) |
| 0 | 12 | 0.7964 | 0.0010 | (0.0006,0.0013) | 0.0582 | (0.0226,0.0938) |
| 0.03 | 1 | 0.9334 | 0.0019 | (0.0015,0.0023) | 0.0674 | (0.0335,0.1013) |
| 0.03 | 2 | 0.8591 | 0.0018 | (0.0013,0.0024) | 0.0661 | (0.0163,0.1159) |
| 0.03 | 3 | 0.8821 | 0.0020 | (0.0015,0.0025) | 0.0366 | (-0.0126,0.0859) |
| 0.03 | 4 | 0.8518 | 0.0019 | (0.0013,0.0025) | 0.0431 | (-0.0118,0.0981) |
| 0.03 | 5 | 0.8954 | 0.0021 | (0.0016,0.0027) | 0.0434 | (-0.0051,0.0919) |
| 0.03 | 6 | 0.9252 | 0.0019 | (0.0015,0.0023) | 0.0526 | (0.0161,0.0890) |
| 0.03 | 7 | 0.8650 | 0.0019 | (0.0013,0.0024) | 0.0653 | (0.0157,0.1149) |
| 0.03 | 8 | 0.9117 | 0.0020 | (0.0016,0.0025) | 0.0464 | (0.0041,0.0887) |
| 0.03 | 9 | 0.9687 | 0.0019 | (0.0017,0.0022) | 0.0359 | (0.0115,0.0603) |
| 0.03 | 10 | 0.9297 | 0.0019 | (0.0015,0.0022) | 0.0387 | (0.0033,0.0741) |
| 0.03 | 11 | 0.8964 | 0.0022 | (0.0017,0.0028) | 0.0776 | (0.0291,0.1261) |
| 0.03 | 12 | 0.8368 | 0.0022 | (0.0015,0.0030) | 0.0811 | (0.0191,0.1431) |
| 0.1 | 1 | 0.9354 | 0.0056 | (0.0043,0.0070) | 0.0646 | (-0.0056,0.1347) |
| 0.1 | 2 | 0.9787 | 0.0060 | (0.0051,0.0068) | 0.0675 | (0.0256,0.1095) |
| 0.1 | 3 | 0.9320 | 0.0057 | (0.0043,0.0071) | 0.0920 | (0.0219,0.1620) |
| 0.1 | 4 | 0.9484 | 0.0059 | (0.0046,0.0072) | 0.0680 | (0.0039,0.1320) |
| 0.1 | 5 | 0.9568 | 0.0058 | (0.0046,0.0069) | 0.0744 | (0.0167,0.1321) |
| 0.1 | 6 | 0.9657 | 0.0059 | (0.0049,0.0070) | 0.0674 | (0.0148,0.1199) |
| 0.1 | 7 | 0.9608 | 0.0061 | (0.0049,0.0072) | 0.0680 | (0.0110,0.1250) |
| 0.1 | 8 | 0.9429 | 0.0056 | (0.0043,0.0069) | 0.0703 | (0.0046,0.1360) |
| 0.1 | 9 | 0.9475 | 0.0051 | (0.0041,0.0062) | 0.0482 | (-0.0134,0.1097) |
| 0.1 | 10 | 0.9794 | 0.0068 | (0.0058,0.0078) | 0.0291 | (-0.0175,0.0758) |
| 0.1 | 11 | 0.9174 | 0.0054 | (0.0039,0.0069) | 0.1007 | (0.0256,0.1758) |
| 0.1 | 12 | 0.9495 | 0.0063 | (0.0049,0.0077) | 0.0878 | (0.0231,0.1525) |

*Table S1 continued overleaf*

Table S1 continued.
Parameter values for the least-squares fits to the uptake data, Equation (1), with T=1.

| nM | Replicate | adjR2 | $k_{ex}$ | | $P_0$ | |
|---|---|---|---|---|---|---|
| 0.3 | 1 | 0.9854 | 0.0222 | (0.0182,0.0262) | 0.0814 | (0.0253,0.1376) |
| 0.3 | 2 | 0.9835 | 0.0182 | (0.0149,0.0215) | 0.0533 | (-0.0045,0.1111) |
| 0.3 | 3 | 0.9388 | 0.0152 | (0.0103,0.0200) | 0.0884 | (-0.0101,0.1869) |
| 0.3 | 4 | 0.9472 | 0.0172 | (0.0117,0.0227) | 0.0590 | (-0.0430,0.1611) |
| 0.3 | 5 | 0.9483 | 0.0161 | (0.0112,0.0209) | 0.1519 | (0.0656,0.2381) |
| 0.3 | 6 | 0.9304 | 0.0121 | (0.0082,0.0160) | 0.1604 | (0.0704,0.2505) |
| 0.3 | 7 | 0.9139 | 0.0092 | (0.0062,0.0121) | 0.0983 | (0.0013,0.1953) |
| 0.3 | 8 | 0.9621 | 0.0085 | (0.0067,0.0103) | 0.0673 | (0.0018,0.1328) |
| 0.3 | 9 | 0.9610 | 0.0117 | (0.0089,0.0145) | 0.0626 | (-0.0131,0.1384) |
| 0.3 | 10 | 0.9855 | 0.0141 | (0.0118,0.0164) | 0.0324 | (-0.0204,0.0852) |
| 0.3 | 11 | 0.9383 | 0.0118 | (0.0083,0.0153) | 0.0792 | (-0.0130,0.1715) |
| 0.3 | 12 | 0.9385 | 0.0094 | (0.0068,0.0120) | 0.1111 | (0.0285,0.1937) |
| 1 | 1 | 0.9920 | 0.0251 | (0.0216,0.0285) | 0.0863 | (0.0428,0.1297) |
| 1 | 2 | 0.9842 | 0.0259 | (0.0209,0.0308) | 0.0645 | (0.0029,0.1261) |
| 1 | 3 | 0.9854 | 0.0245 | (0.0200,0.0290) | 0.0875 | (0.0303,0.1448) |
| 1 | 4 | 0.9740 | 0.0251 | (0.0191,0.0311) | 0.0859 | (0.0106,0.1612) |
| 1 | 5 | 0.9959 | 0.0237 | (0.0213,0.0260) | 0.0811 | (0.0500,0.1121) |
| 1 | 6 | 0.9876 | 0.0229 | (0.0190,0.0267) | 0.0722 | (0.0191,0.1253) |
| 1 | 7 | 0.9846 | 0.0221 | (0.0181,0.0262) | 0.1028 | (0.0471,0.1585) |
| 1 | 8 | 0.9817 | 0.0240 | (0.0192,0.0289) | 0.0928 | (0.0301,0.1555) |
| 1 | 9 | 0.9780 | 0.0216 | (0.0169,0.0262) | 0.0607 | (-0.0079,0.1293) |
| 1 | 10 | 0.9950 | 0.0262 | (0.0232,0.0292) | 0.0459 | (0.0087,0.0832) |
| 1 | 11 | 0.9442 | 0.0240 | (0.0159,0.0321) | 0.1141 | (0.0115,0.2166) |
| 1 | 12 | 0.9847 | 0.0226 | (0.0184,0.0268) | 0.1245 | (0.0692,0.1798) |
| 3 | 1 | 0.9934 | 0.0463 | (0.0402,0.0523) | 0.2111 | (0.1733,0.2488) |
| 3 | 2 | 0.9891 | 0.0461 | (0.0384,0.0538) | 0.1912 | (0.1416,0.2407) |
| 3 | 3 | 0.9776 | 0.0447 | (0.0337,0.0557) | 0.1615 | (0.0862,0.2368) |
| 3 | 4 | 0.9916 | 0.0476 | (0.0403,0.0550) | 0.1211 | (0.0713,0.1709) |
| 3 | 5 | 0.9934 | 0.0504 | (0.0436,0.0573) | 0.1970 | (0.1566,0.2374) |
| 3 | 6 | 0.9917 | 0.0489 | (0.0416,0.0561) | 0.1746 | (0.1292,0.2199) |
| 3 | 7 | 0.9916 | 0.0315 | (0.0270,0.0361) | 0.1109 | (0.0661,0.1558) |
| 3 | 8 | 0.9856 | 0.0367 | (0.0295,0.0438) | 0.0884 | (0.0254,0.1515) |
| 3 | 9 | 0.9897 | 0.0320 | (0.0268,0.0372) | 0.1245 | (0.0749,0.1740) |
| 3 | 10 | 0.9933 | 0.0385 | (0.0331,0.0438) | 0.1150 | (0.0712,0.1587) |
| 3 | 11 | 0.9727 | 0.0342 | (0.0254,0.0429) | 0.1543 | (0.0777,0.2310) |
| 3 | 12 | 0.9922 | 0.0354 | (0.0304,0.0404) | 0.1452 | (0.1026,0.1879) |

*Table S1 continued overleaf*

Table S1 continued.

Parameter values for the least-squares fits to the uptake data, Equation (1), with T=1.

| nM | Replicate | adjR2 | $k_{ex}$ | | $P_0$ | |
|---|---|---|---|---|---|---|
| 10 | 1 | 0.9951 | 0.0454 | (0.0401,0.0507) | 0.1857 | (0.1509,0.2205) |
| 10 | 2 | 0.9931 | 0.0442 | (0.0381,0.0503) | 0.1969 | (0.1562,0.2376) |
| 10 | 3 | 0.9890 | 0.0447 | (0.0370,0.0523) | 0.2455 | (0.1981,0.2929) |
| 10 | 4 | 0.9942 | 0.0472 | (0.0412,0.0531) | 0.1763 | (0.1383,0.2144) |
| 10 | 5 | 0.9864 | 0.0401 | (0.0325,0.0477) | 0.2281 | (0.1754,0.2808) |
| 10 | 6 | 0.9974 | 0.0450 | (0.0413,0.0488) | 0.2043 | (0.1798,0.2287) |
| 10 | 7 | 0.9927 | 0.0460 | (0.0396,0.0524) | 0.1775 | (0.1355,0.2194) |
| 10 | 8 | 0.9866 | 0.0438 | (0.0354,0.0521) | 0.2046 | (0.1496,0.2596) |
| 10 | 9 | 0.9902 | 0.0416 | (0.0349,0.0482) | 0.1852 | (0.1379,0.2325) |
| 10 | 10 | 0.9966 | 0.0468 | (0.0421,0.0514) | 0.1806 | (0.1506,0.2105) |
| 10 | 11 | 0.9886 | 0.0486 | (0.0401,0.0571) | 0.2380 | (0.1888,0.2872) |
| 10 | 12 | 0.9903 | 0.0456 | (0.0381,0.0530) | 0.2145 | (0.1675,0.2615) |
| 30 | 1 | 0.9847 | 0.0501 | (0.0399,0.0603) | 0.2167 | (0.1577,0.2757) |
| 30 | 2 | 0.9936 | 0.0501 | (0.0435,0.0568) | 0.2377 | (0.2003,0.2751) |
| 30 | 3 | 0.9949 | 0.0497 | (0.0440,0.0555) | 0.2459 | (0.2134,0.2784) |
| 30 | 4 | 0.9891 | 0.0478 | (0.0394,0.0562) | 0.2427 | (0.1937,0.2917) |
| 30 | 5 | 0.9906 | 0.0478 | (0.0402,0.0555) | 0.2741 | (0.2313,0.3169) |
| 30 | 6 | 0.9845 | 0.0490 | (0.0387,0.0593) | 0.2252 | (0.1651,0.2853) |
| 30 | 7 | 0.9932 | 0.0522 | (0.0449,0.0594) | 0.2472 | (0.2081,0.2864) |
| 30 | 8 | 0.9911 | 0.0505 | (0.0427,0.0583) | 0.2475 | (0.2045,0.2906) |
| 30 | 9 | 0.9908 | 0.0492 | (0.0415,0.0568) | 0.2653 | (0.2229,0.3076) |
| 30 | 10 | 0.9932 | 0.0534 | (0.0461,0.0607) | 0.2340 | (0.1949,0.2731) |
| 100 | 1 | 0.9901 | 0.0546 | (0.0458,0.0635) | 0.2549 | (0.2097,0.3002) |
| 100 | 2 | 0.9863 | 0.0627 | (0.0499,0.0756) | 0.2540 | (0.1955,0.3124) |
| 100 | 3 | 0.9839 | 0.0523 | (0.0411,0.0635) | 0.2516 | (0.1918,0.3114) |
| 100 | 4 | 0.9841 | 0.0590 | (0.0466,0.0714) | 0.2312 | (0.1699,0.2925) |
| 100 | 5 | 0.9957 | 0.0632 | (0.0562,0.0702) | 0.3114 | (0.2821,0.3406) |
| 100 | 6 | 0.9933 | 0.0641 | (0.0555,0.0727) | 0.3535 | (0.3202,0.3868) |
| 100 | 7 | 0.9975 | 0.0599 | (0.0549,0.0649) | 0.3156 | (0.2936,0.3375) |
| 100 | 8 | 0.9978 | 0.0620 | (0.0571,0.0668) | 0.3107 | (0.2901,0.3313) |
| 100 | 9 | 0.9916 | 0.0831 | (0.0702,0.0959) | 0.3457 | (0.3059,0.3855) |
| 100 | 10 | 0.9955 | 0.0521 | (0.0463,0.0579) | 0.2802 | (0.2502,0.3102) |
| 100 | 11 | 0.9918 | 0.0607 | (0.0515,0.0698) | 0.2871 | (0.2463,0.3279) |
| 100 | 12 | 0.9861 | 0.0498 | (0.0401,0.0596) | 0.2836 | (0.2314,0.3358) |

Table S2. Parameter values for the least-squares fits to the uptake data, Equation (1), with T a function of insulin. The 95% confidence interval is reported in brackets. The adjusted $R^2$ value (adj$R^2$) accounts for the number of degrees of freedom in the model.

| nM | Replicate | adjR2 | $k_{ex}$ | $P_0$ | $T$ |
|---|---|---|---|---|---|
| 0 | 1 | 0.8768 | 0.0088 (0.0000,0.0177) | 0.0260 (-0.0092,0.0613) | 0.2563 (0.1633,0.3493) |
| 0 | 2 | 0.9899 | 0.0063 (0.0042,0.0084) | 0.0193 (0.0075,0.0310) | 0.3449 (0.2912,0.3987) |
| 0 | 3 | 0.9697 | 0.0069 (0.0030,0.0107) | 0.0219 (0.0062,0.0376) | 0.2619 (0.1998,0.3239) |
| 0 | 4 | 0.9929 | 0.0073 (0.0054,0.0092) | 0.0137 (0.0044,0.0229) | 0.3008 (0.2676,0.3340) |
| 0 | 5 | 0.6879 | 0.0097 (-0.0065,0.0260) | 0.0082 (-0.0388,0.0552) | 0.1773 (0.0697,0.2850) |
| 0 | 6 | 0.9728 | 0.0043 (0.0011,0.0075) | 0.0103 (0.0007,0.0200) | 0.2113 (0.1218,0.3009) |
| 0 | 7 | 0.9248 | 0.0028 (-0.0024,0.0080) | 0.0081 (-0.0136,0.0298) | 0.3599 (-0.0968,0.8166) |
| 0 | 8 | 0.8509 | 0.0000 (-0.0072,0.0073) | 0.0119 (-0.0148,0.0386) | 13.5231[1] (-2204,2231) |
| 0 | 9 | 0.9859 | 0.0076 (0.0049,0.0102) | 0.0116 (0.0020,0.0211) | 0.2177 (0.1856,0.2498) |
| 0 | 10 | 0.9761 | 0.0077 (0.0041,0.0112) | 0.0095 (-0.0024,0.0214) | 0.2053 (0.1659,0.2446) |
| 0 | 11 | 0.9644 | 0.0160 (0.0093,0.0227) | 0.0138 (-0.0079,0.0354) | 0.2381 (0.2106,0.2656) |
| 0 | 12 | 0.9194 | 0.0116 (0.0035,0.0197) | 0.0292 (-0.0012,0.0595) | 0.2556 (0.2006,0.3106) |
| 0.03 | 1 | 0.9853 | 0.0083 (0.0055,0.0112) | 0.0396 (0.0194,0.0598) | 0.4490 (0.3910,0.5070) |
| 0.03 | 2 | 0.9862 | 0.0135 (0.0099,0.0171) | 0.0168 (-0.0042,0.0378) | 0.3883 (0.3566,0.4201) |
| 0.03 | 3 | 0.9822 | 0.0112 (0.0075,0.0150) | 0.0000 (-0.0251,0.0251) | 0.4043 (0.3571,0.4516) |
| 0.03 | 4 | 0.9217 | 0.0108 (0.0031,0.0184) | 0.0037 (-0.0484,0.0558) | 0.4083 (0.3044,0.5121) |
| 0.03 | 5 | 0.9471 | 0.0093 (0.0035,0.0150) | 0.0097 (-0.0343,0.0536) | 0.4524 (0.3449,0.5599) |
| 0.03 | 6 | 0.9532 | 0.0070 (0.0022,0.0119) | 0.0303 (-0.0057,0.0663) | 0.4645 (0.3279,0.6010) |
| 0.03 | 7 | 0.9651 | 0.0129 (0.0073,0.0186) | 0.0187 (-0.0151,0.0525) | 0.3957 (0.3421,0.4493) |
| 0.03 | 8 | 0.9939 | 0.0106 (0.0085,0.0126) | 0.0072 (-0.0072,0.0216) | 0.4235 (0.3942,0.4529) |
| 0.03 | 9 | 0.9809 | 0.0054 (0.0026,0.0082) | 0.0204 (-0.0029,0.0436) | 0.5261 (0.3868,0.6654) |
| 0.03 | 10 | 0.9924 | 0.0090 (0.0069,0.0111) | 0.0074 (-0.0075,0.0223) | 0.4156 (0.3773,0.4538) |
| 0.03 | 11 | 0.9904 | 0.0121 (0.0093,0.0149) | 0.0316 (0.0122,0.0510) | 0.4571 (0.4238,0.4904) |
| 0.03 | 12 | 0.9931 | 0.0164 (0.0134,0.0194) | 0.0160 (-0.0016,0.0336) | 0.4323 (0.4105,0.4542) |
| 0.1 | 1 | 0.9913 | 0.0136 (0.0107,0.0164) | 0.0151 (-0.0163,0.0466) | 0.7155 (0.6681,0.7629) |
| 0.1 | 2 | 0.9922 | 0.0097 (0.0075,0.0119) | 0.0439 (0.0145,0.0733) | 0.8168 (0.7492,0.8843) |
| 0.1 | 3 | 0.9889 | 0.0132 (0.0100,0.0165) | 0.0464 (0.0119,0.0810) | 0.7292 (0.6758,0.7825) |
| 0.1 | 4 | 0.9946 | 0.0130 (0.0108,0.0152) | 0.0240 (-0.0011,0.0491) | 0.7410 (0.7012,0.7808) |
| 0.1 | 5 | 0.9847 | 0.0109 (0.0076,0.0142) | 0.0425 (0.0019,0.0830) | 0.7714 (0.6921,0.8507) |
| 0.1 | 6 | 0.9950 | 0.0114 (0.0095,0.0133) | 0.0330 (0.0093,0.0567) | 0.7708 (0.7269,0.8147) |
| 0.1 | 7 | 0.9889 | 0.0116 (0.0087,0.0145) | 0.0331 (-0.0028,0.0690) | 0.7750 (0.7101,0.8400) |
| 0.1 | 8 | 0.9894 | 0.0134 (0.0102,0.0165) | 0.0217 (-0.0128,0.0562) | 0.7209 (0.6681,0.7737) |
| 0.1 | 9 | 0.9876 | 0.0117 (0.0086,0.0149) | 0.0059 (-0.0302,0.0421) | 0.7110 (0.6466,0.7755) |
| 0.1 | 10 | 0.9824 | 0.0089 (0.0057,0.0121) | 0.0156 (-0.0328,0.0641) | 0.8891 (0.7626,1.0155) |
| 0.1 | 11 | 0.9883 | 0.0160 (0.0122,0.0198) | 0.0376 (0.0018,0.0734) | 0.6902 (0.6446,0.7357) |
| 0.1 | 12 | 0.9870 | 0.0137 (0.0102,0.0173) | 0.0418 (0.0022,0.0813) | 0.7582 (0.6995,0.8169) |

*Table S2 continued overleaf*

[1] Note that this value is unphysically high. This was a result of the high variability of the data in the last two time points in this replicate, see Figure S2a. The confidence interval also reflects that this is an unreliable estimate for $T$.

Table S2 continued.

Parameter values for the least-squares fits to the uptake data, Equation (1), with T a function of insulin.

| nM | Replicate | adjR2 | $k_{ex}$ | $P_0$ | $T$ |
|---|---|---|---|---|---|
| 0.3 | 1 | 0.9937 | 0.0265 (0.0221,0.0308) | 0.0682 (0.0286,0.1078) | 0.9413 (0.9072,0.9753) |
| 0.3 | 2 | 0.9967 | 0.0234 (0.0206,0.0263) | 0.0330 (0.0046,0.0614) | 0.9195 (0.8930,0.9460) |
| 0.3 | 3 | 0.9845 | 0.0256 (0.0190,0.0323) | 0.0449 (-0.0127,0.1025) | 0.8567 (0.8061,0.9073) |
| 0.3 | 4 | 0.9593 | 0.0230 (0.0131,0.0329) | 0.0355 (-0.0635,0.1345) | 0.9092 (0.8155,1.0029) |
| 0.3 | 5 | 0.9891 | 0.0273 (0.0214,0.0332) | 0.1080 (0.0619,0.1541) | 0.8732 (0.8344,0.9121) |
| 0.3 | 6 | 0.9870 | 0.0236 (0.0180,0.0292) | 0.1095 (0.0630,0.1560) | 0.8391 (0.7959,0.8823) |
| 0.3 | 7 | 0.9905 | 0.0213 (0.0169,0.0257) | 0.0338 (-0.0060,0.0735) | 0.7808 (0.7410,0.8206) |
| 0.3 | 8 | 0.9929 | 0.0147 (0.0119,0.0175) | 0.0315 (-0.0015,0.0645) | 0.8250 (0.7793,0.8707) |
| 0.3 | 9 | 0.9896 | 0.0193 (0.0151,0.0235) | 0.0227 (-0.0225,0.0678) | 0.8550 (0.8062,0.9038) |
| 0.3 | 10 | 0.9877 | 0.0165 (0.0125,0.0205) | 0.0206 (-0.0318,0.0730) | 0.9471 (0.8822,1.0119) |
| 0.3 | 11 | 0.9928 | 0.0229 (0.0188,0.0270) | 0.0245 (-0.0132,0.0621) | 0.8244 (0.7887,0.8602) |
| 0.3 | 12 | 0.9860 | 0.0189 (0.0141,0.0237) | 0.0606 (0.0135,0.1076) | 0.8129 (0.7611,0.8647) |
| 1 | 1 | 0.9942 | 0.0274 (0.0231,0.0317) | 0.0799 (0.0410,0.1187) | 0.9679 (0.9353,1.0006) |
| 1 | 2 | 0.9928 | 0.0306 (0.0253,0.0360) | 0.0512 (0.0066,0.0958) | 0.9428 (0.9078,0.9779) |
| 1 | 3 | 0.9932 | 0.0291 (0.0242,0.0340) | 0.0740 (0.0323,0.1157) | 0.9448 (0.9110,0.9786) |
| 1 | 4 | 0.9910 | 0.0312 (0.0251,0.0372) | 0.0706 (0.0226,0.1185) | 0.9252 (0.8879,0.9626) |
| 1 | 5 | 0.9976 | 0.0258 (0.0232,0.0284) | 0.0744 (0.0496,0.0992) | 0.9714 (0.9497,0.9931) |
| 1 | 6 | 0.9926 | 0.0262 (0.0215,0.0309) | 0.0622 (0.0185,0.1058) | 0.9524 (0.9147,0.9902) |
| 1 | 7 | 0.9960 | 0.0269 (0.0234,0.0304) | 0.0890 (0.0586,0.1195) | 0.9357 (0.9097,0.9616) |
| 1 | 8 | 0.9939 | 0.0296 (0.0249,0.0343) | 0.0767 (0.0377,0.1157) | 0.9337 (0.9025,0.9650) |
| 1 | 9 | 0.9964 | 0.0285 (0.0250,0.0321) | 0.0370 (0.0063,0.0676) | 0.9142 (0.8891,0.9393) |
| 1 | 10 | 0.9944 | 0.0264 (0.0223,0.0305) | 0.0454 (0.0049,0.0860) | 0.9981 (0.9631,1.0330) |
| 1 | 11 | 0.9921 | 0.0384 (0.0315,0.0452) | 0.0703 (0.0261,0.1145) | 0.8814 (0.8506,0.9122) |
| 1 | 12 | 0.9920 | 0.0276 (0.0225,0.0328) | 0.1076 (0.0642,0.1509) | 0.9446 (0.9083,0.9808) |
| 3 | 1 | 0.9981 | 0.0510 (0.0467,0.0553) | 0.2045 (0.1833,0.2258) | 0.9707 (0.9576,0.9837) |
| 3 | 2 | 0.9943 | 0.0517 (0.0441,0.0592) | 0.1824 (0.1447,0.2201) | 0.9671 (0.9441,0.9900) |
| 3 | 3 | 0.9769 | 0.0470 (0.0330,0.0610) | 0.1586 (0.0796,0.2377) | 0.9815 (0.9315,1.0316) |
| 3 | 4 | 0.9906 | 0.0474 (0.0384,0.0564) | 0.1214 (0.0678,0.1751) | 1.0018 (0.9679,1.0356) |
| 3 | 5 | 0.9927 | 0.0501 (0.0418,0.0584) | 0.1975 (0.1540,0.2409) | 1.0017 (0.9749,1.0285) |
| 3 | 6 | 0.9925 | 0.0516 (0.0430,0.0603) | 0.1706 (0.1259,0.2153) | 0.9828 (0.9555,1.0100) |
| 3 | 7 | 0.9983 | 0.0362 (0.0332,0.0393) | 0.1000 (0.0784,0.1216) | 0.9559 (0.9404,0.9714) |
| 3 | 8 | 0.9871 | 0.0400 (0.0310,0.0491) | 0.0811 (0.0188,0.1433) | 0.9714 (0.9290,1.0139) |
| 3 | 9 | 0.9935 | 0.0361 (0.0303,0.0420) | 0.1145 (0.0730,0.1560) | 0.9641 (0.9343,0.9939) |
| 3 | 10 | 0.9929 | 0.0373 (0.0310,0.0436) | 0.1174 (0.0717,0.1632) | 1.0102 (0.9779,1.0425) |
| 3 | 11 | 0.9888 | 0.0438 (0.0347,0.0530) | 0.1320 (0.0791,0.1850) | 0.9356 (0.9010,0.9702) |
| 3 | 12 | 0.9965 | 0.0397 (0.0350,0.0443) | 0.1360 (0.1059,0.1662) | 0.9658 (0.9452,0.9865) |
| 10 | 1 | 0.9945 | 0.0455 (0.0389,0.0520) | 0.1856 (0.1481,0.2232) | 0.9997 (0.9756,1.0239) |
| 10 | 2 | 0.9923 | 0.0442 (0.0366,0.0518) | 0.1969 (0.1530,0.2408) | 0.9999 (0.9713,1.0285) |
| 10 | 3 | 0.9889 | 0.0467 (0.0371,0.0563) | 0.2426 (0.1933,0.2918) | 0.9868 (0.9555,1.0181) |
| 10 | 4 | 0.9939 | 0.0484 (0.0410,0.0557) | 0.1746 (0.1345,0.2147) | 0.9920 (0.9669,1.0171) |
| 10 | 5 | 0.9877 | 0.0435 (0.0340,0.0530) | 0.2221 (0.1700,0.2743) | 0.9776 (0.9434,1.0118) |
| 10 | 6 | 0.9972 | 0.0455 (0.0408,0.0502) | 0.2036 (0.1774,0.2298) | 0.9968 (0.9800,1.0136) |
| 10 | 7 | 0.9938 | 0.0486 (0.0412,0.0560) | 0.1740 (0.1341,0.2140) | 0.9815 (0.9565,1.0064) |
| 10 | 8 | 0.9865 | 0.0457 (0.0353,0.0562) | 0.2021 (0.1449,0.2592) | 0.9847 (0.9481,1.0214) |
| 10 | 9 | 0.9926 | 0.0455 (0.0378,0.0531) | 0.1782 (0.1354,0.2209) | 0.9740 (0.9465,1.0015) |
| 10 | 10 | 0.9972 | 0.0447 (0.0402,0.0493) | 0.1836 (0.1565,0.2108) | 1.0145 (0.9969,1.0321) |
| 10 | 11 | 0.9892 | 0.0518 (0.0414,0.0622) | 0.2333 (0.1838,0.2829) | 0.9829 (0.9527,1.0130) |
| 10 | 12 | 0.9895 | 0.0469 (0.0375,0.0563) | 0.2124 (0.1623,0.2626) | 0.9920 (0.9602,1.0238) |

*Table S2 continued overleaf*

Table S2 continued.

Parameter values for the least-squares fits to the uptake data, Equation (1), with T a function of insulin.

| nM | Replicate | adjR2 | $k_{ex}$ | $P_0$ | $T$ |
|---|---|---|---|---|---|
| 30 | 1 | 0.9845 | 0.0525 (0.0398,0.0651) | 0.2142 (0.1531,0.2753) | 0.9844 (0.9474,1.0214) |
| 30 | 2 | 0.9931 | 0.0509 (0.0427,0.0591) | 0.2368 (0.1967,0.2769) | 0.9957 (0.9711,1.0202) |
| 30 | 3 | 0.9958 | 0.0521 (0.0456,0.0586) | 0.2432 (0.2129,0.2736) | 0.9854 (0.9670,1.0038) |
| 30 | 4 | 0.9880 | 0.0470 (0.0370,0.0571) | 0.2437 (0.1913,0.2960) | 1.0046 (0.9715,1.0378) |
| 30 | 5 | 0.9900 | 0.0491 (0.0396,0.0587) | 0.2724 (0.2268,0.3179) | 0.9925 (0.9642,1.0208) |
| 30 | 6 | 0.9829 | 0.0484 (0.0360,0.0608) | 0.2259 (0.1615,0.2904) | 1.0038 (0.9634,1.0441) |
| 30 | 7 | 0.9934 | 0.0502 (0.0423,0.0581) | 0.2496 (0.2105,0.2888) | 1.0121 (0.9879,1.0362) |
| 30 | 8 | 0.9918 | 0.0532 (0.0439,0.0625) | 0.2443 (0.2016,0.2870) | 0.9845 (0.9588,1.0102) |
| 30 | 9 | 0.9924 | 0.0528 (0.0439,0.0617) | 0.2604 (0.2204,0.3005) | 0.9809 (0.9568,1.0051) |
| 30 | 10 | 0.9927 | 0.0548 (0.0458,0.0637) | 0.2323 (0.1909,0.2737) | 0.9928 (0.9682,1.0174) |
| 100 | 1 | 0.9918 | 0.0586 (0.0484,0.0688) | 0.2501 (0.2074,0.2929) | 0.9805 (0.9558,1.0053) |
| 100 | 2 | 0.9941 | 0.0554 (0.0472,0.0635) | 0.2621 (0.2240,0.3002) | 1.0377 (1.0152,1.0603) |
| 100 | 3 | 0.9822 | 0.0515 (0.0382,0.0649) | 0.2525 (0.1885,0.3165) | 1.0048 (0.9658,1.0438) |
| 100 | 4 | 0.9823 | 0.0596 (0.0444,0.0748) | 0.2306 (0.1645,0.2966) | 0.9972 (0.9591,1.0352) |
| 100 | 5 | 0.9960 | 0.0611 (0.0537,0.0684) | 0.3133 (0.2848,0.3418) | 1.0098 (0.9935,1.0260) |
| 100 | 6 | 0.9930 | 0.0658 (0.0554,0.0763) | 0.3520 (0.3171,0.3869) | 0.9934 (0.9740,1.0128) |
| 100 | 7 | 0.9974 | 0.0587 (0.0531,0.0644) | 0.3166 (0.2943,0.3390) | 1.0055 (0.9925,1.0185) |
| 100 | 8 | 0.9975 | 0.0623 (0.0564,0.0682) | 0.3104 (0.2882,0.3326) | 0.9987 (0.9861,1.0112) |
| 100 | 9 | 0.9914 | 0.0803 (0.0662,0.0943) | 0.3478 (0.3071,0.3886) | 1.0083 (0.9873,1.0294) |
| 100 | 10 | 0.9950 | 0.0524 (0.0452,0.0595) | 0.2799 (0.2476,0.3122) | 0.9987 (0.9792,1.0183) |
| 100 | 11 | 0.9909 | 0.0608 (0.0498,0.0719) | 0.2869 (0.2429,0.3308) | 0.9993 (0.9742,1.0244) |
| 100 | 12 | 0.9846 | 0.0504 (0.0382,0.0625) | 0.2828 (0.2265,0.3391) | 0.9974 (0.9628,1.0320) |

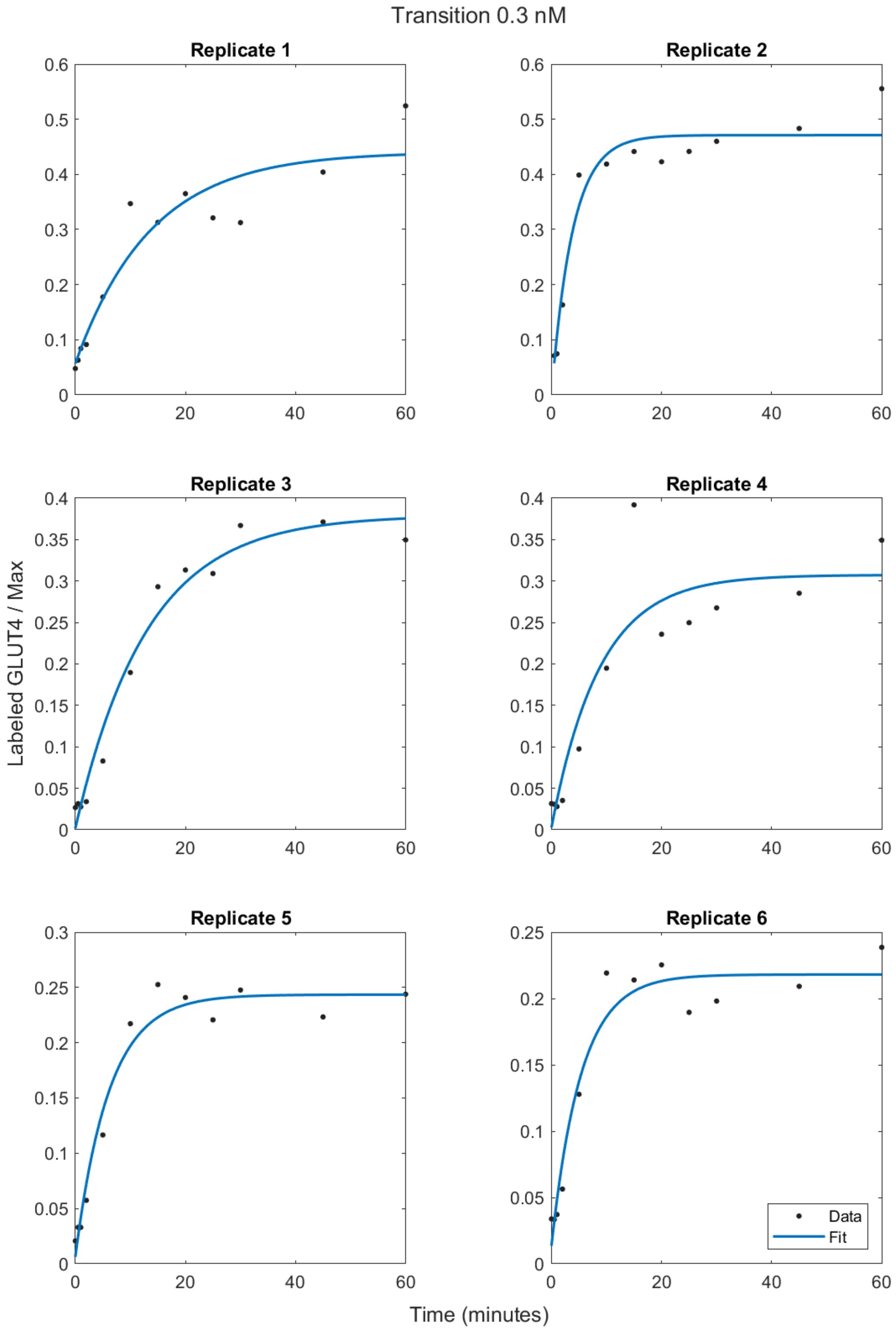


Figure S4a. Least-squares fits to Equation (4) for the Transition Assay as a function of the time of the data for each replicate in the data set as 0.3nM insulin was applied.

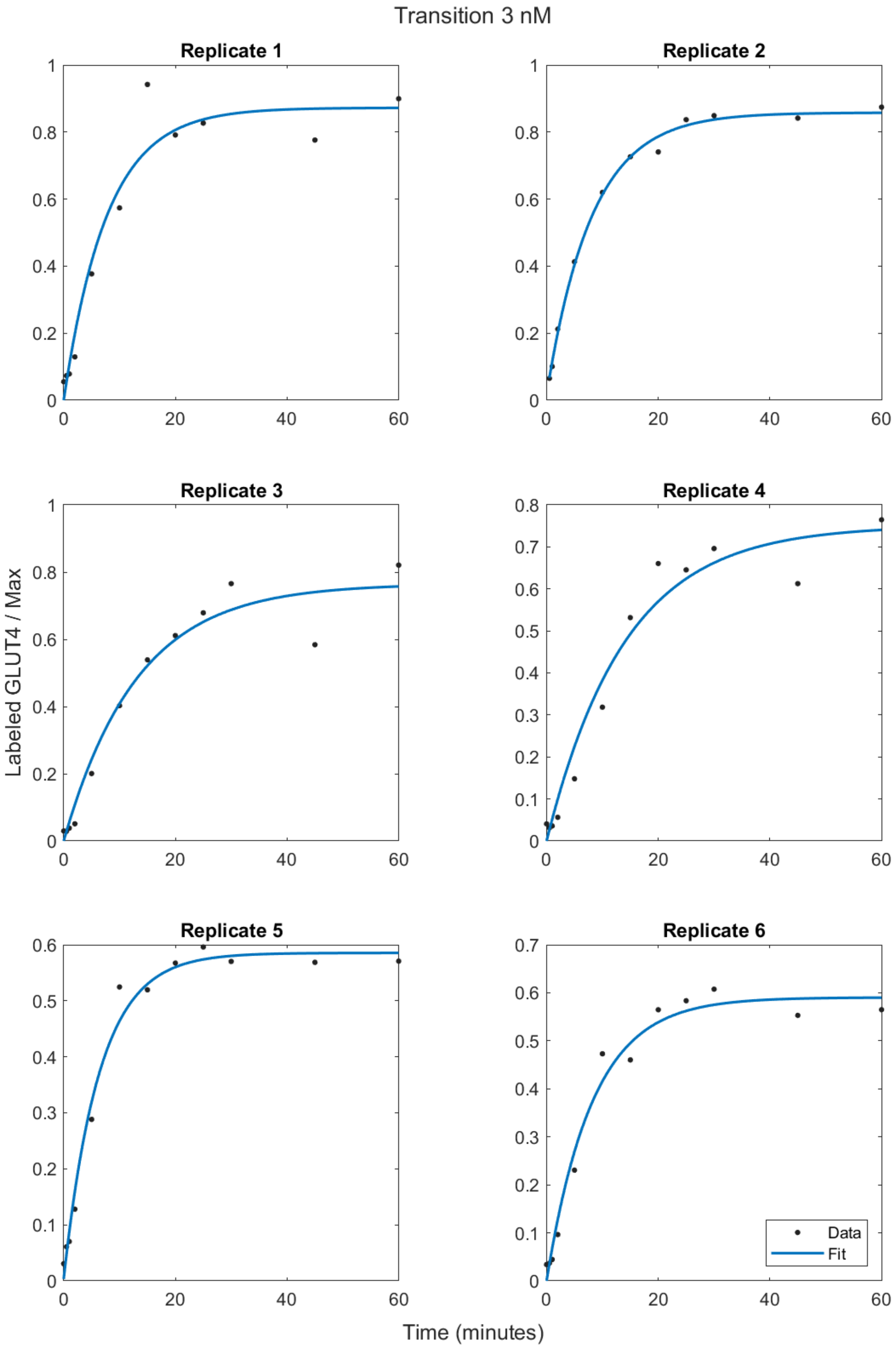


Figure S4b. Least-squares fits to Equation (4) for the Transition Assay as a function of the time of the data for each replicate in the data set as 3nM insulin was applied.

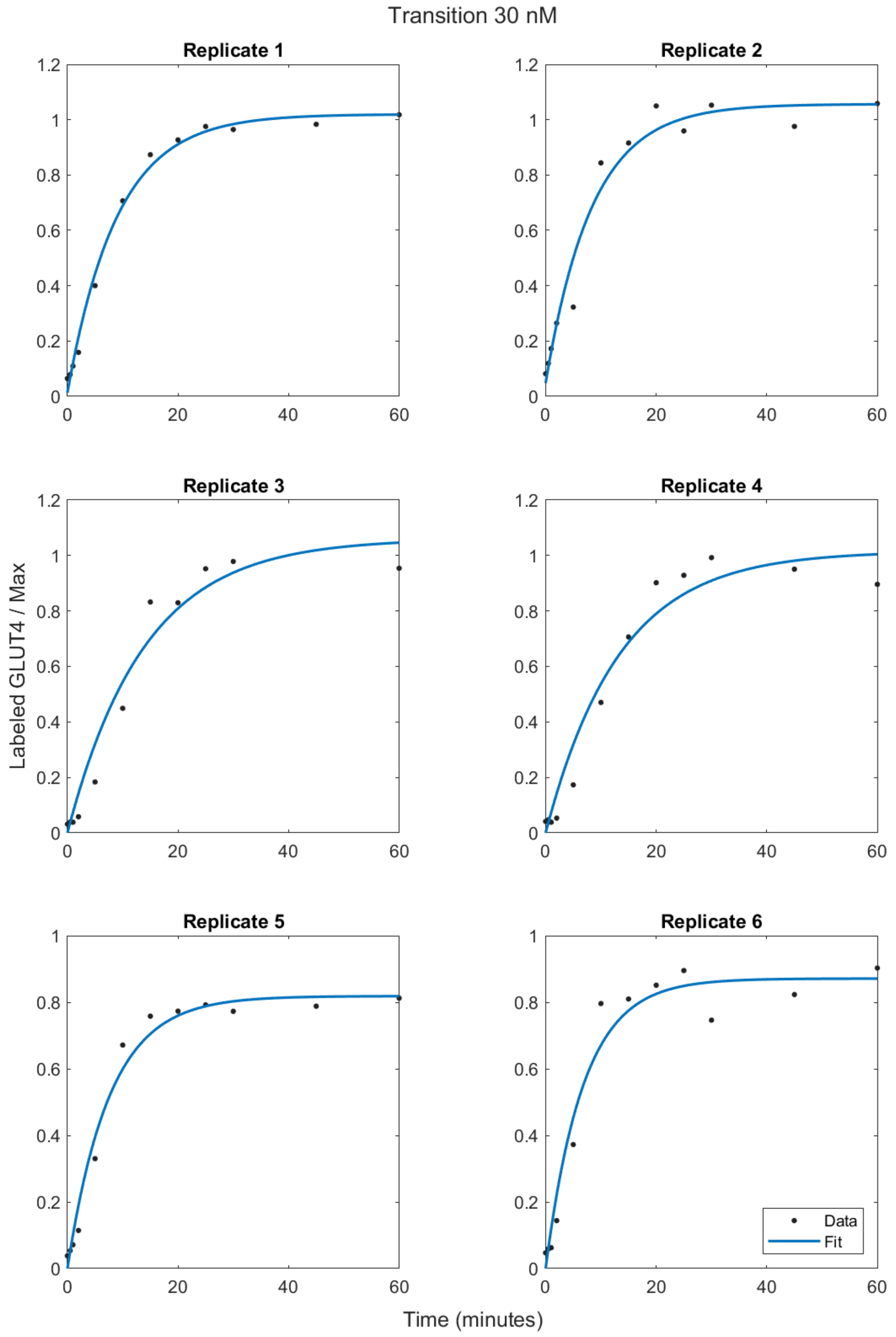


Figure S4c. Least-squares fits to Equation (4) for the Transition Assay as a function of the time of the data for each replicate in the data set as 30nM insulin was applied.

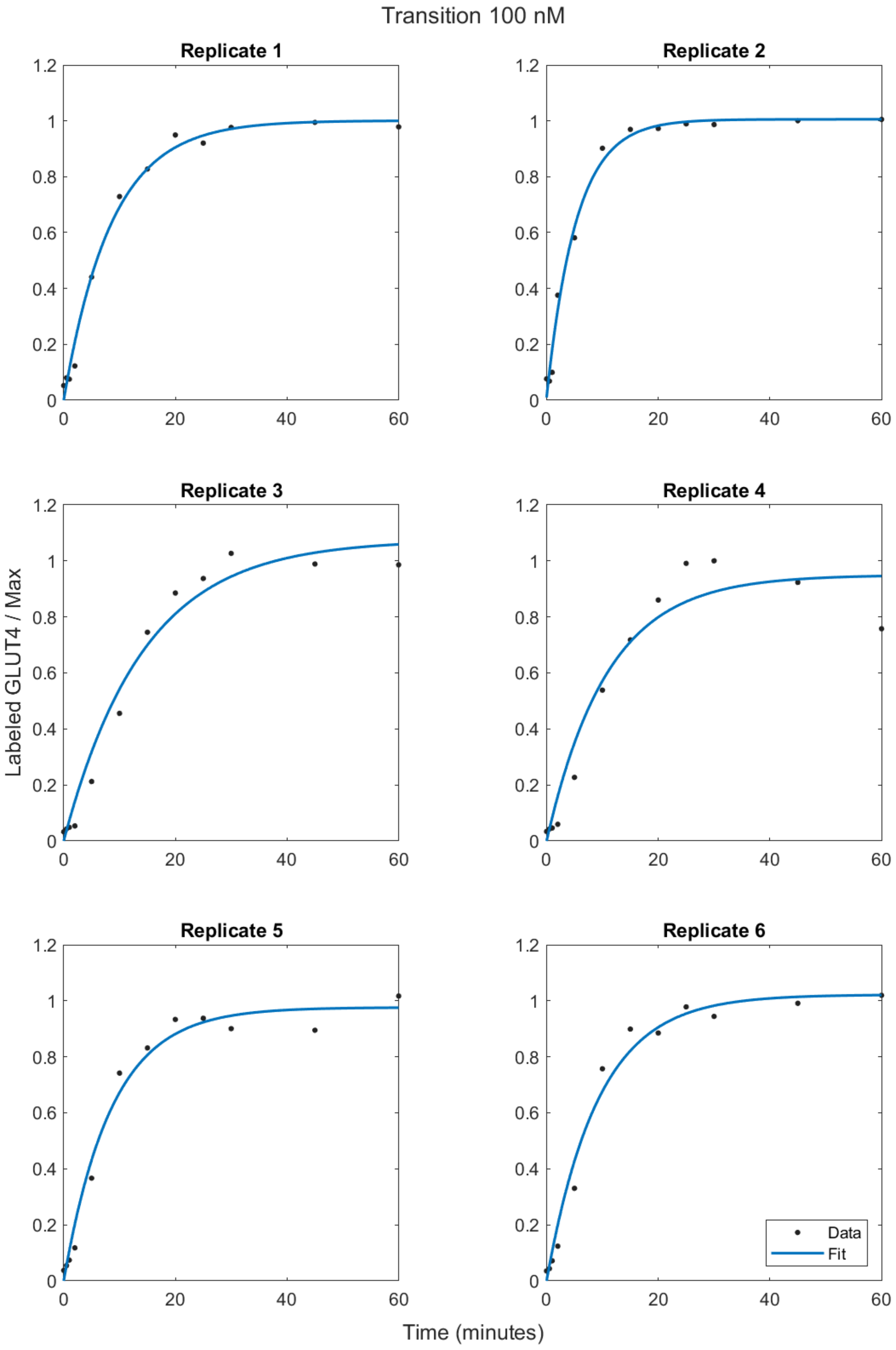


Figure S4d. Least-squares fits to Equation (4) for the Transition Assay as a function of the time of the data for each replicate in the data set as 100nM insulin was applied.

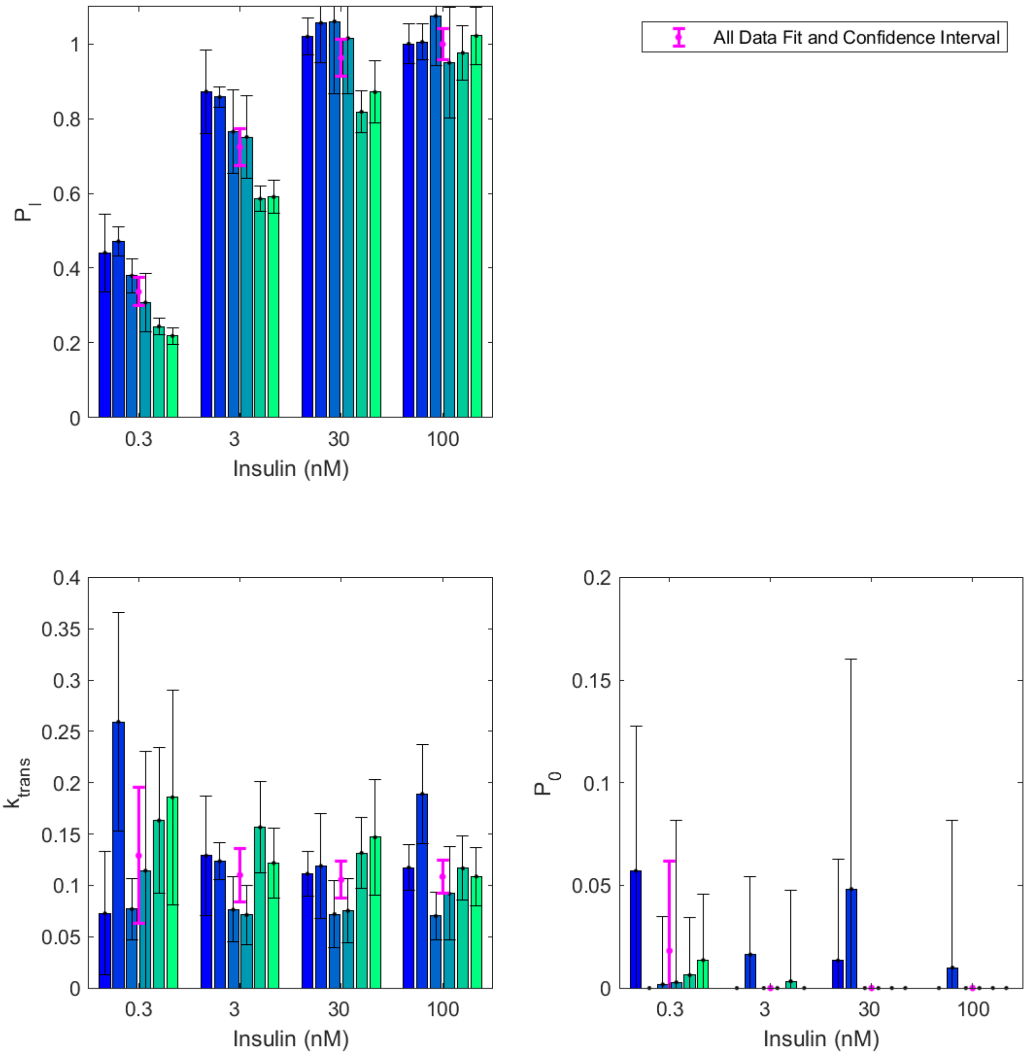


Figure S5. Parameter values of the least-squares fits to Equation (4) for the Transition Assay for each of the replicate data sets as a function of applied insulin concentration (categorical axis). The parameter values for the fits to all the data are shown in magneta. The error bars indicate the 95% confidence intervals for the parameter values.

Table S3. Parameter values for the least-squares fits to the transition data, Equation (4). The 95% confidence interval is reported in brackets. The adjusted $R^2$ value ($adjR^2$) accounts for the number of degrees of freedom in the model.

| nM | Replicate | adjR2 | $P$ | $P_0$ | $k_{trans}$ |
|---|---|---|---|---|---|
| 0.3 | 1 | 0.8730 | 0.4407 (0.3368,0.5447) | 0.0570 (-0.0137,0.1278) | 0.0727 (0.0125,0.1330) |
| 0.3 | 2 | 0.9377 | 0.4711 (0.4314,0.5109) | 0.0000 - | 0.2593 (0.1528,0.3658) |
| 0.3 | 3 | 0.9699 | 0.3792 (0.3333,0.4251) | 0.0017 (-0.0313,0.0348) | 0.0769 (0.0473,0.1065) |
| 0.3 | 4 | 0.8167 | 0.3074 (0.2304,0.3843) | 0.0028 (-0.0759,0.0815) | 0.1143 (-0.0017,0.2303) |
| 0.3 | 5 | 0.9630 | 0.2435 (0.2226,0.2644) | 0.0063 (-0.0218,0.0344) | 0.1632 (0.0925,0.2340) |
| 0.3 | 6 | 0.9389 | 0.2181 (0.1960,0.2401) | 0.0136 (-0.0187,0.0459) | 0.1858 (0.0811,0.2906) |
| 3 | 1 | 0.9486 | 0.8723 (0.7599,0.9848) | 0.0000 - | 0.1288 (0.0704,0.1873) |
| 3 | 2 | 0.9957 | 0.8579 (0.8301,0.8856) | 0.0163 (-0.0217,0.0543) | 0.1235 (0.1053,0.1417) |
| 3 | 3 | 0.9560 | 0.7652 (0.6536,0.8767) | 0.0000 - | 0.0763 (0.0446,0.1080) |
| 3 | 4 | 0.9586 | 0.7509 (0.6396,0.8622) | 0.0000 - | 0.0712 (0.0423,0.1001) |
| 3 | 5 | 0.9840 | 0.5855 (0.5513,0.6196) | 0.0033 (-0.0412,0.0478) | 0.1566 (0.1124,0.2007) |
| 3 | 6 | 0.9785 | 0.5903 (0.5457,0.6349) | 0.0000 - | 0.1218 (0.0879,0.1557) |
| 30 | 1 | 0.9921 | 1.0198 (0.9702,1.0693) | 0.0134 (-0.0362,0.0630) | 0.1113 (0.0896,0.1329) |
| 30 | 2 | 0.9620 | 1.0564 (0.9501,1.1626) | 0.0482 (-0.0640,0.1604) | 0.1189 (0.0673,0.1704) |
| 30 | 3 | 0.9569 | 1.0605 (0.8671,1.2539) | 0.0000 - | 0.0720 (0.0390,0.1049) |
| 30 | 4 | 0.9579 | 1.0155 (0.8663,1.1647) | 0.0000 - | 0.0752 (0.0440,0.1065) |
| 30 | 5 | 0.9810 | 0.8185 (0.7622,0.8748) | 0.0000 - | 0.1316 (0.0969,0.1663) |
| 30 | 6 | 0.9604 | 0.8711 (0.7879,0.9544) | 0.0000 - | 0.1469 (0.0902,0.2035) |
| 100 | 1 | 0.9900 | 1.0009 (0.9487,1.0531) | 0.0000 - | 0.1173 (0.0952,0.1394) |
| 100 | 2 | 0.9868 | 1.0055 (0.9570,1.0540) | 0.0099 (-0.0620,0.0818) | 0.1891 (0.1405,0.2376) |
| 100 | 3 | 0.9723 | 1.0749 (0.9423,1.2075) | 0.0000 - | 0.0702 (0.0467,0.0937) |
| 100 | 4 | 0.9399 | 0.9497 (0.8011,1.0983) | 0.0000 - | 0.0920 (0.0465,0.1375) |
| 100 | 5 | 0.9806 | 0.9763 (0.9037,1.0488) | 0.0000 - | 0.1166 (0.0854,0.1478) |
| 100 | 6 | 0.9816 | 1.0218 (0.9446,1.0990) | 0.0000 - | 0.1086 (0.0802,0.1370) |
| 0.3 | 1 | 0.8730 | 0.4407 (0.3368,0.5447) | 0.0570 (-0.0137,0.1278) | 13.7510 (2.3566,25.1454) |
| 0.3 | 2 | 0.9299 | 0.4711 (0.4272,0.5151) | 0.0000 (-0.1152,0.1152) | 3.8568 (1.3684,6.3452) |
| 0.3 | 3 | 0.9699 | 0.3791 (0.3332,0.4250) | 0.0018 (-0.0312,0.0348) | 12.9967 (7.9992,17.9943) |
| 0.3 | 4 | 0.8167 | 0.3073 (0.2304,0.3843) | 0.0028 (-0.0759,0.0816) | 8.7493 (-0.1340,17.6326) |
| 0.3 | 5 | 0.9630 | 0.2434 (0.2225,0.2643) | 0.0062 (-0.0219,0.0343) | 6.1127 (3.4622,8.7632) |
| 0.3 | 6 | 0.9389 | 0.2180 (0.1960,0.2401) | 0.0135 (-0.0188,0.0458) | 5.3677 (2.3422,8.3931) |
| 3 | 1 | 0.9486 | 0.8723 (0.7599,0.9848) | 0.0000 - | 7.7615 (4.2423,11.2807) |
| 3 | 2 | 0.9957 | 0.8579 (0.8301,0.8856) | 0.0163 (-0.0216,0.0543) | 8.0986 (6.9056,9.2917) |
| 3 | 3 | 0.9560 | 0.7652 (0.6536,0.8768) | 0.0000 - | 13.1047 (7.6641,18.5452) |
| 3 | 4 | 0.9586 | 0.7510 (0.6397,0.8623) | 0.0000 - | 14.0570 (8.3498,19.7642) |
| 3 | 5 | 0.9840 | 0.5854 (0.5513,0.6196) | 0.0033 (-0.0412,0.0478) | 6.3848 (4.5828,8.1868) |
| 3 | 6 | 0.9785 | 0.5903 (0.5457,0.6348) | 0.0000 - | 8.2102 (5.9262,10.4942) |
| 30 | 1 | 0.9921 | 1.0198 (0.9702,1.0693) | 0.0134 (-0.0362,0.0630) | 8.9889 (7.2402,10.7375) |
| 30 | 2 | 0.9620 | 1.0563 (0.9501,1.1626) | 0.0482 (-0.0641,0.1604) | 8.4091 (4.7632,12.0549) |
| 30 | 3 | 0.9569 | 1.0605 (0.8671,1.2539) | 0.0000 - | 13.8990 (7.5430,20.2550) |
| 30 | 4 | 0.9579 | 1.0154 (0.8663,1.1646) | 0.0000 - | 13.2885 (7.7739,18.8031) |
| 30 | 5 | 0.9810 | 0.8185 (0.7622,0.8749) | 0.0000 - | 7.5998 (5.5953,9.6043) |
| 30 | 6 | 0.9604 | 0.8711 (0.7879,0.9544) | 0.0000 - | 6.8084 (4.1819,9.4349) |

*Table S3 continued overleaf*

Table S3 continued.

Parameter values for the least-squares fits to the transition data, Equation (4).

| nM | Replicate | adjR2 | $P$ | | $P_0$ | | $k_{trans}$ | |
|---|---|---|---|---|---|---|---|---|
| 100 | 1 | 0.9900 | 1.0009 | (0.9487,1.0531) | 0.0000 | - | 8.5246 | (6.9192,10.1300) |
| 100 | 2 | 0.9868 | 1.0055 | (0.9570,1.0541) | 0.0100 | (-0.0619,0.0819) | 5.2893 | (3.9313,6.6473) |
| 100 | 3 | 0.9723 | 1.0749 | (0.9423,1.2075) | 0.0000 | - | 14.2464 | (9.4688,19.0239) |
| 100 | 4 | 0.9399 | 0.9495 | (0.8010,1.0979) | 0.0000 | - | 10.8575 | (5.4883,16.2267) |
| 100 | 5 | 0.9806 | 0.9763 | (0.9037,1.0488) | 0.0000 | - | 8.5757 | (6.2813,10.8701) |
| 100 | 6 | 0.9816 | 1.0218 | (0.9447,1.0990) | 0.0000 | - | 9.2082 | (6.7987,11.6177) |

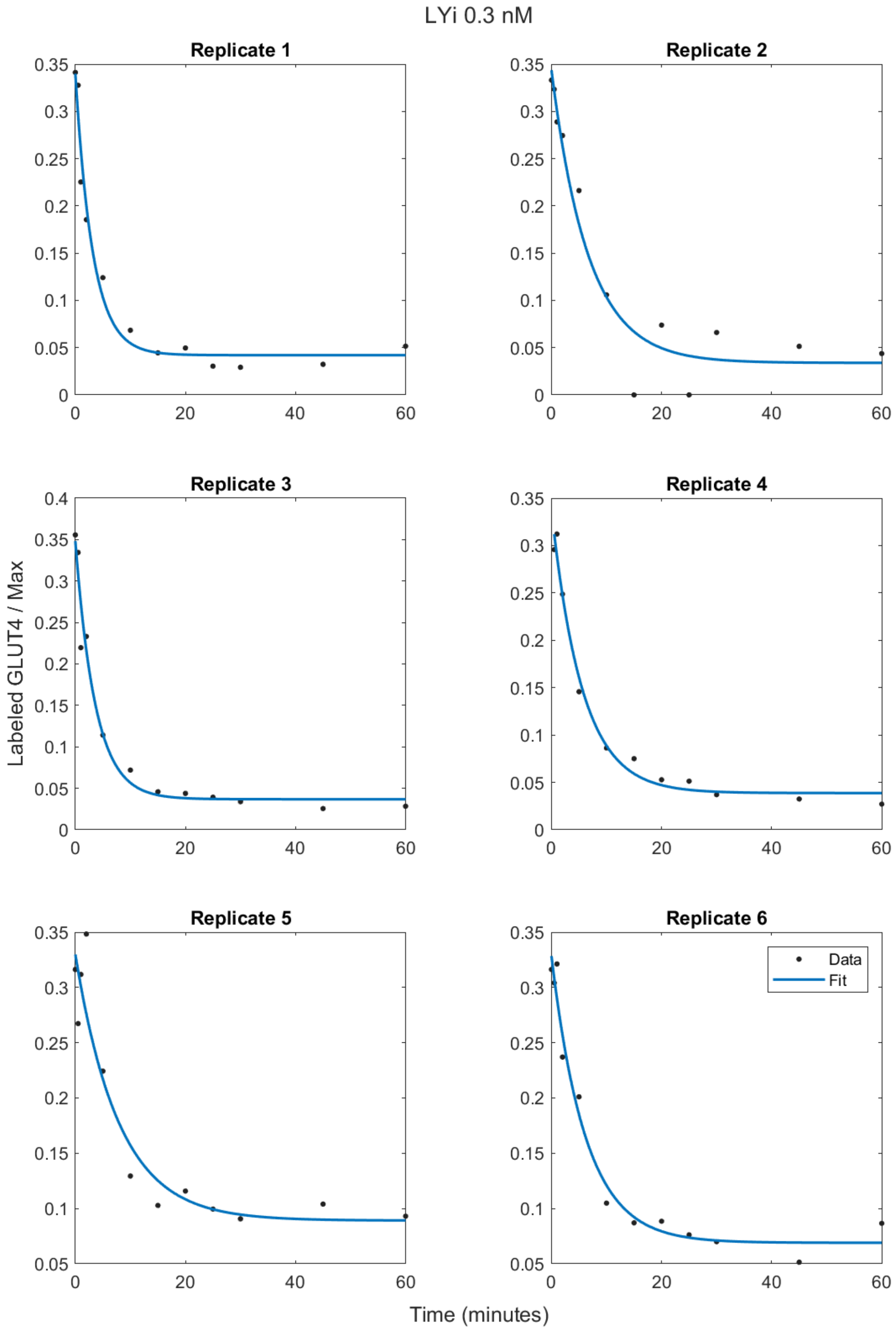


Figure S6a. Least-squares fits to Equation (6) for the Inhibition Assay as a function of the time of the data for each replicate in the data set as 0.3nM insulin was applied.

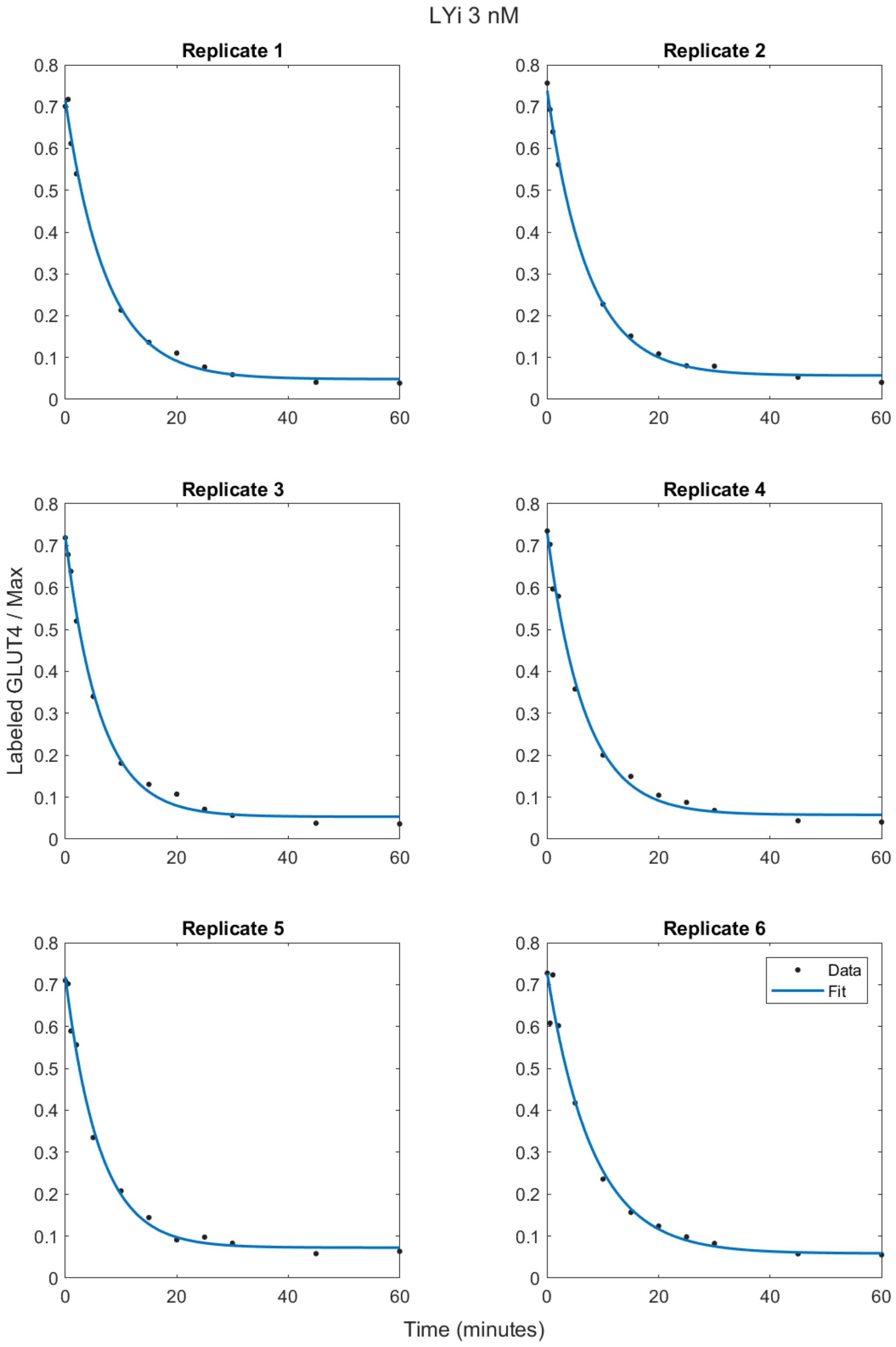


Figure S6b. Least-squares fits to Equation (6) for the Inhibition Assay as a function of the time of the data for each replicate in the data set as 3nM insulin was applied.

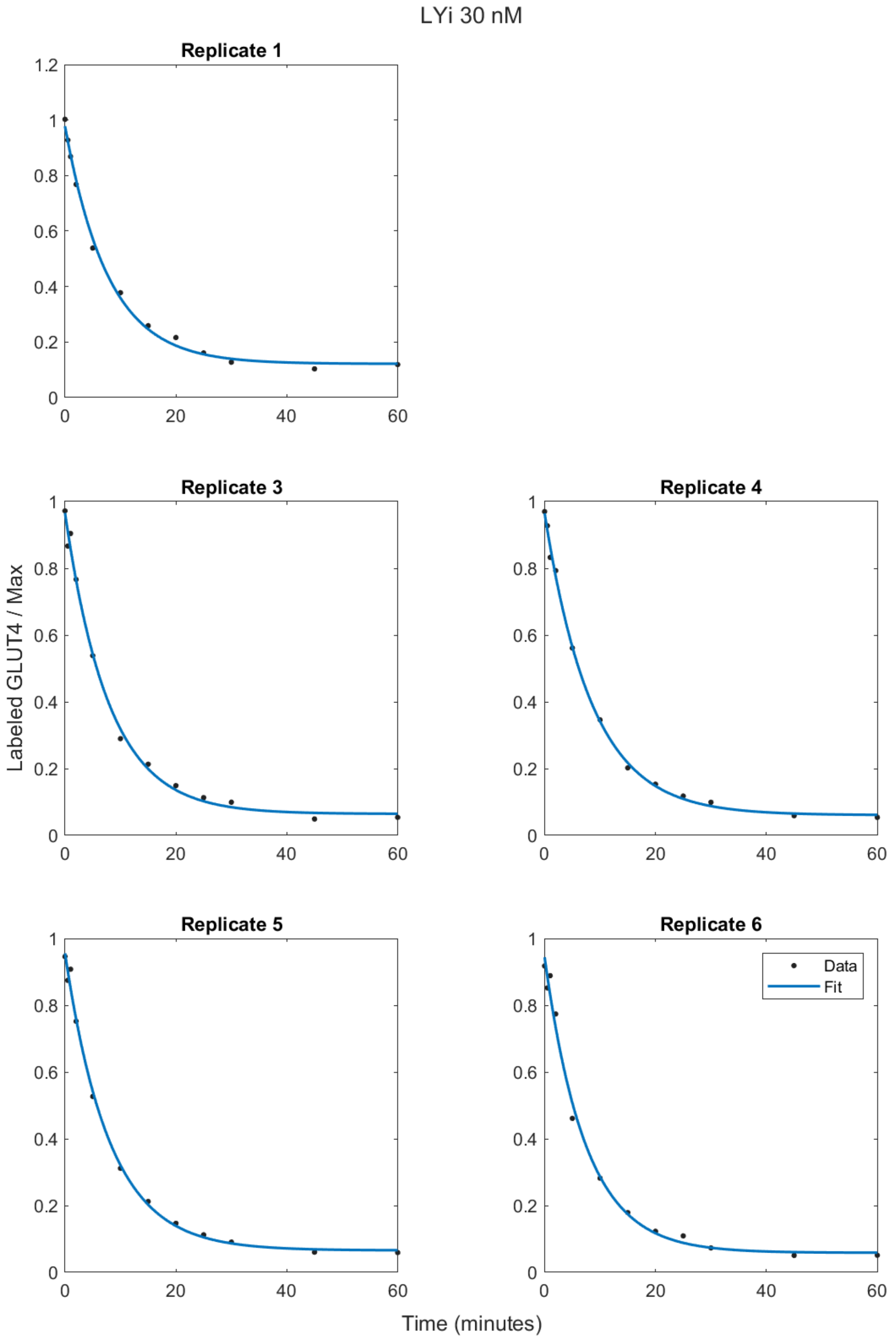


Figure S6c. Least-squares fits to Equation (6) for the Inhibition Assay as a function of the time of the data for each replicate in the data set as 30nM insulin was applied.

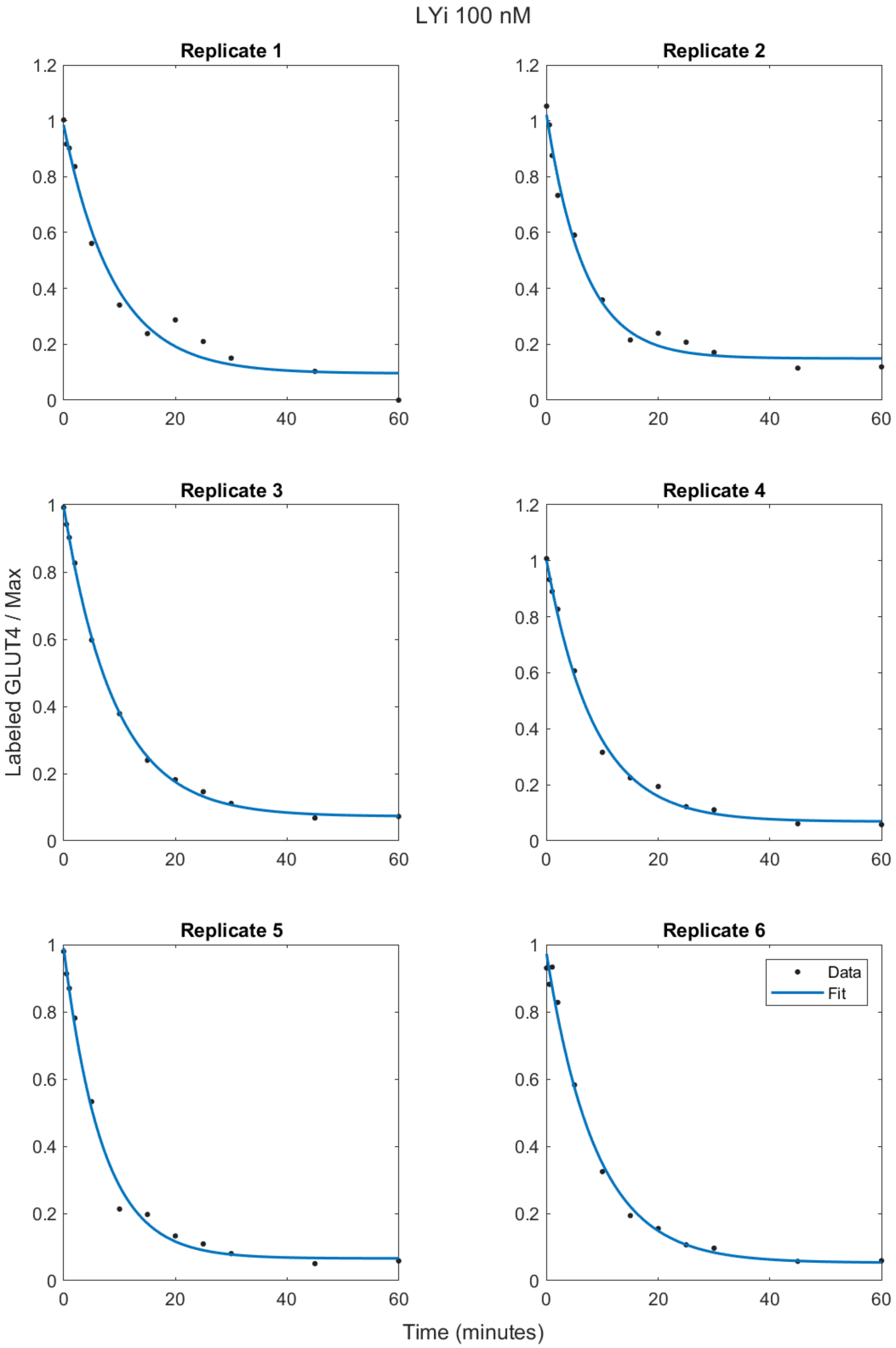


Figure S6d. Least-squares fits to Equation (6) for the Inhibition Assay as a function of the time of the data for each replicate in the data set as 100nM insulin was applied.

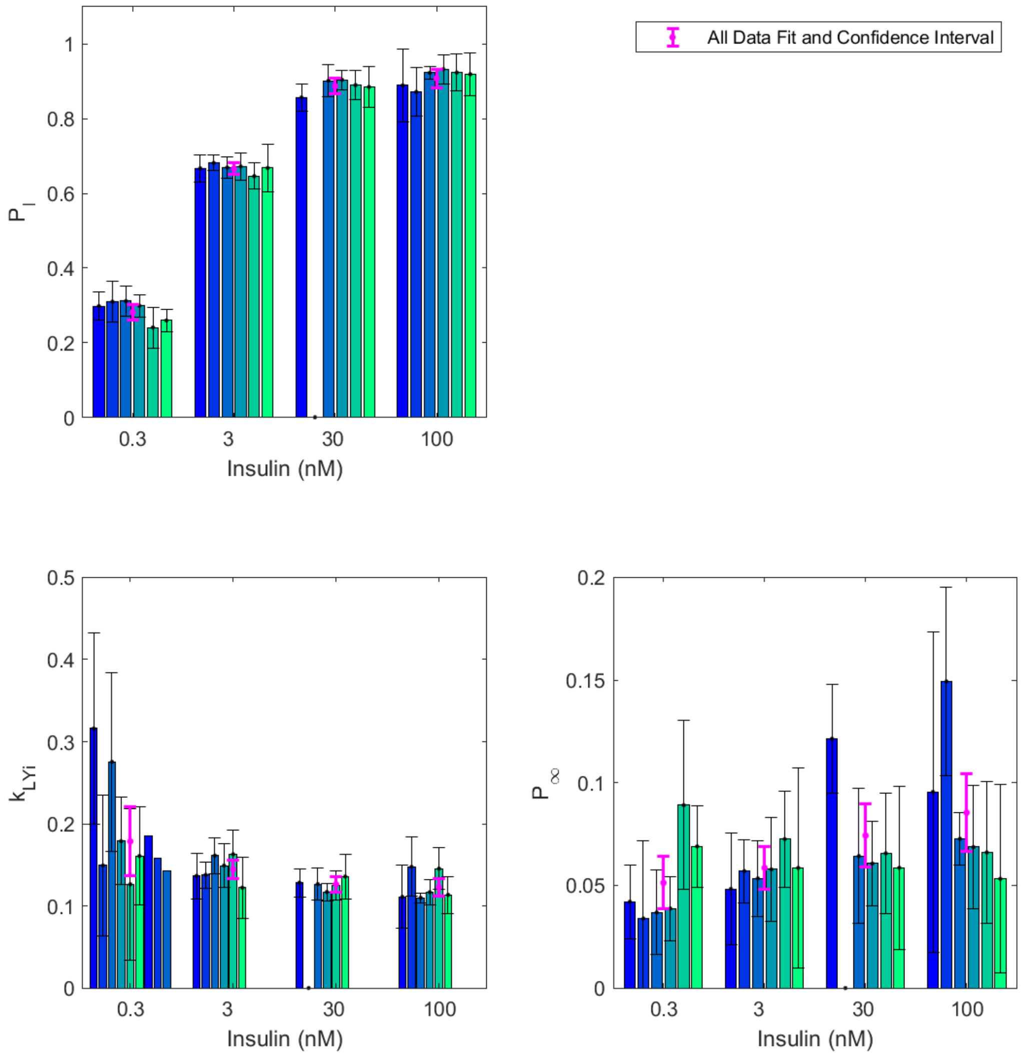


Figure S7. Parameter values of the least-squares fits to Equation (6) for the Inhibition Assay for each of the replicate data sets as a function of applied insulin concentration (categorical axis). The parameter values for the fits to all the data are shown in magneta. The error bars indicate the 95% confidence intervals for the parameter values.

Table S4. Parameter values for the least-squares fits to the inhibition data, Equation (6). The 95% confidence interval is reported in brackets. The adjusted $R^2$ value (adj$R^2$) accounts for the number of degrees of freedom in the model.

| nM | Replicate | adjR2 | $P$ | $P_\infty$ | $k_{LYi}$ |
|---|---|---|---|---|---|
| 0.3 | 1 | 0.9712 | 0.2979 (0.2607,0.3351) | 0.0420 (0.0242,0.0599) | 0.3160 (0.1990,0.4331) |
| 0.3 | 2 | 0.9360 | 0.3097 (0.2548,0.3646) | 0.0339 (-0.0041,0.0719) | 0.1493 (0.0634,0.2351) |
| 0.3 | 3 | 0.9680 | 0.3114 (0.2709,0.3519) | 0.0368 (0.0162,0.0574) | 0.2753 (0.1661,0.3845) |
| 0.3 | 4 | 0.9833 | 0.2989 (0.2686,0.3292) | 0.0387 (0.0230,0.0544) | 0.1789 (0.1257,0.2320) |
| 0.3 | 5 | 0.8992 | 0.2408 (0.1861,0.2954) | 0.0891 (0.0480,0.1302) | 0.1264 (0.0344,0.2184) |
| 0.3 | 6 | 0.9725 | 0.2594 (0.2296,0.2891) | 0.0690 (0.0492,0.0888) | 0.1608 (0.1009,0.2207) |
| 3 | 1 | 0.9945 | 0.6670 (0.6306,0.7035) | 0.0483 (0.0213,0.0754) | 0.1365 (0.1088,0.1643) |
| 3 | 2 | 0.9982 | 0.6815 (0.6605,0.7024) | 0.0570 (0.0415,0.0724) | 0.1378 (0.1219,0.1536) |
| 3 | 3 | 0.9963 | 0.6693 (0.6414,0.6973) | 0.0534 (0.0348,0.0720) | 0.1613 (0.1394,0.1831) |
| 3 | 4 | 0.9936 | 0.6721 (0.6353,0.7089) | 0.0578 (0.0324,0.0833) | 0.1492 (0.1227,0.1757) |
| 3 | 5 | 0.9936 | 0.6461 (0.6106,0.6816) | 0.0725 (0.0491,0.0960) | 0.1629 (0.1339,0.1920) |
| 3 | 6 | 0.9810 | 0.6680 (0.6043,0.7317) | 0.0585 (0.0099,0.1072) | 0.1222 (0.0850,0.1595) |
| 30 | 1 | 0.9963 | 0.8568 (0.8212,0.8925) | 0.1214 (0.0948,0.1481) | 0.1283 (0.1112,0.1455) |
| 30 | 3 | 0.9950 | 0.9017 (0.8579,0.9456) | 0.0643 (0.0314,0.0972) | 0.1267 (0.1069,0.1464) |
| 30 | 4 | 0.9982 | 0.9043 (0.8781,0.9306) | 0.0607 (0.0402,0.0812) | 0.1170 (0.1062,0.1278) |
| 30 | 5 | 0.9959 | 0.8901 (0.8511,0.9291) | 0.0656 (0.0361,0.0951) | 0.1248 (0.1073,0.1423) |
| 30 | 6 | 0.9918 | 0.8852 (0.8303,0.9400) | 0.0586 (0.0188,0.0984) | 0.1358 (0.1086,0.1629) |
| 100 | 1 | 0.9754 | 0.8893 (0.7917,0.9870) | 0.0956 (0.0175,0.1736) | 0.1109 (0.0725,0.1493) |
| 100 | 2 | 0.9878 | 0.8717 (0.8059,0.9376) | 0.1494 (0.1035,0.1952) | 0.1477 (0.1115,0.1839) |
| 100 | 3 | 0.9994 | 0.9229 (0.9068,0.9390) | 0.0727 (0.0598,0.0856) | 0.1099 (0.1038,0.1159) |
| 100 | 4 | 0.9964 | 0.9327 (0.8939,0.9714) | 0.0687 (0.0384,0.0989) | 0.1167 (0.1013,0.1321) |
| 100 | 5 | 0.9940 | 0.9237 (0.8747,0.9728) | 0.0661 (0.0316,0.1005) | 0.1455 (0.1204,0.1705) |
| 100 | 6 | 0.9917 | 0.9192 (0.8613,0.9771) | 0.0532 (0.0074,0.0990) | 0.1135 (0.0909,0.1362) |
| 0.3 | 1 | 0.9712 | 0.2979 (0.2607,0.3351) | 0.0420 (0.0242,0.0599) | 3.1633 (1.9921,4.3345) |
| 0.3 | 2 | 0.9360 | 0.3097 (0.2548,0.3646) | 0.0338 (-0.0042,0.0718) | 6.7037 (2.8484,10.5589) |
| 0.3 | 3 | 0.9680 | 0.3114 (0.2709,0.3519) | 0.0368 (0.0162,0.0574) | 3.6306 (2.1908,5.0704) |
| 0.3 | 4 | 0.9833 | 0.2989 (0.2686,0.3292) | 0.0387 (0.0231,0.0544) | 5.5878 (3.9274,7.2482) |
| 0.3 | 5 | 0.8992 | 0.2408 (0.1861,0.2955) | 0.0890 (0.0478,0.1301) | 7.9409 (2.1607,13.7211) |
| 0.3 | 6 | 0.9725 | 0.2594 (0.2296,0.2891) | 0.0690 (0.0492,0.0888) | 6.2187 (3.9034,8.5340) |
| 3 | 1 | 0.9945 | 0.6670 (0.6306,0.7035) | 0.0483 (0.0213,0.0753) | 7.3249 (5.8341,8.8156) |
| 3 | 2 | 0.9982 | 0.6815 (0.6605,0.7025) | 0.0569 (0.0414,0.0724) | 7.2605 (6.4258,8.0951) |
| 3 | 3 | 0.9963 | 0.6693 (0.6414,0.6973) | 0.0534 (0.0348,0.0720) | 6.2000 (5.3604,7.0397) |
| 3 | 4 | 0.9936 | 0.6721 (0.6353,0.7089) | 0.0578 (0.0323,0.0833) | 6.7042 (5.5142,7.8943) |
| 3 | 5 | 0.9936 | 0.6461 (0.6106,0.6816) | 0.0725 (0.0491,0.0960) | 6.1376 (5.0435,7.2316) |
| 3 | 6 | 0.9810 | 0.6680 (0.6043,0.7317) | 0.0585 (0.0099,0.1072) | 8.1821 (5.6892,10.6750) |
| 30 | 1 | 0.9963 | 0.8568 (0.8212,0.8925) | 0.1214 (0.0948,0.1480) | 7.7951 (6.7528,8.8375) |
| 30 | 3 | 0.9950 | 0.9017 (0.8579,0.9456) | 0.0643 (0.0313,0.0972) | 7.8958 (6.6643,9.1273) |
| 30 | 4 | 0.9982 | 0.9043 (0.8781,0.9306) | 0.0607 (0.0402,0.0812) | 8.5499 (7.7609,9.3388) |
| 30 | 5 | 0.9959 | 0.8901 (0.8511,0.9291) | 0.0656 (0.0361,0.0951) | 8.0146 (6.8902,9.1391) |
| 30 | 6 | 0.9918 | 0.8852 (0.8303,0.9400) | 0.0586 (0.0188,0.0984) | 7.3663 (5.8928,8.8397) |

*Table S4 continued overleaf*

Table S4 continued.

Parameter values for the least-squares fits to the inhibition data, Equation (6).

| nM | Replicate | adjR2 | $P$ | $P_{\infty}$ | $k_{LYi}$ |
|---|---|---|---|---|---|
| 100 | 1 | 0.9754 | 0.8894 (0.7917,0.9871) | 0.0955 (0.0174,0.1735) | 9.0197 (5.8981,12.1414) |
| 100 | 2 | 0.9878 | 0.8717 (0.8059,0.9376) | 0.1493 (0.1035,0.1952) | 6.7740 (5.1140,8.4340) |
| 100 | 3 | 0.9994 | 0.9229 (0.9068,0.9390) | 0.0727 (0.0597,0.0856) | 9.1032 (8.6033,9.6031) |
| 100 | 4 | 0.9964 | 0.9327 (0.8939,0.9714) | 0.0687 (0.0384,0.0989) | 8.5660 (7.4356,9.6965) |
| 100 | 5 | 0.9940 | 0.9237 (0.8747,0.9728) | 0.0661 (0.0316,0.1005) | 6.8738 (5.6896,8.0580) |
| 100 | 6 | 0.9917 | 0.9192 (0.8613,0.9771) | 0.0532 (0.0074,0.0990) | 8.8066 (7.0516,10.5616) |
